# Development of Non-Spatial Grid-Like Neural Codes Predicts Inference and Intelligence

**DOI:** 10.1101/2024.11.20.624569

**Authors:** Yukun Qu, Jianxin Ou, Luoyao Pang, Shuqi Wu, Yuejia Luo, Tim Behrens, Yunzhe Liu

**Affiliations:** State Key Laboratory of Cognitive Neuroscience and Learning, IDG/McGovern Institute for Brain Research, Beijing Normal University, Beijing, China; Chinese Institute for Brain Research, Beijing, China; School of Psychology, Center for Brain Disorders and Cognitive Science, Shenzhen University, Shenzhen, China; Wellcome Centre for Human Neuroimaging, UCL, London, UK; Wellcome Centre for Integrative Neuroimaging, University of Oxford, Oxford, UK; Sainsbury Wellcome Centre for Neural Circuits and Behaviour, UCL, London, UK

**Author notes:** Correspondence (Y.L.).

**Keywords:** cognitive map, development, inference, grid-like code, entorhinal cortex (EC), medial prefrontal cortex (mPFC), schema

## Abstract

Piaget’s theory posits that children develop structured knowledge schemas for inferring and assimilating new information, yet the underlying neural mechanisms remain unclear. In 203 participants aged 8–25 years, we investigated how maturation of a two-dimensional (2D) knowledge map underpins inferential reasoning and knowledge assimilation. Grid-cell-like codes in the entorhinal cortex (EC) strengthened with age, reflecting schema representations in non-spatial conceptual spaces, and were directly linked to improved inferential reasoning. These grid-like codes also supported the medial prefrontal cortex (mPFC) in encoding distance relationships between objects on the 2D map. As participants assimilated new information, they integrated it into existing grid patterns in the EC. Moreover, the maturation of these neural codes predicted real-world intelligence measures, particularly in reasoning abilities. Our findings demonstrate that the development of non-spatial grid-like neural codes offers a mechanistic account of cognitive development, bridging psychological theory with a fundamental cellular representation of cognitive map.

## INTRODUCTION

As we grow older, our understanding of the world deepens. Initially, we might remember facts as isolated pieces of information. As we mature, we begin to see not just the concepts themselves, but also how they connect to form a broader knowledge map. This emerging structure helps us infer relationships between concepts, even those not directly observed, ultimately allowing us to assimilate new concepts within the same framework. This structured knowledge, referred to as a cognitive map^1^ or schema^2,3^, undergoes ‘intellectual evolution’ as we develop from childhood to adulthood^4–7^. This represents a significant underpinning of human cognition^8^. However, the neural mechanism governing its development, remains largely unexplored.

There are two ways to develop structured knowledge^9^. The first involves creating links between concepts, allowing us to determine the distance between disjoint concepts when inferring their relationship. This ‘distance code’ representation, akin to a specific map, can be developed from experiences, such as forming a successor representation^10–12^. Alternatively, one can generalize a common abstract structure from different experiences, resembling a schema^13,14^. In two-dimensional (2D) environments, schema representation is supported by grid cells in the entorhinal cortex (EC). Grid cells fire in hexagonal patterns and generalize the grid pattern across environments^15,16^. This contrasts with the representation by hippocampal place cells, which encode a specific map and undergo remapping in different environments^17^. This applies to both physical space^18^ and arbitrary conceptual spaces^19,20^. In cognitive development, for instance, Piaget ^4^ posited that children begin to infer transitive rankings around the age of 7 or 8 years. A key question is whether this inferential reasoning ability arises from constructing a specific map representation or from forming a more general concept—namely, a schema.

These two representations are not mutually exclusive. Having a schema, for instance, allows for quicker construction of the current map and enables immediate inference without prior exposure^8^. We are interested in how such neural representations and their consequent inference ability evolves during development. We use a distance code to investigate map-like representations and a grid-like code for schema representation. If individuals construct their current map based on the schema, we expect to see a significant influence of the grid-like code on the distance code.

Once a schema is established, the process of assimilating new knowledge changes. Rather than creating a new map from scratch, one can incorporate local adjustments into the existing framework. This assimilation mechanism begins in childhood and persists throughout life, facilitating the ongoing development of cognition^21^. Animal studies shows that the medial prefrontal cortex (mPFC) is crucial in linking new cues to known environments, enabling rapid assimilation^22^. In such instances, the existing schema representation should remain stable. In the current study, we investigate how the assimilation of new knowledge develops with age, specifically after a conceptual schema is formed. This question is especially intriguing because the mPFC, which continues to mature into adulthood^23,24^, is thought to play a crucial role in the accumulation of knowledge^25,26^. We hypothesise that effective assimilation depends on how well novel information aligns with the existing schema, and that the mPFC involvement reflects an age-dependent process.

## RESULTS

### Age-related improvement in inferential reasoning and new knowledge assimilation

We recruited 231 participants aged 8–25 years in formal fMRI experiment, ensuring that each age year included at least ten subjects. Before the main experiment, participants memorised rank differences between 25 objects, each with two independent attributes (attack power, AP, and defence power, DP). Unbeknownst to them, these objects were arranged into a 2D map with five ranks per dimension. Participants learnt only the rank differences between adjacent objects in each dimension (see STAR Methods). During fMRI scanning, they had to infer relationships across both dimensions, resembling navigation on a 2D knowledge map (**Figure 1A**).

**Figure 1.**
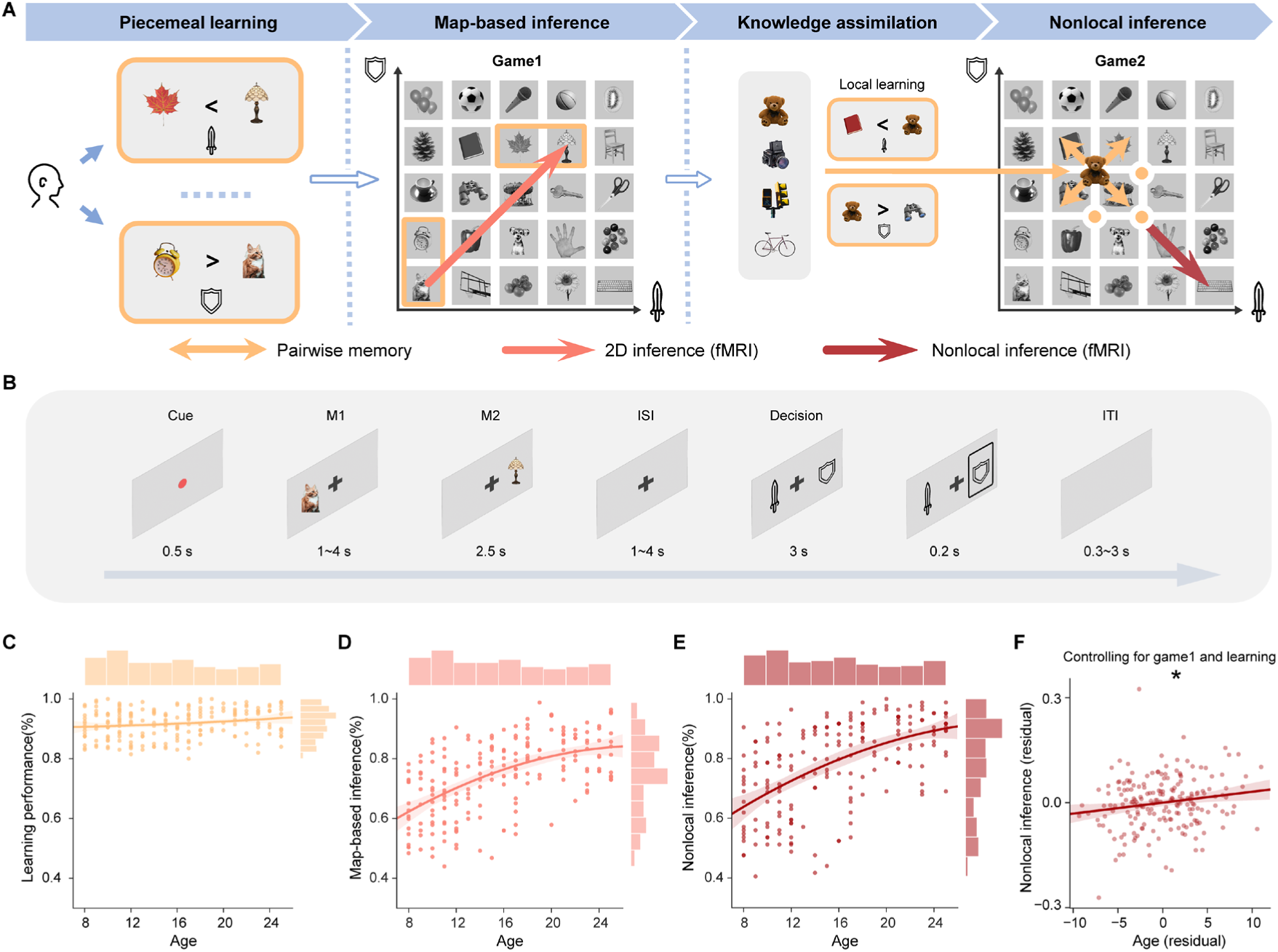
Experimental design and the age-related increase in inference ability on a 2D knowledge map. **(A)** The map comprises two independent dimensions—attack power (AP) and defence power (DP)—and contains 25 objects (or ‘monsters’ in cover story). During the memory training phase, participants learnt one-dimensional neighbouring relationships (yellow rectangle) without knowing the latent 2D structure. Only those reaching ≥80% accuracy proceeded to the map-based inference task (Game 1), conducted with fMRI. Game 1 required participants to consider both dimensions and infer relationships between novel object pairs (red arrows) across 252 trials covering all trajectory directions. Afterwards, four new objects were placed at the map centre (e.g., the “bear” and three orange points). Participants first learnt their relationships with local neighbours (yellow arrows), then made nonlocal inferences between these new objects and distant existing ones (dark red arrow) in Game 2, testing their ability to assimilate new knowledge. **(B)** In each inference trial, Monster1 (M1) and Monster2 (M2) appear sequentially, followed by a fighting rule that specifies whether AP or DP is compared to determine the winner. Changing rules encourage participants to separate mental navigation from the decision-making process (see also **Figure S4** for validation results). **(C-E)** Developmental changes of behavioural performance, separately for piecemeal learning (**C**), map-based inference (**D**), and non-local inference with new knowledge (**E**). **(F)** Even after controlling for learning performance and Game 1 inference, the correlation between age and Game 2 ability remained significant. Each circle denotes one participant; error bars indicate s.e.m. *: *p* < 0.05.

In the training phase, participants engaged in “piecemeal learning”. They learnt one-rank differences between objects in each dimension (see also **Figure S1**). They were informed that these one-rank differences were equidistant, but were not told about the underlying 2D structure. This training could span multiple days. Only those consistently reaching at least 80% accuracy proceeded to the fMRI experiment, ensuring sufficient exposure to potentially form a mental 2D map. Before scanning, participants reviewed all learnt pairs and completed a memory probe test without feedback. Those exceeding 80% accuracy advanced to the fMRI session.

To help younger participants (under 12 years old) become familiar with the MRI environment, they underwent a mock MRI session (see STAR Methods). All participants received clear task instructions and confirmed their understanding before scanning. We also verified through post-scan debriefings that everyone, including the youngest participants, fully comprehended the tasks, ensuring they performed as intended.

During the fMRI experiment, participants encountered novel inference questions requiring knowledge from both dimensions, termed “map-based inference” or “Game 1.” In each trial, they viewed two objects sequentially, then, after a brief delay, were shown a decision rule indicating whether to compare attack power against defence power to identify the winner (**Figure 1B**; see also STAR Methods). If participants successfully organised their existing knowledge into a five-by-five 2D map, each inference trial would resemble mental navigation within that map.

After excluding participants who failed memory tests or exhibited excessive head movement, 203 remained for the final analysis (see also **Table S1**). Following the map-based inference (“Game 1”) phase, four new objects were introduced at the map centre. Participants learnt their rank differences relative to immediate neighbours in a “local learning” phase. They were told that existing object relationships would remain unchanged. Those who successfully assimilated this new information into the existing map could make “nonlocal” inferences between the new objects and distant existing ones, termed “Game 2” or “nonlocal inference.” Of the 203 participants, 193 completed Game 2 with usable fMRI data.

To validate our task design for capturing age-related development in structured knowledge, we analysed behavioural performance in both Game 1 and Game 2. We observed a positive correlation between age and memory performance (*r* = 0.19, *p* = 0.01, *N* = 203; **Figure 1C**). More importantly, there was a strong age effect on map-based inference performance in Game 1 (*r* = 0.56, *p* < 0.001; **Figure 1D**), which remained significant after adjusting for memory differences (*r* = 0.54, *p* < 0.001). These results indicate that age influences inferential reasoning over and above memory capacity.

A similar pattern emerged for nonlocal inference in Game 2 (*r* = 0.56, *p* < 0.001; adjusted for memory: *r* = 0.54, *p* < 0.001, *N* = 193; **Figure 1E**). A linear regression model revealed a significant effect of age on nonlocal inference performance (*t*(188) = 2.02, *p* = 0.04, Cohen’s *d* = 0.10), whereas memory performance during learning did not (training memory: *t*(188) = –0.35, *p* = 0.73, Cohen’s *d* = –0.01; new knowledge memory: *t*(188) = 1.47, *p* = 0.14, Cohen’s *d* = 0.07). Notably, the age effect on Game 2 performance persisted after controlling for Game 1 performance (*r* = 0.17, *p* = 0.02; **Figure 1F**), suggesting a distinct developmental effect for assimilating new information beyond established map knowledge.

To ensure that the age-related effect was not simply due to younger participants struggling with the task, we confined our analysis to those under 12 years old and found they performed significantly above chance on all tasks (map-based inference: *t* = 12.46, *p* < 0.001; nonlocal inference: *t* = 11.01, *p* < 0.001). We then tested whether differences in memory capacity could explain the age-related improvement in inference by excluding trials where objects were incorrectly identified in the memory test. The age effect on map-based inference remained significant (*r* = 0.51, *p* < 0.001). Finally, we performed extensive analyses to rule out possible age-related confounding factors such as sampling biases, forgetting, and participant fatigue, none of which accounted for the observed improvements (see details in **Data S1**). Altogether, these findings indicate that inferential reasoning and knowledge assimilation develop with age, above and beyond memory capacity, highlighting a selective developmental role in forming knowledge structures and integrating new information.

### Development of grid-like code in the EC underlies inferential reasoning

We aimed to uncover the neural basis underlying the development of structured knowledge and its resulting flexible behaviour. First, we asked whether the grid-like code^15,18^—a cellular mechanism for representing schemas in 2D spaces—also applies to knowledge maps in conceptual tasks, and how it evolves with age to support inferential reasoning abilities.

To this end, we employed three established fMRI methods for detecting grid-like code. These are: a) hexagonal modulation analysis^20,27^; b) hexagonal consistency analysis^18–20^; and c) representation similarity analysis (RSA) of angular difference between inferred trajectories^28^. All these approaches detect alignment between neural signals and grid angle, specifically examining six-fold or hexagonal symmetry (Figure 2A).

**Figure 2.**
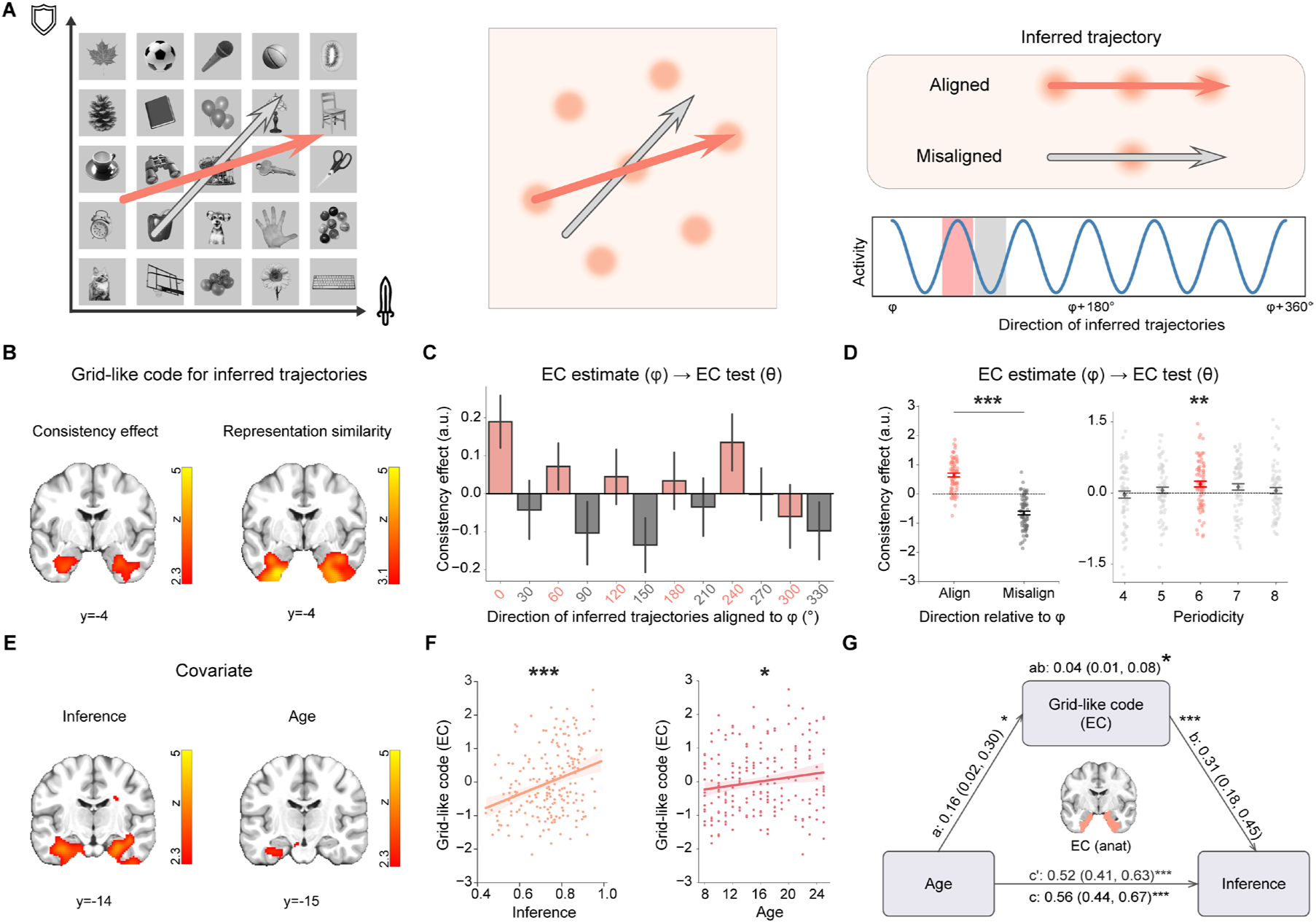
Grid-like code in the EC supports age-related improvements in 2D map-based inference. **(A)** Illustration of the hexagonal grid-code pattern. Inferred trajectories traverse both the conceptual knowledge map and the firing fields of grid cells. When a trajectory’s direction (θ) aligns with the mean orientation (φ) of neuronal ensembles, it intersects more grid fields, yielding stronger neural activity than misaligned trajectories. **(B)** Grid-like codes for inferred trajectories were identified in the EC via both univariate parametric analysis (left) and multivariate representation similarity analysis (right), observed in participants with high inference accuracy (accuracy > 0.8; N = 75). **(C)** The EC exhibits a six-fold modulation when a trajectory’s direction (θ) aligns with the mean orientation (φ). Responses are shown separately for aligned (red) and misaligned (grey) trajectories. **(D)** Left: Aligned trajectories elicit significantly stronger neural activity than misaligned ones. Right: The six-fold modulation aligned to φ is specific to six-fold periodicity, with no effect at other periodicities. **(E)** Whole-brain covariate analysis incorporating age and inference performance as separate second-level covariates highlights positive effects in the EC (thresholded at p < 0.01, uncorrected, for visualisation). **(F)** ROI analysis of the grid-like code in the anatomically defined bilateral EC shows positive correlations with inference and age. **(G)** Mediation analysis indicates that stronger EC grid-like codes mediate the age-related increase in inferential reasoning. Statistical inferences used bootstrapping with 10,000 iterations; paths are labelled with coefficients and 95% confidence intervals. *: *p* < 0.05. **: *p* < 0.01. ***: *p* < 0.001.

Across all three methods, we consistently identified significant grid-like code in the EC (Figure 2B; whole-brain cluster-based family-wise error (FWE) correction at *p* < 0.05; hexagonal modulation effect: peak *z* = 5.48, MNI coordinate [−20, 1, −34], consistency effect: peak *z* = 4.69, MNI coordinate [−28, 3, −38], left panel; and representation similarity: peak *z* = 4.93, MNI coordinate [−28, −2, −42], right panel, also see **Figure S2** and **Data S2**). To visualise the hexagonal activation pattern, we grouped all inferred trajectories into 12 bins based on their angle directions relative to the grid angle (Figure 2C). Neural activity in the EC anatomical ROI ^29^ was significantly higher for aligned than for misaligned trajectories (paired-*t* test, *t*(74) = 10.15, *p* < 0.001, Cohen’s *d* = 2.34; Figure 2D). Notably, this consistency effect was significant only for six-fold symmetry (one-sample *t* test, *t*(74) = 3.12, *p* = 0.003, Cohen’s *d* = 0.36), not for 4, 5, 7, or 8-fold periodicities (Figure 2D), and these effects persisted also after controlling for distance differences between angular bins (*t*(74) = 2.58, *p* = 0.01, Cohen’s *d* = 0.3; see also **Figure S3**).

After identifying the grid-like code on the 2D knowledge map across subjects, we examined its relationship with age and cognitive map-based inference. We measured the grid-like code using RSA, which does not require estimating the ROI-specific mean orientation^28^. A whole-brain covariate analysis revealed an increase in EC grid-like code with both inference performance and age (age: peak *z* = 2.70, MNI coordinate [−32, −16, −28]; inference: peak *z* = 4.72, MNI coordinate [27, −10, −28]; Figure 2E). Further ROI analysis confirmed these effects (inference: *r* = 0.31, *p* < 0.001; age: *r* = 0.16, *p* = 0.02, Figure 2F).

To verify the robustness of the observed age-related increase in EC grid-like codes, we performed multiple control analyses. First, we tested whether the correlation with age could be attributed to general changes in the EC’s signal-to-noise ratio (SNR). No significant relationship was found (Game 1: *r* = −0.09, *p* = 0.19; Game2: *r* = −0.06, *p* = 0.38; see **Data S3**), indicating that the findings are not due to SNR differences across ages. Next, to rule out systematic differences between adults and children, we repeated the analysis with participants under 18 years old. The EC grid-like code still correlated significantly with both inferential performance (*r* = 0.35, *p* < 0.001) and age (*r* = 0.18, *p* = 0.04). Moreover, when examining only the youngest children, we discovered that the relationship between grid-like code and improved inference performance was strongest in children aged 8 to 12 years (*r* = 0.32, *p* = 0.004).

These findings suggest that developing grid-like codes in the EC plays a crucial role in supporting inferential abilities. A mediation analysis directly tested this idea, showing that the age-related increase in inferential performance was mediated by the EC grid-like code (mediation effect: *β*_ab_ = 0.04, *p* = 0.03, CI [0.01, 0.08]; Figure 2G). This mediation effect remained significant even after controlling for socioeconomic status (*β*ab = 0.06, *p* = 0.02, CI [0.01, 015], see also **Data S1**).

Together, these findings demonstrate that grid-like code in the EC represents the 2D map structure. As individuals mature, these grid-like codes become stronger, underpinning the observed improvements in inferential reasoning.

### Development of EC grid-like code supports map-like representation in the mPFC

Having identified the schema representation (grid-like code) in the EC, we next investigated the map-like representation (distance code) in the mPFC and its relationship to the schema. The distance code represents the Euclidean distances between objects in the map and has been previously identified in the mPFC^19,30,31^. During the mental navigation phase (when the second object, M2, appears), we estimated the Euclidean distance between inferred object pairs (Figure 3A). Crucially, if the current map representation is built from the schema, we expect the grid-like code to influence this distance code.

**Figure 3.**
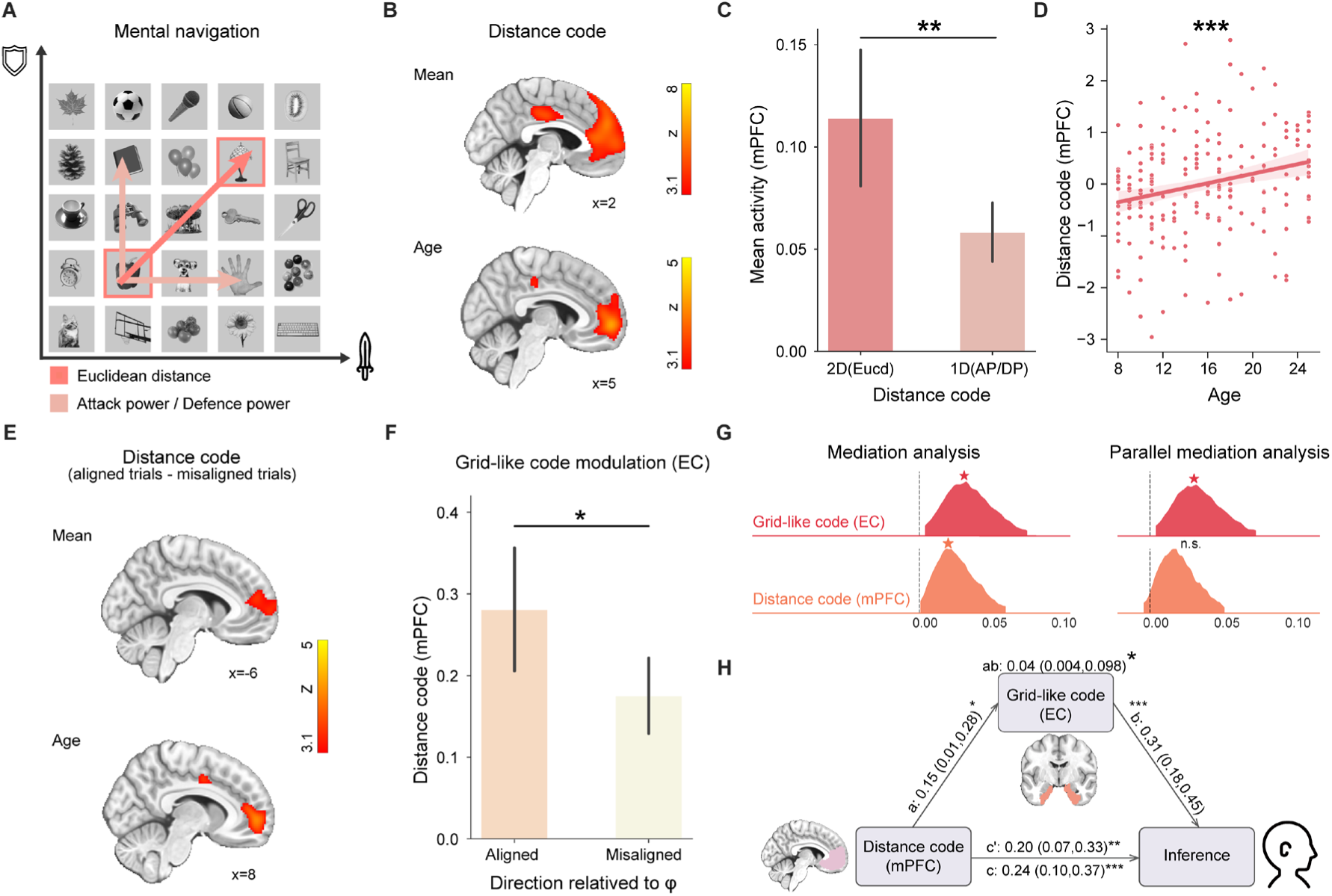
Development of the grid-like code in the EC supports the map-based distance code in the mPFC. **(A)** An example trajectory (e.g., paprika → lamp) traverses the 2D knowledge map during mental navigation (when M2 appears). We compute the Euclidean distance (red arrow) to capture map-like relationships across both dimensions, beyond 1D differences (light pink arrow). **(B)** Whole-brain analysis reveals significant mPFC activation encoding Euclidean distance (top) and an age-related increase in the mPFC distance code (bottom). **(C)** ROI analysis in the mPFC shows stronger representation of the 2D Euclidean distance than a 1D hierarchy (see also **Figure S4**). **(D)** The mPFC distance code positively correlates with age. **(E)** Whole-brain modulation of the EC grid-like code on distance coding identifies significant mPFC activation for aligned versus misaligned trials (top), with an age-related increase (bottom). Alignment is defined when the inferred trajectory’s direction (θ) deviates by less than 15° from the EC mean orientation (φ). Deviations > 15° are considered misaligned. **(F)** ROI analysis confirms stronger distance coding for aligned trials in the mPFC than for misaligned ones. **(G)** Mediation analysis of task-related neural measures on the relationship between age and behavioural performance. The dotted line represents zero. Mediation effects was significant when bootstrap confidence intervals exclude zero. **(H)** Further analysis indicates that the EC grid-like code mediates the mPFC distance code on inference performance. Statistical inferences used 10,000 bootstrapped iterations, with paths labelled by coefficients and confidence intervals. All brain maps were whole-brain FWE corrected at the cluster level (*p* < 0.05) with a cluster-forming voxel threshold of *p* < 0.001. Each circle is data from individual subject. Error bars denote s.e.m. *: *p* < 0.05. **: *p* < 0.01. ***: *p* < 0.001. n.s. non-significant.

Across all subjects, we observed significant mPFC involvement in encoding the distance code (whole-brain cluster-based FWE correction at *p* < 0.05, peak *z* = 6.23, MNI coordinate [7, 47, 23]; Figure 3B, upper panel), as well as activation in the hippocampus, LOFC and PCC (see also **Data S4**). A whole-brain covariate analysis also revealed a positive correlation between the distance code and age in the mPFC (peak *z* = 4.30, MNI coordinate [5, 57, −2]; Figure 3B, bottom panel). At the ROI level, we defined the mPFC independently using a NeuroVault mask (see STAR Methods). Notably, the mPFC encoded more of the 2D Euclidean distance than a one-dimensional (1D) hierarchy (paired t-test, *t*(202) = 2.69, *p* = 0.008, Cohen’s *d* = 0.15; Figure 3C), consistent with previous findings indicating that the mPFC integrates 2D information rather than representing each dimension separately^27^. The mPFC distance code also showed positive correlations with both age (*r* = 0.24, *p* < 0.001, Figure 3D) and inference performance (*r* = 0.23, *p* < 0.001).

Next, we asked whether subjects constructed the map-like representation based on the schema (see **Data S5** for a schematic distinction between schema and map). We classified all trials as ‘aligned’ or ‘misaligned’ according to their angle relative to the mean orientation estimated from the EC. At the whole-brain level, we found a stronger distance code for aligned trials than for misaligned trials, with only the mPFC surviving whole-brain correction (peak *z* = 3.57, MNI coordinate [0, 33, 11]; Figure 3E). ROI analysis further supported these results (All subjects: *t*(202) = 1.95, *p* = 0.03, Cohen’s *d* = 0.15; high performance subjects: *t*(74) = 2.72, *p* = 0.004, Cohen’s *d* = 0.22, paired *t* test; Figure 3F; see also **Figure S3**). These findings indicate a consistent relationship between the EC grid-like code and the mPFC distance code, suggesting that the map-like representation is derived from the schema.

Finally, we examined how the mPFC contributes to the age-related increase in cognitive map-based inference, particularly in relation to the EC grid-like code. A mediation analysis showed that the distance code significantly mediated the age-related improvement in inference (*β*_ab_ = 0.03, *p* = 0.04, CI [0.002, 0.067], Figure 3G, left panel). However, when considering both the EC grid-like code and the mPFC distance code together in a parallel mediation analysis^32^, only the EC effect remained significant (EC, *β*_ab_ = 0.04, *p* = 0.03, CI [0.006, 0.082]; mPFC, *β*_ab_ = 0.02, *p* = 0.12, CI [−0.002, 0.057]; Figure 3G, right panel). Further tests confirmed a significant mediation effect by the EC grid-like code on the mPFC influence on inference performance (*β*_ab_ = 0.04, *p* = 0.04, CI [0.004, 0.098]; Figure 3H).

All these effects were specific to the mental navigation when mPFC representing the distance code, rather than to the decision time (**Figure S4**; see also **Data S6-8**). Together, these results suggest that subjects constructed their current map representation using the EC-based schema, and that the development of this schema supports the map-like representation in the mPFC.

### Consistency in the grid-like code facilitates new knowledge assimilation

After completing the main inference task, participants were introduced to four new objects placed at the centre of the original map and learnt their rank relationships with nearby items. They then made novel inferences between these new objects and distant existing ones— referred to as nonlocal inference, or Game 2 (see STAR Methods). The overall map structure, and thus the schema, remained constant across both inference tasks. If participants effectively integrated these local changes into the existing schema (Figure 4A; **Data S9**), they should maintain strong nonlocal inference performance. To examine this, we tested whether the grid-like code persisted after introducing the new objects.

**Figure 4.**
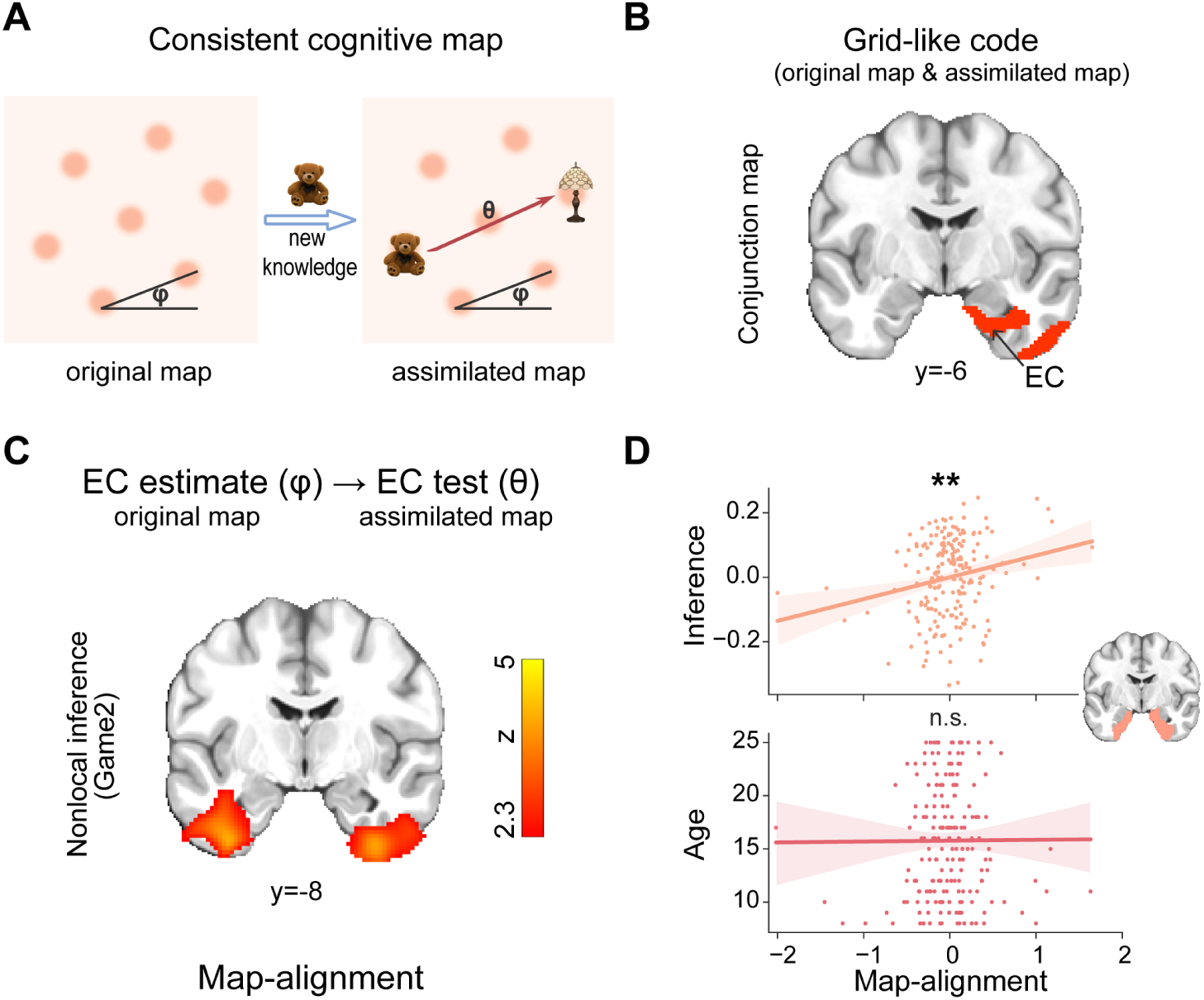
Consistency in the EC grid-like code facilitates nonlocal inference on a new map. **(A)** Illustration on map alignment. If the schema underlies knowledge assimilation, the original grid angles (φ) should remain consistent after new objects are introduced. Thus, trajectories (θ) on the new objects should align with the original grid angle. **(B)** A conjunction analysis of grid-like codes from the original map (Game 1) and the knowledge-assimilated map (Game 2) revealed a significant overlap in the right EC. **(C)** To test map alignment, we compared inferred trajectories on the new map (Game 2) to the EC mean grid angle from the original map (Game 1). We identified significant EC clusters predicting nonlocal inference performance (*p* < 0.05, cluster-based FWE; cluster-forming threshold *p* < 0.01), even after adjusting for age. **(D)** ROI analysis showed that map-alignment strength in the EC positively predicted nonlocal inference performance but was unrelated to age. Each circle is data from individual subject. **: *p* < 0.01. n.s. non-significant.

In Game2, we first replicated the mean brain activation pattern observed during inference in Game1 (**Figure S5**, see also **Data S10**). We then repeated the grid-code analysis, identifying significant grid-code in the EC (peak *z* = 3.75, MNI coordinate [19, −10, −32]), mPFC (peak *z* = 4.10, MNI coordinate [−4, 21, −6]), LOFC (peak *z* = 4.31, MNI coordinate [15, 21, −26]) and PCC (peak *z* = 3.9, MNI coordinate [1, −44, 27], also see **Figure S6** and **Data S11**), all surviving whole-brain correction (cluster-based FWE corrected at *p* < 0.05). These regions matched those that previously showed grid-like codes in Game 1. A conjunction analysis comparing grid-like codes between the original and knowledge-assimilated maps revealed a significant cluster in the right EC (Figure 4B), indicating that participants retained a consistent schema representation after assimilating new information.

We next assessed the alignment of the grid-like code across the two maps by deriving a grid angle from the EC in Game 1 and comparing it with inferred trajectories in Game 2. The EC displayed a significant map-alignment effect that positively correlated with nonlocal inference performance (whole-brain cluster-based FWE correction at *p* < 0.05, peak *z* = 4.32, MNI coordinate [29, 0, −42], Figure 4C), even after controlling for age. ROI analysis further showed that the map-alignment strength in EC predicted nonlocal inference performance (*r* = 0.21, *p* = 0.003) but was not related to age (*r* = 0.01, *p* = 0.94; Figure 4D). These findings suggest that preserving a consistent schema enables participants to integrate new knowledge and make novel inferences effectively.

### mPFC selectively supports age-related development of knowledge assimilation

After establishing a consistent schema representation across maps, we asked how the current map adapts to new information. Previous work ^22^ indicates that linking new information to an existing schema supports rapid integration, a process in which the mPFC is critical. We therefore expected the mPFC representation of the new map to be key for knowledge assimilation, particularly given that the human mPFC continues to mature relatively late^23,24^, implying a developmental trajectory for its role in knowledge assimilation.

In Game 2, we again observed a significant distance code in the mPFC (peak *z* = 7.13, MNI coordinate [−8, 55, 3], Figure 5A, left panel), along with activation in the hippocampus, LOFC, and PCC (**Data S12**). These findings survived whole-brain correction (cluster-based FWE corrected at *p* < 0.05). A conjunction analysis of the distance code in the original and new maps also showed a significant cluster in the mPFC (Figure 5A, right panel). Involvement of the mPFC during Game 2 was similarly noted in a whole-brain covariate analysis with both age (peak *z* = 6.41, MNI coordinate [−4, 39, −9]) and nonlocal inference performance (peak *z* = 7.69, MNI coordinate [3, 33, −12], Figure 5A, bottom panel). As in Game 1, the mPFC distance code in Game 2 was modulated by the EC grid-like code, showing stronger activation in aligned than misaligned trials (paired *t* test, *t*(192) = 4.14, *p* < 0.001, Cohen’s *d* = 0.23, see **Figure S6**).

**Figure 5.**
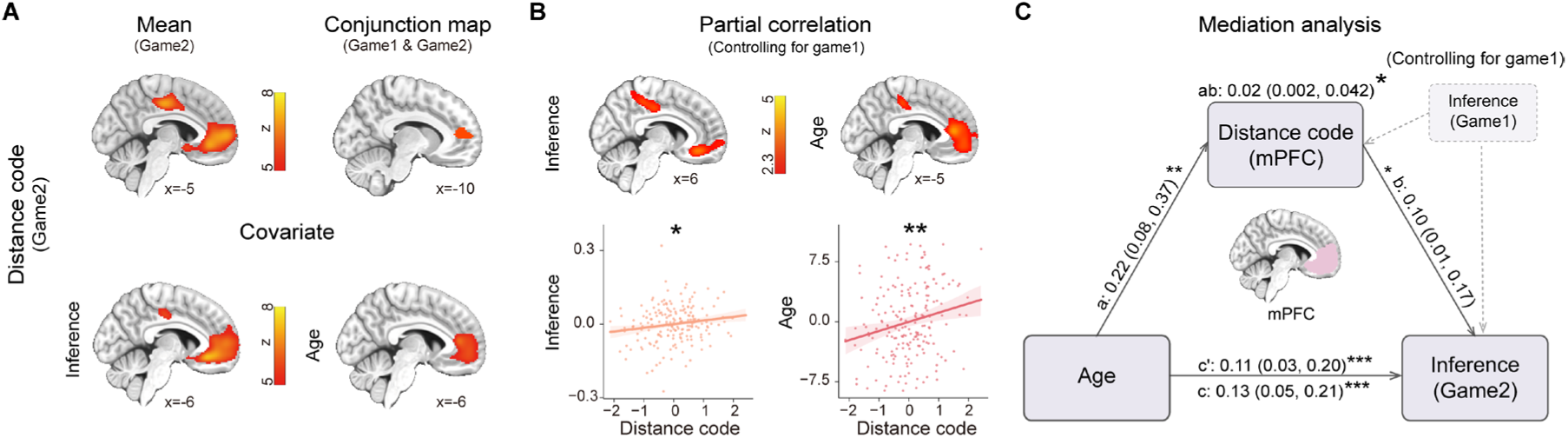
mPFC distance code selectively supports age-related development of nonlocal inference. **(A)** Whole-brain analysis reveals significant mPFC encoding of the Euclidean distance associated with new knowledge in Game 2 (top left). A conjunction analysis with Game 1 also shows overlapping mPFC activity (top right). Whole-brain covariate analyses using age or inference performance confirm consistent mPFC involvement in Game 2 (bottom). All maps are cluster-based FWE-corrected at *p* < 0.05 with a cluster-forming voxel threshold of *p* < 0.001 (see also **Figure S6**). **(B)** Both whole-brain and ROI analyses indicate that mPFC distance code strength correlates positively with nonlocal inference (left) and age (right), even when controlling for Game 1 performance. For visualisation, the whole-brain covariate maps are thresholded at *p* < 0.01, uncorrected. **(C)** A mediation analysis shows that the mPFC distance code mediates the effect of age on nonlocal inference in Game 2, even after controlling for Game 1 inference. Paths are labelled with coefficients and confidence intervals from 10,000 bootstrapped iterations. Each dot is data from individual subject. *: *p* < 0.05. **: *p* < 0.01. ***: *p* < 0.001.

Next, we asked whether assimilating new knowledge into the cognitive map differs from forming the original map, and how this process changes with age. After controlling for Game 1 performance, the mPFC remained correlated with both age and Game 2 nonlocal inference (Figure 5B upper panel), also confirmed at the ROI level (age: *r* = 0.21, *p* = 0.003, inference: *r* = 0.16, *p* = 0.02; Figure 5B bottom panel). A mediation analysis indicated that, even accounting for Game 1, the mPFC distance code continued to mediate the age-related increase in nonlocal inference (*β*_ab_ = 0.02, *p* = 0.04, CI [0.002, 0.042], Figure 5C). Intriguingly, this was not the case for the grid-like code, suggesting that building a new map representation in the mPFC is age-dependent and underpins rapid assimilation of novel knowledge, in addition to the EC grid signal.

### Structural basis of cognitive map through development

Brain structure also evolves significantly during development^23^, and these structural changes are linked to increased inference abilities^6^. To investigate the structural basis of cognitive map development and its connection to task-related representations during inference, we examined developmental changes in both grey-matter structure and white-matter connectivity (see STAR Methods), as well as their associations with cognitive map-based inferential abilities.

Using structural MRI, we measured grey matter volume (GMV) to assess cortical morphometry^33,34^. Total cortical GMV decreased between ages 8 and 25, consistent with the developmental trajectory depicted in the human “brain chart” project^23^ (Figure 6A). Notably, although this decreasing trend appeared at the whole-brain level, GMV in the EC continued to increase with age and inference ability from 8 to 25 (Figure 6B).

**Figure 6.**
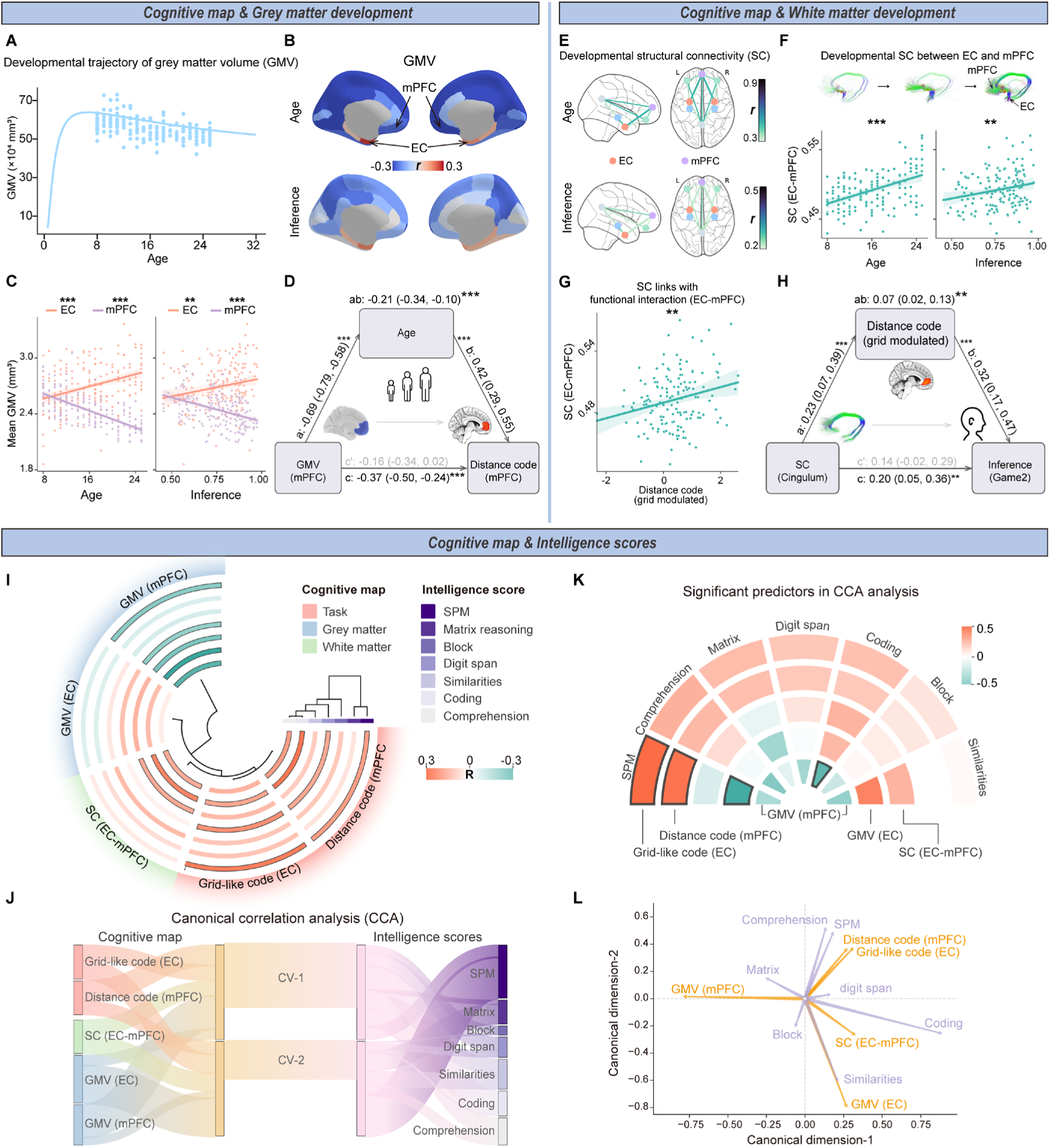
Neurodevelopmental basis of cognitive map and its link to intelligence measures. **(A)** The developmental trajectory of grey matter volume (GMV) from age 8 to 25, with data from this study (dots) overlaid on the ‘brain charts’. This aligns with the general decline in cortical GMV across adolescence. **(B)** Regional correlations between GMV and both age (top) and inference performance (bottom) for 34 bilateral regions from the Desikan–Killiany parcellation. Unlike most areas, EC increases with both age and inference ability, contrasting the overall decreasing trend elsewhere. **(C)** ROI analyses in the EC (coral) and mPFC (purple) highlight their divergent developmental patterns in GMV. **(D)** Mediation analysis shows that age-related changes in mPFC structure (measured by GMV) affect its on-task cognitive map representation (distance code). **(E)** Developmental structural connectivity (SC), measured by mean fractional anisotropy (FA) in white matter tracts. Tracts connecting the EC and mPFC exhibit the strongest correlation with age. The ROIs are EC (coral), mPFC (purple), HC (blue), LOFC (green), PCC (grey). **(F)** Age-related increases in EC–mPFC structural connectivity, illustrated by representative tract images at ages 8 (top left), 16 (top middle), 23 (top right). **(G)** Positive correlation between EC–mPFC structural connectivity and the EC grid-modulated distance code in Game2. **(H)** Mediation analysis reveals that the EC grid-modulated distance code, particularly via the cingulum, mediates the effect of EC– mPFC structural connectivity on Game2 inference performance. Statistical inferences used 10,000 bootstrapped iterations with paths labelled by coefficients and 95% confidence intervals. **(I)** Neural signatures of the cognitive map correlate with real-world reasoning scores in IQ tests. The heatmap shows the associations between intelligence measures (rows) and cognitive map measures (columns), segmented into five sets of measures and colour-coded (blue = negative; red = positive). Hierarchical lines denote clustering within each category. SPM refers to Raven’s Standard Progressive Matrices; other labels are WISC sub-tests. **(J)** A loading plot displays linear combinations of cognitive map measures for the canonical correlation analysis (CCA) derived canonical variates. The width of each streamline indicates the contribution of each variable to the canonical variable and the correlations among the canonical variates. **(K)** Coefficients of each cognitive map measure for predicting intelligence domains in the CCA model. **(L)** Projection of both sets onto the 2D canonical space, with vectors in purple for intelligence scores and orange for cognitive map measures, showing alignment and covariance among them. Significant results are outlined in black in (I) and (K). Each dot represents data from an individual subject. **: *p* < 0.01, ***: *p* < 0.001.

Focusing on the EC and mPFC (as defined in the task-fMRI analyses), both regions showed significant correlations with age and inferential performance. However, they differed in their developmental trajectories (age: *r*_*E*C_ = 0.36, *p* < 0.001; *r*_*m*PFC_ = −0.68, *p* < 0.001; inference: *r*_*E*C_ = 0.21, *p* = 0.004; *r*_*m*PFC_ = −0.38, *p* < 0.001; Figure 6C). We confirmed these findings across various parcellation methods (see **Data S13**), indicating that the developmental patterns are genuine and not artefacts of a specific atlas. We also observed a significant negative correlation between mPFC GMV and on-task cognitive map representations (the distance code) in both Game1 (*r* = −0.23, *p* < 0.001 and Game2 (*r* = −0.36, *p* < 0.001). No significant relationship was found in the EC between its structural changes and the grid-like code. A mediation analysis further showed that age-related structural changes in the mPFC influenced its task representation of the cognitive map (*β*_ab_ = −0.21, *p* < 0.001, CI [−0.34, −0.10], Figure 6D), indicating a developmental transformation of structure-function mapping in the mPFC.

Using diffusion MRI, we next measured structural connectivity by examining mean fractional anisotropy (FA) in white matter tracts^35^. Our results indicated that tracts linking the EC and mPFC showed the strongest age-related changes and were significantly correlated with inference performance (age: *r* = 0.48, *p* < 0.001; inference: *r* = 0.22, *p* = 0.006, Figure 6E & Figure 6F). Among these EC–mPFC tracts, the cingulum and uncinate correlated significantly with inference performance, whereas the fornix did not (**Figure S7**). Moreover, this EC–mPFC structural connectivity was closely associated with their functional interaction. Specifically, we observed a correlation with the EC grid-modulated distance code in the mPFC during Game2 (SC_EC-mPFC_: *r* = 0.24, *p* = 0.003, cingulum: *r* = 0.23, *p* = 0.004, Figure 6G). Further analysis revealed that this EC grid-modulated distance code mediated the effect of EC–mPFC structural connectivity on Game2 inference performance (SC_EC-mPFC_: *β*_ab_ = 0.07, *p* = 0.001, CI [0.03, 0.13]), primarily through the cingulum (cingulum: *β*_ab_ = 0.07, *p* = 0.01, CI [0.02, 0.13], Figure 6H), rather than the uncinate or the fornix (all *p* > 0.05).

Together, these findings suggest that the maturation of the EC and mPFC, along with their structural connectivity, underpins the development of task-related representations of cognitive map and the associated flexible behaviour.

### Cognitive map-related measures generalise to standardised intelligence tests

Next, we explored how the development of the cognitive map relates to real-world cognitive abilities measured by standardised intelligence quotient (IQ) tests. Rather than simply identifying correlations, our goal was to identify which aspects of intelligence, as assessed by these tests, align most closely with cognitive map-related measures during development.

We employed two prominent IQ assessments: Raven’s Standard Progressive Matrices (SPM)^36^ and the Wechsler Intelligence Scale for Children (WISC)^37^. These tests evaluate a broad range of cognitive abilities, from working memory (e.g., digit span) to higher-level skills such as reasoning and comprehension of real-world concepts (e.g., ‘explaining why seat belts are used’). We found significant correlations between neural measures of the cognitive map—including task representations, brain structural development, and white matter connectivity—and various IQ scores (Figure 6I). For instance, participants with higher SPM scores tended to perform better on the inference task (*r* = 0.52, *p* < 0.001) and required fewer training sessions to achieve proficiency (*r* = –0.34, *p* = 0.001). Mediation analyses further revealed that SPM scores partially mediated the age-related improvements in inference performance (*β*_ab_ = 0.15, *p* < 0.001, CI [0.08, 0.25]).

To delve deeper, we applied canonical correlation analysis (CCA)^38^, which allowed us to identify combinations of neural cognitive map measures and intelligence test scores that share the most variance (Figure 6J). We used a strict permutation framework^39^ to minimise false positives. Only the first two canonical variates reached significance (CV-1: *r* = 0.64, *p* < 0.001; CV-2: *r* = 0.39, *p* = 0.009; corrected for multiple comparisons).

Notably, both the EC grid-like code and the mPFC distance code significantly predicted SPM reasoning scores (*β*_grid code_ = 0.55, *p* = 0.04; *β*_distance code_ = 0.52, *p* = 0.05; Figure 6K). To visualise these relationships, we projected the original measures onto the space defined by the two significant canonical dimensions (Figure 6L). This projection revealed that particular neural measures—specifically the grid-like code and distance code—were closely associated with SPM reasoning and WISC comprehension.

These findings, linking the cellular representation to standardised IQ tests, highlight that the developing neural representation of cognitive map is intrinsically linked to children’s expanding capacity for abstract reasoning and real-world comprehension in everyday life.

### Hippocampus contributes to inference and learning in children

Thus far, our work has focused on how inferential reasoning develops within a cognitive map. A key question, however, is how children with less coherent grid codes can still perform the task. Using a median split, we confirmed that children (aged 8–12) with low grid-code nevertheless achieved above-chance inference accuracy (*t*(36) = 8.07, *p* < 0.001), although their performance was significantly lower than of peers with stronger grid codes (*t*(73) = −2.47, *p* = 0.02; Figure 7A).

**Figure 7.**
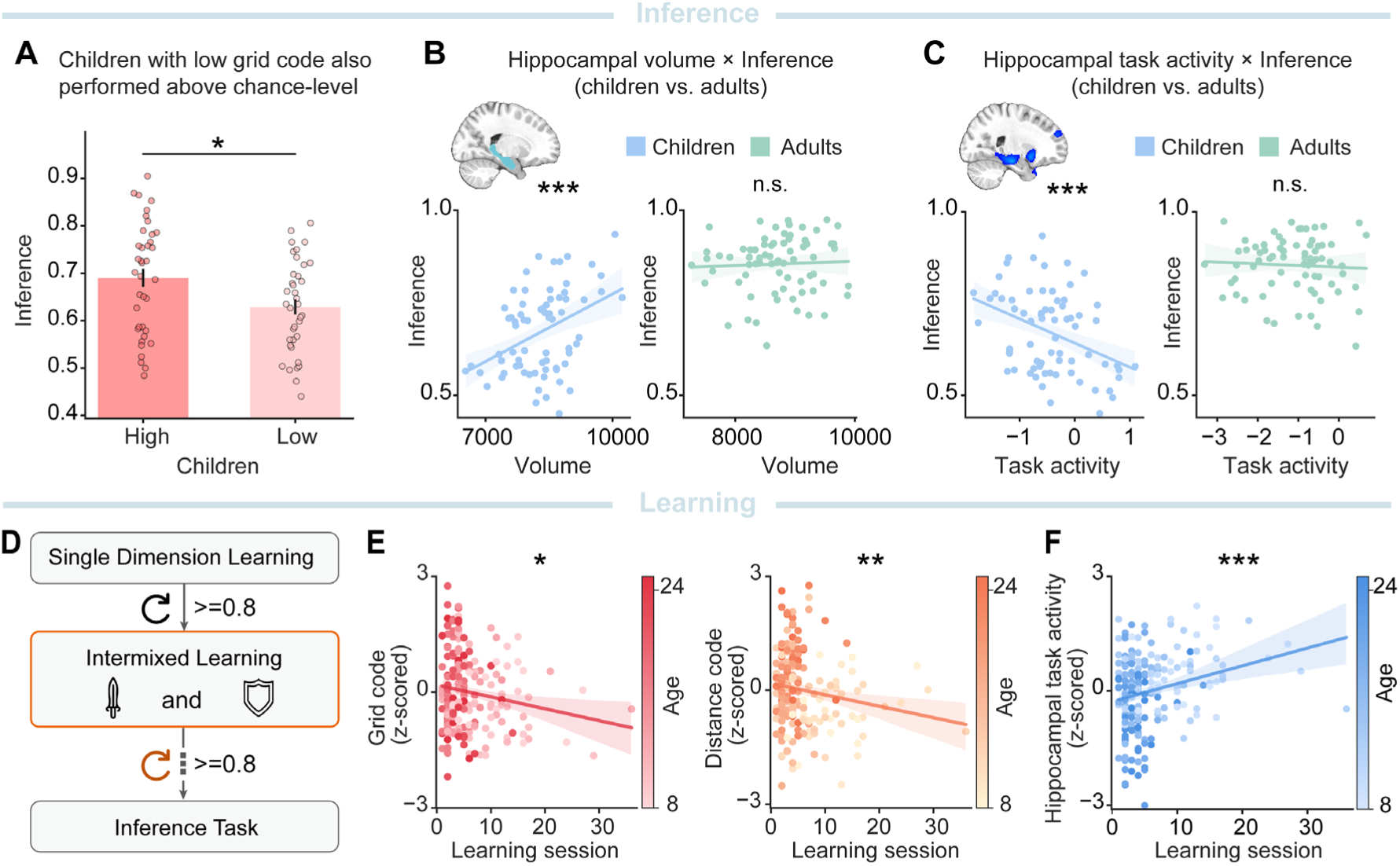
Hippocampus contributes to inference and learning in children. **(A)** Children (aged 8–12) with low grid code still achieved above-chance inference performance, though their accuracy was significantly lower than their peers with high grid code (median split). **(B)** Hippocampal volume positively correlated with children’s inference performance (averaged over two tasks) for ages 8–12, but not in adults. **(C)** Hippocampal activity during inference time negatively correlated with inference performance in children but not in adults. **(D)** Schematic of the training protocol (see also **Figure S1** and Methods), including single-dimensional and intermixed learning. Participants proceeded to the formal inference task only after reaching ≥80% accuracy on memory probes of all pairs (without feedback) during the intermixed phase. **(E)** The number of learning sessions required to reach ≥80% accuracy in intermixed learning was negatively correlated with both EC grid code (left) and mPFC distance code (right), whereas **(F)** it was positively correlated with hippocampal activity during inference time. Each dot denotes one participant. Error bars denote s.e.m. The line represents the best linear fit, and the shaded area indicates the 95% confidence interval. *: *p* < 0.05, **: *p* < 0.01, ***: *p* < 0.001. n.s. non-significant.

One possibility is that these children build a direct associative map, such as a successor representation^10–12^, when they have not yet formed a robust grid-schema representation. Replay is known to facilitate this associative map formation^40,41^, and in rats it emerges early, during the fourth postnatal week^42^. It is therefore plausible that younger individuals might rely more on replay. In a complementary MEG experiment on over one hundred participants, aged 8–25, performing a similar inference task and resting-state periods^43^, we indeed found that younger children showed selectively stronger replay of experiences for building the map during post-task rest.

Although we cannot directly observe replay events with fMRI in the current study, this mechanism predicts heavier reliance on the hippocampus in children, which we tested. Indeed, while we found no significant relationships with distance code, GMV of the EC or mPFC, and their structural connectivity (all *p* > 0.05), only hippocampal volume correlated positively with inference ability in children who had weaker grid codes (*r* = 0.43; surviving Bonferroni correction, *p*_corr_ = 0.04), and remained significant across all children (*r* = 0.40, *p* = 0.001; Figure 7B, averaged over both inference tasks). This effect persisted even after adjusting for age (*r* = 0.40, *p* = 0.001), and was absent in adults (*r* = 0.05, *p* = 0.70), with a significant difference between the two groups (Fisher *z* = 2.20, *p* = 0.03). Further evidence came from analysis of children’s hippocampal activity during inference time, which showed a negative correlation with inference performance (*r* = −0.38, *p* = 0.001; Figure 7C), whereas adults demonstrated no such effect (*r* = −0.05, *p* = 0.65; children vs. adults: Fisher *z* = −2.02, *p* = 0.04). These findings suggest that children with less coherent EC grid codes can rely more on the hippocampus when solving the task.

This reliance on hippocampal mechanisms may also be reflected in learning patterns, where children were required to memorise 200 unique pairs (Figure 7D; see also STAR Methods “Training procedure”). We observed a pronounced developmental shift in learning efficiency: children aged 8–12 required significantly more sessions (20.05 ± 1.49) than adolescents (13–17 years; 9.11 ± 0.37) or adults (18–25 years; 8.00 ± 0.29) to reach ≥ 80% accuracy. Notably, the total number of learning sessions did not correlate with final memory on test day (*r* = −0.10, *p* = 0.14), indicating that extended training did not bias their memory basis for the formal inference tasks.

The largest age differences appeared in the intermixed learning phase (**Figure S1** and **Data S14**), which tested participants on pairwise memory across both dimensions. Focusing on this phase, we found that the EC grid code and mPFC distance code negatively correlated with the number of learning sessions (Figure 7E; grid: *r* = –0.16, *p* = 0.02; distance: *r* = – 0.22, *p* = 0.002), whereas hippocampal activity during inference time positively correlated (Figure 7F; *r* = 0.25, *p* < 0.001). These patterns imply that children who required more practice also relied heavily on the hippocampus for inference, a dependence that diminishes with development as a more structured (e.g., grid-like) knowledge schema emerges.

## DISCUSSION

Our study tried to understand the neural mechanisms underlying cognitive development by examining how structured knowledge emerges with age, leading to more intelligent behaviour. We found that the maturation of grid-cell-like codes—a fundamental cellular mechanism in the EC representing 2D knowledge maps—underpins both inferential reasoning and the assimilation of new information. These grid-like codes strengthen over time and enable the mPFC to encode relationships between concepts during inference. Notably, their development also aligns with standardised IQ measures, particularly in abstract reasoning and understanding of real-world relational concepts.

Maintaining a stable schema helps individuals infer novel concepts more effectively, potentially via path-integration mechanisms^44,45^. In physical space, grid cells in rodents generalise structural regularities across different environments^15,17^, and our findings reveal a similar grid-like schema code for a 2D knowledge map in humans. Its consistency across original and new maps facilitates the assimilation of novel information. As individuals grow older, their grid code becomes stronger, supporting increasingly effective inference. This improvement likely reflects their progressively refined ability to organise experience into a coherent schema, providing a basis for inferring knowledge beyond direct observations. Such a broad role for grid-like codes in structuring conceptual knowledge resonates with the notion of “core knowledge”^46^, or the “universal law of generalisation”^47,48^, implying innate cognitive map representation that scaffold development^49^.

Forming a map-like representation in the current study goes beyond memory capacity alone. Previous research suggested that young children struggle with memory integration in associative inference tasks^50,51^. In the current findings, we controlled for memory differences, yet age-related increases in inferential reasoning and assimilation remained. Although children could solve some inference tasks by reactivating individual memories rather than forming a unified 2D map, a schema-based grid-like code in the EC confers greater efficiency. Representing a 5 × 5 map, for example, requires recalling only 25 object–location relationships rather than 400 pairwise associations, aligning with theories of more efficient memory reorganisation^52,53^.

In children with less coherent grid codes, we observed weaker inference abilities compared with their peers who had a stronger grid schema; nevertheless, these children still performed above chance. Our data suggest they may rely on an alternative hippocampal mechanism. The hippocampus can support statistical learning^6,54^ or replay mechanisms^55^ that may enable children to directly build the associative map^41^. Although this alternative route is less effective than a coherent grid-based schema, it may allow children to compensate for weaker grid codes and achieve moderate task performance, possibly through increased replay^43^, a hypothesis that warrant further investigation.

We also observed that a stable schema in the EC facilitates building new map representations when linking novel information to established knowledge. The mPFC is critical in this process^56^, and our data show it encodes the distance relations between new and existing concepts, an ability that strengthens with age. In contrast to the EC grid signal, the mPFC distance coding exhibited an independent, developmental role in assimilating new knowledge, pointing to an additional mechanism of assimilation in the mPFC, beyond the fundamental schema representation.

The mPFC continues to mature through adolescence, undergoing structural changes that include grey matter volume decreases^57–59^. By contrast, we found the EC grey matter volume rises, indicating different developmental trajectories of these regions in supporting the cognitive map. We also found significant age-related increase in structural connectivity between the EC and mPFC, particularly via the cingulum bundle rather than indirect pathways such as the fornix. This bundle is crucial to the default mode network, connecting the mPFC and PCC, regions previously associated with grid-like representations^19,20^.

These functional and structural changes coincide with the late adolescent emergence of theory of mind^60,61^. Adolescents show increased mPFC engagement in mentalising tasks compared to adults^62^. Our findings suggest the neural mechanisms underlying mentalising and cognitive map may share overlapping processes. These have important educational implications. Learning fundamentally involves integrating new ideas into existing schemas^63^. Our further CCA results showed that neural markers of cognitive map correlate with IQ tests assessing reasoning and comprehension, which is especially relevant during adolescence when the neural basis of the cognitive map is still maturing. Strengthening schema-based strategies in educational settings, and encouraging students to connect new concepts to their prior knowledge, could thus foster more advanced reasoning and comprehension, offering real-world benefits^64^.

### Limitations of the study

Our study addresses how neural representations of structured knowledge during inference emerge and develop with age. However, we did not record neural activity during the learning process itself, leaving it unclear how learning shapes these representations, especially in children. Consequently, it is difficult to determine whether younger participants struggle to effectively use an already-established cognitive map during inference tasks or whether they face difficulty forming this map during learning. Moreover, our cross-sectional design uses age as an instrumental variable in mediation analyses, which suggests statistical association but does not prove a causal link between grid code development and reasoning ability. Finally, although reasoning is a crucial component of intelligence, IQ encompasses multiple dimensions beyond the scope of this study. Future studies could consider longitudinal designs to clarify the neural trajectory throughout learning and explore other facets of intelligence.

## Conclusion

In conclusion, our findings demonstrate that the development of grid-like codes in the EC underpins inferential reasoning and assimilation. The mPFC supports this process by mapping new information in an age-dependent manner, enabling more sophisticated knowledge integration over time. These functions closely relate to structural maturation in both the EC and mPFC, along with their connectivity, laying the groundwork for uncovering the cellular mechanisms underlying cognitive development and intelligence.

## RESOURCE AVAILABILITY

### Lead contact

Further information and requests for resources and reagents should be directed to and will be fulfilled by the lead contact, Yunzhe Liu.

### Materials availability

This study did not generate new unique reagents.

### Data and code availability

fMRI and behavioural data that support the conclusions in this study is available on https://osf.io/ajz6g/ upon publication. The raw individual participant MRI data will be available upon reasonable request to the corresponding author, subject to participant consent. The analysis code is available on https://gitlab.com/liu_lab/develop_cognitive_map.git upon publication.

## ACKNOWLEDGMENTS

This study is supported by the National Science and Technology Innovation 2030 Major Program (2022ZD0205500), the National Natural Science Foundation of China (32271093), the Beijing Natural Science Foundation (Z230010, L222033), and the Fundamental Research Funds for the Central Universities.

## AUTHOR CONTRIBUTIONS

Conceptualization, Y.L., Y.Q., and T.B.; Investigation, Q.Y., J.O., L.P., S.W., Y.Luo., Y.L., and T.B.; Writing – Original Draft, Y.Q., Y.L.; Writing – Review & Editing, Y.Q., Y.L., J.O., and T.B.

## STAR★METHODS

### KEY RESOURCES TABLE

| REAGENT or RESOURCE | SOURCE | IDENTIFIER |
| --- | --- | --- |
| Deposited data |  |  |
| MRI and behaviour data | This paper | https://osf.io/ajz6g/ |
| Software and algorithms |  |  |
| fMRIPrep (v20.2.7) | Esteban et al.<sup>66</sup> | https://fmriprep.org/en/stable/ |
| FMRIB Software Library (v6.0.5) | Smith et al.<sup>102</sup> | https://fsl.fmrib.ox.ac.uk/fsl/docs/#/ |
| Freesurfer (v7.2.0) | Dale et al.<sup>33</sup> | https://surfer.nmr.mgh.harvard.edu/ |
| Nipype (v1.8.5) | Gorgolewski et al.<sup>67</sup> | https://nipype.readthedocs.io/en/latest/ |
| Nilearn (v0.10.4) | Abraham et al.<sup>103</sup> | https://nilearn.github.io/stable/index.html |
| rsatoolbox (v0.1.3) | Nili et al.<sup>106</sup> | https://rsatoolbox.readthedocs.io/en/stable/ |
| QSIprep (v0.19.0) | Cieslak et al.<sup>80</sup> | https://qsiprep.readthedocs.io/en/latest/ |
| MRtrix3 (v3.0.4) | Tournier et al.<sup>96</sup> | https://www.mrtrix.org/ |
| Pingouin (v0.5.3) | Vallat et al.<sup>101</sup> | https://pingouin-stats.org/ |
| scikit-learn (v1.3.0) | Pedregosa et al.<sup>105</sup> | https://scikit-learn.org/stable/ |
| statsmodels (v0.14.0) | Seabold et al.<sup>104</sup> | https://www.statsmodels.org/stable/index.html |
| Custom code | This paper | https://gitlab.com/liu_lab/develop_cognitive_map.git |
| Other |  |  |
| Magnetic Resonance Imaging system | Siemens Prisma 3T scanner | https://www.siemens-healthineers.com/magnetic-resonance-imaging/3t-mri-scanner/magnetom-prisma |

### EXPERIMENTAL MODEL AND STUDY PARTICIPANT DETAILS

We recruited a total of 231 subjects in the fMRI experiment, with ages ranging from 8 to 25 years. Of these, 163 were between the ages of 8 and 18, ensuring a minimum of 10 subjects per age group. The remaining 68 were adults, aged 19 to 25, selected to reflect the later developmental stages of the mPFC as indicated by previous studies^23,24^. Subjects who were under 16 years old, were also scored on the Raven’s Standard Progressive Matrices (SPM) test^36^ and the Wechsler Intelligence Scale for Children (WISC) in accordance with the WISC grading norms for the Chinese population^37^. After excluding 28 participants (7 adults and 21 children) due to reasons such as failed memory probe (accuracy < 0.8), excessive head movement during fMRI (with a mean FD exceeding 0.2 mm), and technical issues, we had a total of 203 participants (age: 15.64 ± 0.37, 115 female) for the primary analysis (**Table S1**). Among them, 193 (age: 15.77 ± 0.39, 109 female) also undertook the subsequent knowledge assimilation phase after the primary inference task. In addition to the fMRI experiment, all participants underwent structural MRI scans, and 147 obtained diffusion MRI scans (age: 16 ± 0.43, 85 female). Before the actual experiment, those under 12 years old attended a mock scanner session. In this session, they also watched a video introducing them to the sounds of magnetic resonance and highlighting the need to stay calm and motionless during the scan. This step helped younger participants practice head stability and become more comfortable with the MRI environment. All participants reported correct understanding of the task goal and instructions.

The experiment protocol was approved by the Institutional Review Board of the Faculty of Psychology, Beijing Normal University (ethics number: IRB_A_0051_2021002). All participants were healthy with no history of psychiatric or neurological disorders, and provided informed consent before the task. For participants under 18 years old, parental consent was obtained. All procedures adhered to the standards set by the Institutional Review Board of Beijing Normal University.

## METHOD DETAILS

### 2D Knowledge map

The knowledge map consists of a 5×5 grid spanning two independent dimensions: attack (AP) and defence power (DP), each with 5 rank levels. As in Park *et al.* ^19^, participants understood these ranks as equidistant intervals, enabling us to measure the hypothesized vector angle in the task space. The task space features 25 unique visual stimuli (monsters in the cover story). The association between a stimulus and its task space position is randomized for each participant, preventing any stimulus preference from skewing the task space representation on average.

During training, participants were not informed of the structure of the 2D knowledge map, such as the number of ranks per dimension, the count of stimuli sharing a rank, or their specific arrangement. Instead, they were given information specifically about the pairwise one-rank difference relationship within each dimension. In the formal fMRI task, our goal was to determine if and how participants, drawing from memory, constructed the knowledge map to enable flexible inference beyond just pairwise relationships.

### Training procedure

Our training procedure was designed to ensure that all participants, regardless of age, memorised all pairwise relationships before proceeding to the formal fMRI experiment. This training was divided into three stages and could span multiple days if participants could not consistently meet the predefined standard. Across daily one-hour sessions (mean days: 12.75 ± 0.69; see also **Figure S1** for procedure), each participant learned 200 pairwise relationships.

### Single Dimension Learning

In this first stage, participants learned one-rank difference relationships for each dimension (DP and AP) separately. On any given day within this stage, they focused on only one dimension—either AP or DP—and the learning order of DP and AP was randomised across individuals. Each of the 100 unique pairwise relationships in a dimension was shown twice with the positions swapped, producing 200 presentations per dimension. To reduce the burden, these 200 trials were split into 20 blocks, each containing 10 stimulus pairs.

Within each block, participants underwent two parts: a learning part and a test part. During learning, they saw two stimuli differing by one rank in the specified dimension. A symbol at the bottom of the screen indicated whether AP or DP was being trained, while a “>” or “<” sign in the centre showed the relative rank. To accommodate varying learning speeds across ages, participants advanced at their own pace, pressing a button when they had memorised each pair. In the test part, the same pairs were shown without the magnitude symbols, and participants selected the stimulus with the higher rank in the dimension. Following each response, a feedback symbol (“correct” or “incorrect”) was displayed for 1.5 seconds. Participants repeated each block until reaching at least 80% accuracy. After completing all blocks, they underwent a probe test without feedback featuring all learned pairs, again requiring at least 80% accuracy to move on; otherwise, they repeated this stage.

### Intermixed Learning

The second stage kept the self-paced learning format but interleaved DP and AP trials within a block. It also comprised two parts: a memory probe and a relearning part. During the memory probe, participants were tested on 200 unique pairs (100 DP + 100 AP), with randomly swapped positions and sequences. They identified the higher-ranked stimulus without feedback. This design prevented order effects and ensured participants retained both AP and DP ranks for each monster. As before, participants needed ≥80% accuracy across both dimensions. Failing this criterion led to the relearning phase, where they revisited incorrectly answered pairs under the same self-paced regime.

### Review

During the three days preceding the formal fMRI experiment, participants reviewed all learned pairs in a self-paced manner. This final step ensured that all participants, irrespective of age, entered the main task with the necessary information to construct the mental map. Consequently, if younger participants performed less well at cognitive map-based inference in the fMRI study, it cannot be attributed to inadequate information. The training took place online with daily progress tracking. Each day, experimenters reminded participants to complete the scheduled learning tasks and recorded their progress. Each full practice run was noted as one training session. Training ended only when participants achieved at least 80% accuracy or voluntarily withdrew. This adaptive method accommodated individual learning rates among different age groups while maintaining a consistent memory performance standard.

### Formal fMRI experiment

Subjects who had passed the training procedure were invited for the formal fMRI experiment. Upon arrival, they were probed again on their knowledge retention of all pairwise relationships. Probing trials for both dimensions were randomized, and no feedback was provided. The test performance served as an index for training accuracy (see Figure 1C), the memory accuracy was 91.79 ± 0.34 %.

To ensure that all participants understood the task, each subject completed a pilot inference task. This pilot task had the same structure as the formal inference task but used different stimulus pairs (24 trials). This helped younger participants learn the rules and become familiar with the task. After completing the pilot task, participants were asked to report the task rules to confirm their understanding.

Subjects then participated in a 2D inference task while undergoing concurrent fMRI scanning. Unlike traditional transitive inference tasks, this required integrating information from both dimensions. Participants had to determine if the AP order of one monster was higher than the DP order of another. The decision rule, which identified the attacker and defender, varied trial by trial and was only revealed after the presentation of the second monster (M2). This design encouraged subjects to maintain a 2D knowledge map in mind during M2 presentation, and only compute value upon the decision rule. This was shown to be effective (see **Figure S4**).

Following this, subjects were invited to engage in a novel knowledge assimilation task. In this phase, four new objects were introduced to the map for the first time. After learning their relationships with nearby monsters on the map (local learning), subjects were tasked to infer the relationships between these new entities and non-adjacent ones, with which they had no direct learning experience (non-local inference), all while undergoing fMRI scanning.

### 2D inference task

During the fMRI scanning, the main task involved participants inferring relationships between two monsters based on their ranks in both AP and DP dimensions, termed “2D inference”. Each trial began with a 0.5s fixation, followed by the appearance of the first monster (M1) for 1-4s. The second monster (M2) then appeared for 2.5s, followed by a jittered inter-stimulus-interval (ISI) of 1-4s. Upon the appearance of M2, participants could mentally navigate the 2D knowledge map by drawing a hypothesized vector that connected M1 and M2. Subsequently, the decision rule, accompanied by the target dimension symbols for both monsters, was presented for up to 3s and disappeared upon choice. Each trial ended with an inter-trial-interval (ITI) that lasted between 0.3-3.3s. The main task consisted of 252 trials, divided into 6 runs, covering all possible angles on the 5×5 map with 21 trials for each angle bin (12 bins in total, equally spanned 0-2π). The presentation order of monsters within-pair were randomised.

### Knowledge assimilation task

To investigate how the cognitive map was used to assimilate new knowledge, participants next performed a knowledge assimilation task. Four new monsters, each represented by a unique visual stimulus, were introduced near the centre of the existing map. During this phase, participants learnt the relative rank differences between these new monsters and the adjacent existing ones, with each new monster connected to four neighbouring items in the established map—referred to as “local learning.” Participants were informed that relationships among existing objects would remain unchanged, allowing them to integrate the new knowledge into a stable framework. Although the new monsters were positioned at 0.5 rank intervals relative to existing items, this spacing was not explicitly disclosed to participants. Instead, they were only instructed to learn the relative rankings between novel and existing items while preserving the established relationships. Efficient assimilation here required integrating these new relationships into the existing knowledge structure.

This “local learning” was performed in block format, comprising 256 trials equally divided across four blocks (64 trials per block). Each local-pair relationship was presented eight times per dimension. Block presentation followed either an ADDA or DAAD sequence for the two dimensions (A for attack power, D for defence power), counterbalanced across participants. In each trial, participants viewed pairs of new stimuli with their relative magnitudes indicated by directional symbols in a self-paced manner as before. After these learning blocks, they completed a memory probe test, achieving a mean accuracy of 83.32 ± 1.01%.

Subsequently, participants undertook a “non-local inference” task (or “Game 2”), identical in timing and trial structure to the main task but focusing solely on relationships between the newly introduced monsters and non-local existing monsters that were not included in the learning set. This non-local inference phase was conducted under fMRI scanning. In total, 84 unique pairs were presented, with each new monster paired with 21 non-local existing monsters. Trials were randomised and evenly split across two runs.

### Behavioural analysis

To evaluate how inference performance improved with age, we used Pearson correlations and partial correlations on map-based inference (Game1) and nonlocal inference (Game2). For Game1, the partial correlation analysis included training accuracy as a covariate to control for memory performance, which also tend to improve with age. To assess the selective developmental impact on knowledge assimilation in Game2, we conducted a partial correlation, adjusting for Game1 performance and the learning performance of newly introduced objects. Additionally, linear regression models were performed to support the findings both in Game1 and Game2. In Game1, the model including age and memory performance, showed a significant age effect on map-based inference performance. In Game2, the regression model included age, Game1 performance, and memory performance during training and new knowledge learning as explanatory variables, with nonlocal inference as the dependent variable.

### MRI data acquisition

We acquired structural image and functional images on Siemens Prisma 3T scanner. T2-weighted functional images were acquired using an echo planar imaging (EPI) sequence with the following parameters: 49 slices in an interleaved order repetition time, resolution = 3 × 3 × 3 mm^3^, repetition time (TR) = 3000 ms, echo time (TE) = 30 ms, flip angle = 90°, field of view (FoV) = 192 mm. We used a 30° slice angle relative to the anterior-posterior commissure line to minimize signal loss in the orbitofrontal cortex region ^65^. We also acquired a T1-weighted structural image using a magnetization-prepared rapid gradient echo (MPRAGE) sequence with the following parameters: resolution = 0.5 × 0.5 × 1 mm^3^, TR = 2530 ms, TE = 2.98 ms, flip angle = 7°, FoV = 256 mm. The diffusion data were acquired using a spin-echo diffusion-weighted sequence with the following parameters: resolution = 2 × 2 × 2 mm^3^, 64 diffusion direction, TR = 2800 ms, TE = 75 ms, b = 0, 1000, 2000, 3000 s/mm^2^, PA phase encode direction. Phase-reversed b0 images were collected to derive a fieldmap for distortion correction of diffusion data.

### MRI Data preprocessing

Results included in this manuscript come from preprocessing performed using fMRIPrep 20.2.7^66^ and QSIPrep 0.19.0, which are based on Nipype^67^.

### Structural data preprocessing

The T1-weighted (T1w) image was corrected for intensity non-uniformity (INU) with N4BiasFieldCorrection^68^, distributed with ANTs 2.3.3^69^, and used as T1w-reference throughout the workflow. The T1w-reference was then skull-stripped with a Nipype implementation of the antsBrainExtraction.sh workflow (from ANTs), using OASIS30ANTs as target template. Brain tissue segmentation of cerebrospinal fluid (CSF), white-matter (WM) and grey-matter (GM) was performed on the brain-extracted T1w using fast^70^. Volume-based spatial normalization to one 2 mm standard space (MNI152NLin2009cAsym) was performed through nonlinear registration with antsRegistration (ANTs 2.3.3), using brain-extracted versions of both T1w reference and the T1w template. The following template was selected for spatial normalization: ICBM 152 Nonlinear Asymmetrical template version 2009c MNI152NLin2009cAsym^71^.

### Functional data preprocessing

For each run, the following preprocessing was performed. First, a reference volume and its skull-stripped version were generated using fMRIPrep. A deformation field to correct for susceptibility distortions was estimated based on fMRIPrep’s fieldmap-less approach. The deformation field is that resulting from co-registering the BOLD reference to the same-subject T1w-reference with its intensity inverted^72,73^. Registration is performed with antsRegistration (ANTs 2.3.3), and the process regularized by constraining deformation to be nonzero only along the phase-encoding direction, and modulated with an average fieldmap template^74^. Based on the estimated susceptibility distortion, a corrected EPI (echo-planar imaging) reference was calculated for a more accurate co-registration with the anatomical reference.

The BOLD reference was then co-registered to the T1w reference using flirt^75^ with the boundary-based registration^76^ cost-function. Co-registration was configured with nine degrees of freedom to account for distortions remaining in the BOLD reference. Head-motion parameters with respect to the BOLD reference (transformation matrices, and six corresponding rotation and translation parameters) are estimated before any spatiotemporal filtering using mcflirt^77^.

BOLD runs were slice-time corrected to 1.46s (0.5 of slice acquisition range 0s-2.92s) using 3dTshift from AFNI 20160207^78^. The BOLD time-series (including slice-timing correction when applied) were resampled onto their original, native space by applying a single, composite transform to correct for head-motion and susceptibility distortions. These resampled BOLD time-series will be referred to as preprocessed BOLD in original space, or just preprocessed BOLD. The BOLD time-series were resampled into standard space, generating a preprocessed BOLD run in MNI152NLin2009cAsym space.

Several confounding time-series were calculated based on the preprocessed BOLD: FD and three region-wise global signals. For each run, FD with root mean square matrix were calculated following the definitions by Jenkinson *et al.* ^77^. The three global signals are extracted within the CSF, the WM, and the whole-brain masks. The head-motion estimates calculated in the correction step were also placed within the corresponding confounds file. Frames that exceeded a threshold of 0.5 mm FD were annotated as motion outliers.

All resamplings can be performed with a single interpolation step by composing all the pertinent transformations (i.e., head-motion transform matrices, susceptibility distortion correction when available, and co-registrations to anatomical and output spaces). Gridded (volumetric) resamplings were performed using antsApplyTransforms (ANTs), configured with Lanczos interpolation to minimize the smoothing effects of other kernels^79^. Non-gridded (surface) resamplings were performed using mri_vol2surf (FreeSurfer).

### Diffusion data preprocessing

Preprocessing of the diffusion data was conducted using QSIPrep 0.19.0, built on Nipype 1.8.6^67,80^. In this process, images with a b-value less than 100 s/mm^2^ were classified as b=0 images. We applied MP-PCA denoising, using MRtrix3’s dwidenoise tool with a 5-voxel window, following the method outlined by Veraart et al. ^81^. Post-denoising, we adjusted the mean intensity of the DWI series so that the mean intensity of b=0 images was matched across different DWI scans. Correction for B1 field inhomogeneity was done using MRtrix3’s dwibiascorrect with the N4 algorithm^68^.

For head motion and Eddy current correction, we employed FSL’s eddy tool, following the guidelines of Andersson and Sotiropoulos ^82^. Our configuration for eddy included a q-space smoothing factor of 10, five iterations, and the use of 1000 voxels for estimating hyperparameters. We applied linear models at both first and second levels to account for Eddy current-related spatial distortion. Shells were assigned to q-space coordinates, and an effort was made to separate field offset from subject movement. Eddy’s outlier replacement was run following Andersson et al. ^83^. We grouped the data by slice and any group deviating more than four standard deviations from the prediction had their data replaced with imputed values.

Data collection involved the use of reversed phase-encode blips to generate image pairs with opposite directional distortions. For this, we used b=0 reference images with reversed phase encoding and an equal number of b=0 images from the DWI scans. The susceptibility-induced off-resonance field was estimated from these pairs, following^84^. The resulting fieldmaps were then incorporated into the interpolation process for Eddy current and head motion correction, with the final interpolation being performed using the ‘jac’ method.

In the final stage of preprocessing, we calculated several confounding time-series based on the preprocessed DWI, including FD. Finally, the DWI time-series were resampled to ACPC, resulting in a preprocessed DWI run in ACPC space with 2 mm isotropic voxels.

### Functional MRI data analysis

We performed general linear model (GLM) to capture event-related activations. The preprocessed data were smoothed with an 8-mm full-width at half maximum Gaussian kernel and filtered by a high-pass cutoff filter of 100 Hz. All GLMs modelled three basic events (M1, M2, Decision) as regressors. This was achieved by using a boxcar function with the corresponding duration, which accounted for the mean response specific to those visual stimuli that were not of interest (see **Figure S5** for related activation patterns). All regressors were convolved with a canonical hemodynamic response function. In an effort to mitigate the effects of potential head motion noise, all GLMs incorporated six rigid-body motion parameters (3 translation and 3 rotation parameters), in addition to the motion outliers, cerebrospinal fluid (CSF) and white matter (WM) signals in each run.

We modelled parametric regressors for different target representations: grid-like code, distance code, and value code, as described below. Further second-level covariate analysis were performed with one-sample t-test using age or inference performance as the covariate.

### Grid-like code

To detect grid-like code in fMRI, we used three established and independent methods: first, a hexagonal modulation analysis that relies on the *z*-transformed *F*-statistic of inferred trajectories^20,27^; second, an analysis of the hexagonal consistency effect, which compares inferred trajectories with the mean orientation of a specific brain region, such as the EC^18,19^; and third, a similarity analysis focused on the neural representations of angular differences in inferred trajectories^28^.

In the hexagonal modulation analysis, we began by determining the angle, θ, of the inferred trajectory. This process starts with the presentation of M2, from which a trajectory on the 2D map can be inferred. Consequently, the GLM modelled a six-fold sinusoidal modulation with two parametric regressors: sin6θ and cos6θ at the time of M2’s presentation and decision. Here, θ represents the direction of inferred trajectories in each trial. We modelled the events for correct trials and error trials separately, setting those parametric modulations on correct trial events. The parametric estimates of these two regressors form a quadrature filter, with its amplitude 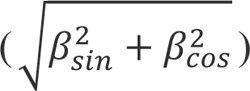 being sensitive to hexagonal symmetry. To compute this amplitude, we applied an *F*-test to detect any linear combination of these two regressors: β_*s*in_∗ sin(6θ) + β_*c*o*s*_∗ cos(6θ)^18,20^. Then, we transformed the statistic at the first level into *z*-statistic based on an asymptotic approximation for further group-level analysis^85^. This analysis was conducted for both Game1 and Game2. Similar to previous studies^20^, the *F*-test-based method didn’t account for consistent mean orientation across runs. It allowed us to define a functional EC ROI in an unbiased manner. This mask was just used to estimate mean orientation and included voxels with hexagonal symmetry at a threshold of *p* = 0.001 in the anatomical EC mask.

In the analysis of the hexagonal consistency effect, we employed a cross-validation procedure. We first estimated the mean orientation of the grid signal using one half of the dataset and then tested the hexagonal consistency on the remaining half. In training GLM, we modelled two 6-fold sinusoidal parametric regressors (sin(6θ) & cos(6θ)) separately for odd and even trials at the time of M2’s presentation and decision. We estimated the mean orientation φ for each participant within the *F*-test-based EC ROI, separately for both odd and even trials. The orientation φ of each voxel in the EC was calculated from the β coefficient of the sine and cosine regressors using the formula: 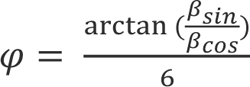. We calculated the mean orientation for each subject by averaging the estimated orientation across voxels in the EC ROI using the circular mean. Next, we incorporated the mean orientation into a test GLM to estimate the hexagonal consistency effect aligned with the mean orientation. The test GLM maintained the regressors of basic events and nuisance regressors but modelled the sinusoidal modulation with the regressor: cos (6[θ − φ]) instead of previous parametric modulations (sine & cosine) for odd and even trials. Here, θ refers to the direction of inferred trajectories in each trial, and φ represents the estimated mean orientation for each participant. Importantly, we applied the mean orientation estimated from the odd trials of training GLM to the even trials of the test GLM, and vice versa. This cross-validation procedure was also repeated for other periodicities, including 4-fold, 5-fold, 7-fold and 8-fold. Based on this cross-validation procedure, we categorized the inferred directions into 12 equal bins based on whether the directions aligned or misaligned with the estimated mean orientation. We modelled regressors for each bin to estimate their mean activities at the time of M2’s presentation to unfold the pattern of hexagonal activity.

The grid-like code can also be measured based on RSA^86^. The first step of this analysis involved the construction of a model for the representational dissimilarity matrix (RDM) to depict grid-like representation, as detailed in Bellmund *et al.* ^28^. The model RDM hypothesized that the similarity of neural activity is contingent upon the angular difference between inferred directions, an attribute stemming from the 6-fold periodicity of grid-like activity. The activity patterns showed a greater degree of similarity for angular differences within the 0-15° and 45-60° range, as opposed to the intermediate angular range of 15-45°. This pattern exhibits a cyclical nature, spanning from 0° to 360° with a recurring period of 60°. Consequently, the similarity in activity between any two directions can be divided into two distinct categories: similarity and dissimilarity. This classification enabled us to construct a model RDM for the grid representation, predicated on the similarity between pairs of inferred directions. In this RDM, we assigned a value of zero to similarity and one to dissimilarity.

Subsequently, we conducted a whole-brain searchlight analysis to compute the neural RDM of the inferred directions for each voxel. For this purpose, we modelled an independent regressor for each direction at the time of M2’s presentation in the GLM to estimate the mean activities of each inferred direction. Following this, for each voxel, we obtained its searchlight sphere that included its surrounding multi-voxels. The neural RDM of each sphere was estimated by calculating the correlation distance (1 − *r*) between the multivariate activity patterns associated with different directions. We then evaluated the representation similarity between the grid model RDM and the neural RDM, using Pearson’s correlation. The resulting correlation coefficients were re-mapped to the central voxel associated with each sphere and were subsequently transformed into z-scores for each brain voxel.

In Game2, we conducted a GLM to estimate a map alignment effect between two maps. The map alignment effect was quantified by the amplitude of hexagonal modulation in the knowledge-assimilated map aligned with the grid angle of the original map. Specifically, we used a similar cross-validation procedure as above to estimate the mean orientation φ_*g*ame1_ from the β coefficients of two 6-fold sinusoidal parametric regressors (sin(6θ) & sin(6θ)) from Game1’s data. We then incorporated this mean orientation into a sinusoidal parametric regressors (cos (6[θ_*g*ame2_ − φ_*g*ame1_]) in test GLM on the Game2’s data. Here, the θ_*g*ame2_ refers to the direction of inferred trajectories in Game2, which traverse pre-existing objects and new objects. The positions of new objects were hypothesized around the centre of the map and had a half-rank distance with pre-existing objects.

### Distance code

We adopted fMRI adaptation to measure the neural representation similarity between objects when inferring the relationship for novel pairs during mental navigation. This method evaluates neural representation similarity by measuring suppression or enhancement when two stimuli are presented in rapid succession^87^. We modelled the Euclidean distance between objects as parametric regressor during the presentation of M2. This Euclidean distance was calculated by the square root of the sum of the squared differences between their AP dimension and DP dimension, as following formula: 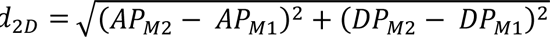. We also modelled the 1D rank difference in the AP and DP dimensions as parametric regressors in the GLMs for comparison with the 2D Euclidean distance representations. Specifically, we modelled both Euclidean distance and one-dimensional rank distance using the same timing and duration in separate GLMs and compared their parameter estimates.

### Value code

We modelled value code as a parametric modulation during the presentation time of the fight rule. During this time, subjects were required to make a decision by comparing the Attack Power (AP) of the attacking monster with the Defence Power (DP) of the defending monster to calculate the decision variables. In our first level of analysis, we modelled two potential decision variables: the absolute value of difference |*A*P − *D*P| and the difference of value (*A*P − *D*P). We also performed a GLM which included both the distance code and value code at the presentation time of M2 and decision. In this GLM, the distance code was modelled using Euclidean distance and the value code at decision-making was modelled using absolute value of difference |*A*P− *D*P|. The value code at the presentation time of M2 was defined by the mean of two possible decision values |*A*P_*M*1_ − *D*P_*M*2_| and |*A*P_*M*2_ − *D*P_*M*1_|, due to the rule being unknown at this stage.

### Conjunction analysis

After identifying the cognitive map representations in both games, we conducted a conjunction analysis to compare the representations of the original and knowledge-assimilated maps. This comparison was made for both the grid-like code and distance code at whole-brain level. We computed the area of overlap across the significant statistical maps of both Game1 and Game2 for grid-like code and distance code. It is important to note that all group maps that were included in the conjunction analysis had already been corrected for multiple comparisons to ensure the statistical significance of the areas of conjunction.

### Structural MRI data analysis

We reconstructed the cortical surfaces from T1-weighted MRI scans using FreeSurfer and obtained measurements of grey matter volume and thickness from individual surfaces^33,88^. The characteristics of the grey matter were then segmented into the 34 bilateral brain regions defined by the Desikan–Killiany parcellation^34^. For each brain region, we computed the Pearson correlation between its grey matter characteristics and both age and inference ability. Subsequently, these results were projected onto the corresponding surface parcellation. These correlation results were then projected onto the corresponding surface parcellation. We analysed the hippocampal volume derived from FreeSurfer’s brain segmentation using a similar procedure.

### Structural connectivity analysis

Fiber reconstruction was conducted using QSIprep 0.19.0, based on Nipype 1.8.6^67,80^. To achieve more accurate tissue boundary delineation, we employed a hybrid surface/volume segmentation approach, as proposed by Smith et al. ^89^. The outputs from FreeSurfer, along with previously defined ROIs, specifically, EC, mPFC, PCC, LOFC, and hippocampus, were registered to the outputs preprocessed by QSIPrep.

For the estimation of multi-tissue fiber response functions, we utilized the algorithm from Dhollander et al. ^90^. Fiber orientation distributions (FODs) in each voxel were then estimated using constrained spherical deconvolution (CSD), following the methodologies of ^91,92^, and employing an unsupervised multi-tissue method as described by Dhollander et al. ^93^. The FODs underwent intensity normalization using mtnormalize^94^.

The actual fiber reconstruction was executed in MRtrix3^94,95^. This process incorporated anatomically constrained tractography^96^ and spherical-deconvolution informed filtering of tracts^97^, thereby enhancing the biological plausibility and accuracy of the fiber connectivity measures.

To further analyze structural connectivity, we extracted and averaged FA values for the reconstructed tracts connecting pairs of ROIs. Initially, we fitted the diffusion tensor to the preprocessed diffusion-weighted images using iterated weighted least squares (IWLS) via MRtrix3. This process enabled the computation of fractional anisotropy (FA) across the brain^35,98^. Subsequently, the FA values were sampled from the FA image and averaged for each streamline. Then, for each connectome edge, the mean value was calculated by averaging the FA of all streamlines connecting the target pair, indexing the structural connectivity between ROIs.

To conduct a more detailed analysis of tracts connecting the EC and mPFC, we utilized labeled white-matter tracts from an independent atlas to identify the three canonical tracts: the cingulum bundle, uncinate fasciculus, and fornix^99^. These white-matter tracts connect the frontal lobe and temporal lobe via distinct pathways. In this analysis, we mapped streamlines from this atlas into probability density maps using MRtrix3. We applied a 10% threshold to these maps, creating three binary masks. These masks were then transformed to individual spaces, aligning with the FA images. Finally, we used these three masks to extract the mean FA values of the white-matter tracts.

### ROI analysis

We also examined the grid-like code, distance code, value code and structural measurements, as well as their relationships with age and inference performance at ROI level. We employed five ROIs that were independently defined outside of our study, including the EC, mPFC, hippocampus, LOFC and PCC. We used a bilaterally, anatomically defined EC ROI based on the Juelich Histological Atlas^29^. The hippocampus ROI was defined based on Harvard-Oxford subcortical structural atlas^34^. The mPFC ROI was defined based on a vmPFC mask from NeuroVault (https://identifiers.org/neurovault.image:18650). The LOFC ROI was defined based on the Desikan–Killiany parcellation^34^. The PCC ROI was defined based on the parcellation of default mode network from NeuroVault (https://identifiers.org/neurovault.image:319482)^100^.

In functional analysis, we averaged the parameter estimates or z-scores obtained from the first level analysis or RSA across all voxels within each ROI. We performed one sample t-tests to determine the significance of mean effect within each ROI. In correlation analysis, we standardized these values obtained from the first level analysis across subjects and examined their correlation with age and inference performance. We utilized a paired t-test to compare various scenarios: alignment versus misalignment, 2D distance versus 1D distance, and “*a*bs(*A*P − *D*P)” versus “*A*P − *D*P”. We also performed an analysis of variance to investigate whether a significant interaction effect exists between task stage (mental navigation and decision-making) and representational content (distance code and value code).

In structural analysis, the identical ROIs, defined in the same manner as in the fMRI analysis, were registered to the FreeSurfer’s fsaverage template. The individual grey matter volume and thickness were also mapped onto the fsaverage template. We extracted the corresponding grey matter characteristics from the ROIs. Then we calculated their correlations with age, inference performance, and the strength of task representations.

### Mediation analysis

We execute mediation analysis with a non-parametric bootstrap method^32,101^ to investigates how the influence of an independent variable X on a dependent variable Y is partially mediated by a third variable M. We derived statistical inference with a total of 10,000 iterations to generate confidence intervals (CI) and the two-sided p-value of indirect effect. The coefficients and confidence intervals of indirect effect were reported in Results.

We performed a series of path models to probe whether the task representations mediate the impact of development on inference performance. In these models, age was positioned as the independent variable (X), and inference performance served as the dependent variable (Y). We used the grid-like code within the EC, as well as the distance code and value code within the mPFC, as mediators. Prior to conducting the mediation, all metrics were standardized across subjects. For representations that showed significant indirect effects (grid-like code & distance code), we also performed a parallel mediation analysis that included both mediators. To test the relationship between grid-like code, distance code and inference performance, we performed a mediation analysis among the distance code within mPFC (X), grid-like code within EC (M), and map-based inference performance (Y). In the analysis of Game2, the map-based inference performance (Game1) was additionally incorporated into the paths of mediation analysis as covariate. Finally, we conducted mediation analyses by including age as a mediator, the structural features (grey matter volume & grey matter thickness) as an independent variable (X), and the task representations as the dependent variable (Y).

### Canonical correlation analysis (CCA)

We performed CCA to explore the relationship between cognitive map measures and intelligence scores. This analysis involved identifying a set of canonical variates—linear combinations of the original variables designed to maximize correlations between two sets of variables. To compute these canonical components, we employed the nonlinear iterative partial least squares (NIPALS) algorithm^38^. This algorithm iteratively ensures the orthogonality of successive variates by removing the variance of each identified component from the original data. After generating a series of weights for these linear combinations, we applied them to compute the canonical variates. We then calculated the loadings by correlating the canonical variates with the original measures, thereby illustrating the association between the original measures and the canonical variates. Finally, to determine the canonical correlation, we computed Pearson’s correlation between the canonical variates derived from cognitive map measures and intelligence scores. Statistical significance was assessed using 10,000 permutations, which involved randomizing the order of subjects within each measure set^39^. For each permutation, we reran the CCA to establish a valid null distribution, against which we compared the actual correlations. *P*-values were corrected for multiple comparisons across all modes estimated. The significance of each predictive coefficient was similarly estimated using this permutation approach.

## QUANTIFICATION AND STATISTICAL ANALYSIS

We presented the mean and standard error throughout both the figures and the manuscript. Statistical significance was reported using two-tailed *p*-values, except for the comparison between the alignment and misalignment conditions, where we used a one-tailed paired t-test based on our hypothesis that alignment would be greater. For one-sample *t*-tests, Cohen’s *d* effect sizes were calculated by dividing the mean of each group by its standard deviation. For paired *t*-tests, Cohen’s *d* was determined by dividing the mean difference between groups by the pooled standard deviation of both samples. To assess statistical differences between correlation coefficients, we applied Fisher’s *r*-to-*z* transformation and calculated corresponding *p*-values.

In the fMRI analysis, multiple comparison corrections were performed using the FMRIB Software Library (FSL) software^102^. Significant brain regions were reported with whole-brain family-wise error (FWE) correction at the cluster level (*p* < 0.05), with a cluster-forming voxel threshold of *p* < 0.001, except for the EC. Due to susceptibility to signal loss in the EC, a more lenient cluster-forming voxel threshold of *p* < 0.01 was used, as commonly reported in previous studies ^19,20^.

For the ROI analyses, we focused on predefined regions based on prior hypotheses: the grid-like code in the EC and the distance code in the mPFC. No statistical inferences were made beyond these specific ROIs, as our analyses were hypothesis-driven and limited to these regions, reducing the risk of multiple comparison issues.

We also conducted multiple correlations between measures of intelligence from standardised IQ tests and cognitive map-based inference from our study. To control for false positives, we used permutation tests within a CCA framework. This non-parametric approach, following Smith *et al.* ^39^, effectively adjusted for multiple comparisons, enhancing the reliability of our findings.

All analyses were conducted using Python and its libraries, including Nilearn, statsmodels, Pingouin, scikit-learn, and rsatoolbox, along with the FSL^101,103–106^. The circular heatmap was generated using ChiPlot (https://www.chiplot.online). Other quantitative and statistical methods are described above within the context of individual analyses in the method details section.

## SUPPLEMENTARY INFORMATION

**Figure S1.**
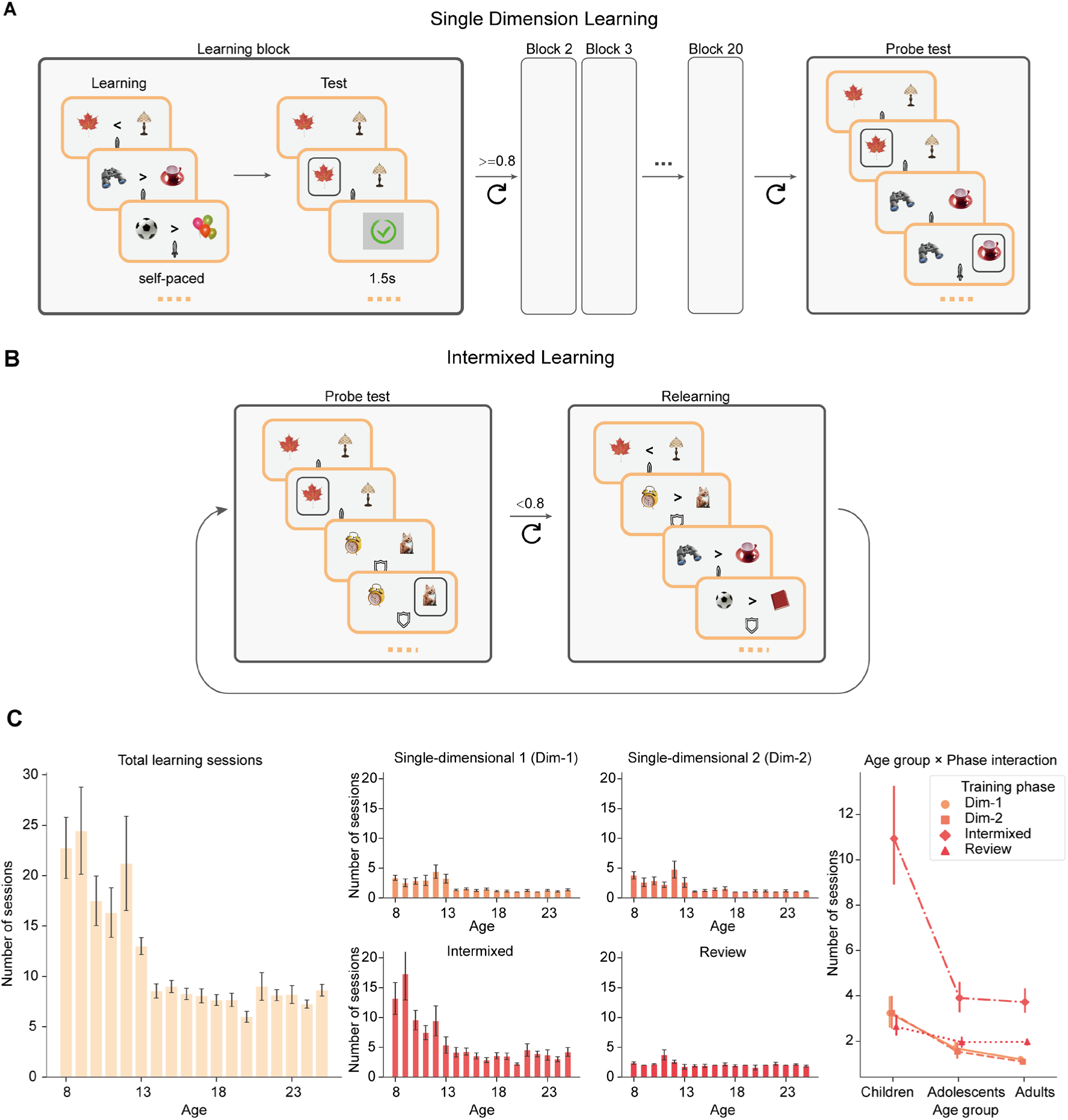
Training procedure and distribution of learning sessions across age groups, related to **Figure 1**, **Figure 7** and STAR Methods “Training procedure” section. (A) Single-dimension learning protocol. In each block, participants first completed a self-paced learning phase with explicit magnitude indicators (“>” or “<”), then proceeded to a testing phase without those indicators, receiving immediate feedback (1.5 s). They advanced to subsequent blocks upon achieving ≥ 80% accuracy and ultimately completed 20 blocks before a final probe test assessed overall retention. (B) Intermixed learning phase with an adaptive learning strategy. Participants alternated between probe tests (where they identified the higher-ranked stimulus without feedback for all pairs) and relearning sessions (where incorrectly answered pairs were re-taught with explicit magnitude indicators). The format remained self-paced. (C) Distribution of training sessions across age groups. The left panel shows the total number of sessions required to reach ≥ 80% accuracy for ages 8–25. The middle panel breaks down the sessions by single-dimension training, intermixed training, and review, highlighting that intermixed training had the greatest age-related variability. The right panel is a point plot illustrating the interaction between age groups (children: 8–12; adolescents: 13–17; adults: 18–25) and phase, indicating that children struggled most during the intermixed phase. Error bars represent s.e.m.

**Figure S2.**
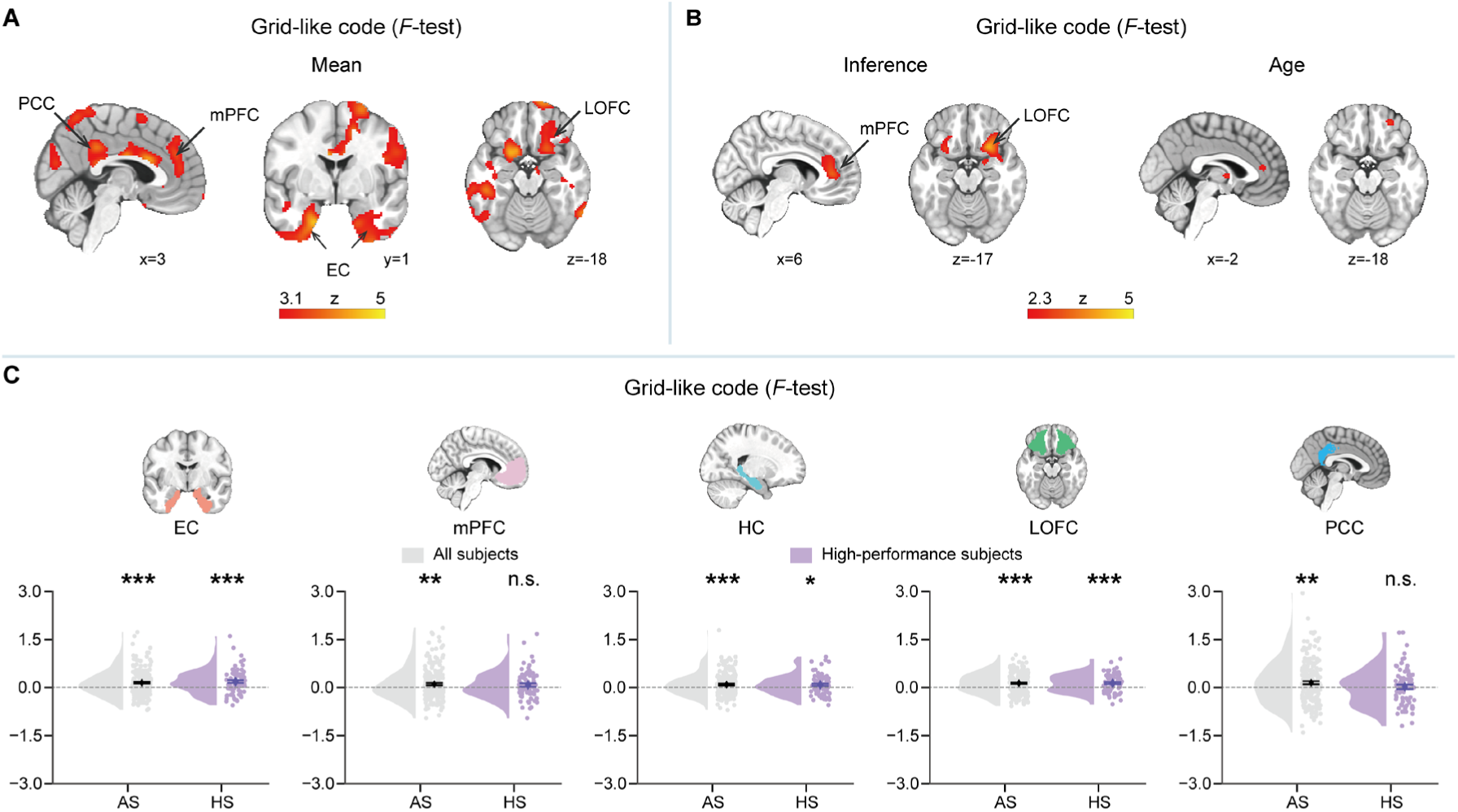
Grid-like code for inferred trajectories, related to. **Figure 2. (A)** Grid-like code was identified using the z-transform *F* statistic. Whole-brain analysis displayed grid-like code in brain regions, including EC, mPFC, LOFC, PCC and precentral gyrus (see also **Data S2**). The brain maps survived whole-brain FWE corrected at the cluster level (*p* < 0.05) with a cluster-forming voxel threshold of *p* < 0.01. **(B)** The whole-brain covariate maps show that the mPFC and LOFC exhibit positive correlations between the grid-like code and the ability of inferential reasoning. However, corresponding areas have only shown a feeble age effect. The covariate maps were thresholded at *p* < 0.01, uncorrected, for visualization purpose. **(C)** ROI analysis of grid-like code (*F*-test). The top row shows the ROIs, which were independently defined from previous studies. The second row shows the mean activity of grid-like code within the corresponding ROIs for all subjects (AS, grey) and high-performance subjects (HS, purple). Significance is denoted as: *\* p* < 0.05. ** *p* < 0.01. *** *p* < 0.001. n.s. non-significant.

**Figure S3.**
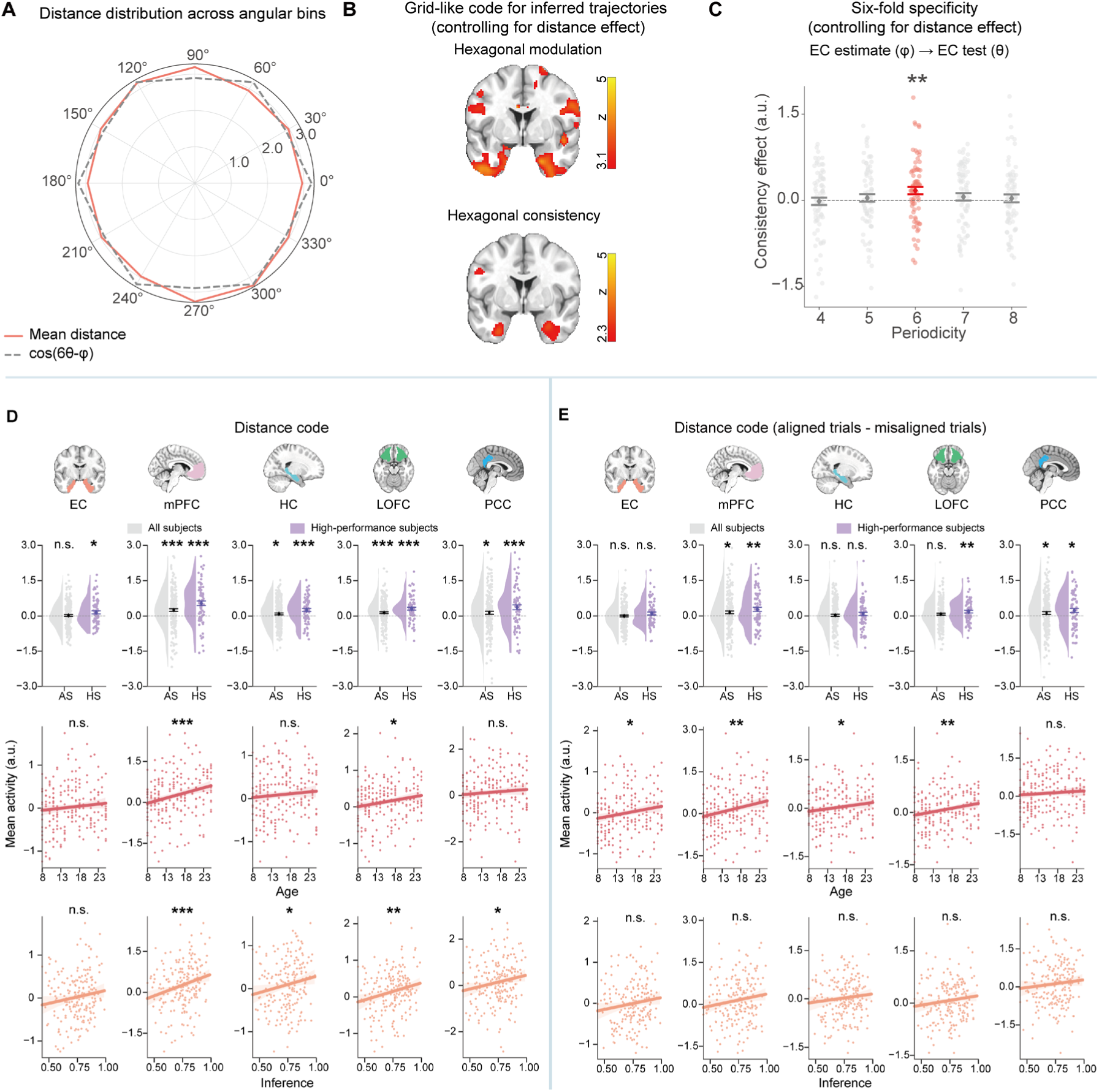
Analysis related to distance effect in Game1, related to Figure 2 and Figure 3. (A) A polar plot shows the distance distribution across 12 angular bins (red line) compared to the theoretical hexagonal modulation (grey line). The amplitude of the hexagonal modulation was normalised to the mean and standard deviation of the Euclidean distance distribution. (B) Grid-like code for inferred trajectories persisted in the EC after controlling for Euclidean distance effect. (C) The grid modulation aligned with the mean orientation remained consistently specific to a six-fold pattern after controlling for Euclidean distance, and was not observed for other control periodicities. (D) ROI analysis of distance code. The top row shows the ROIs, which were independently defined from previous studies. The second row shows the mean activity of distance code within the corresponding ROIs for all subjects (AS, grey) and high-performance subjects (HS, purple). The third row shows the correlations between age and the distance code within the ROIs. The fourth row shows the correlations between inference performance and the distance code within the ROIs. **(E)** ROI analysis for grid-like modulation on distance code. This involves contrasting the distance code of aligned and misaligned trials within ROIs. The trials were labelled based on whether the inferred direction aligned or misaligned with the estimated mean orientation from EC. The layout mirrors panel A. Significance is denoted as: *\* p* < 0.05. ** *p* < 0.01. *** *p* < 0.001. n.s. non-significant. Error bars denote s.e.m.

**Figure S4.**
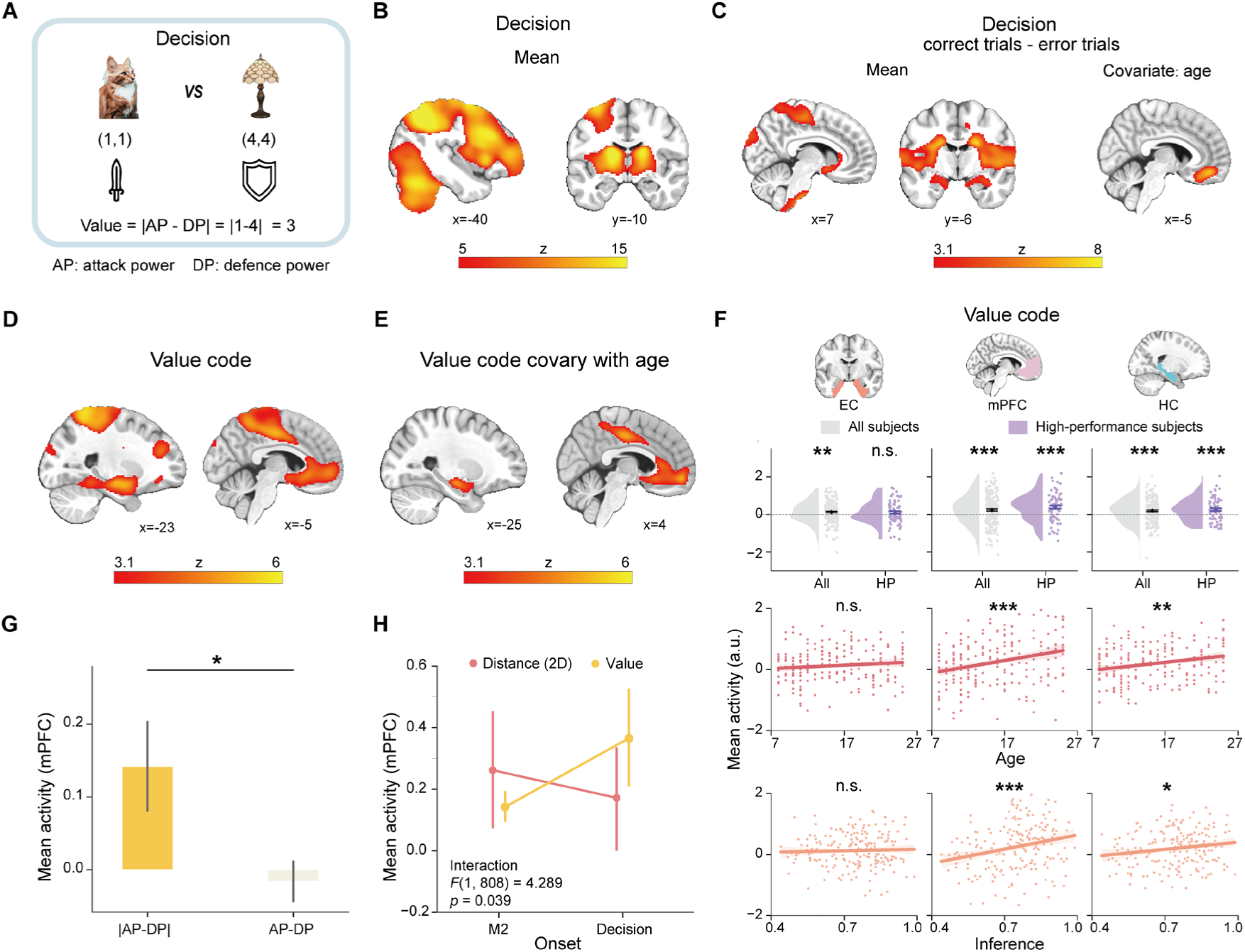
Value code at decision time, related to Figure 3 and STAR Methods “Value code” section. (A) A trial demonstration illustrates how the value code was calculated. We compared the AP of attacking monsters to the DP of defending monsters and calculated the absolute difference between AP and DP as the value code. (B) Whole-brain analysis shows the mean activity during decision time with significant activation in the cerebellum, visual cortex, striatum, primary motor cortex and dorsolateral prefrontal cortex. (C) Contrast brain map between correct versus error response trials, showing stronger activation in ventral striatum, amygdala and primary motor cortex. The covariate results (right panel) show that the difference between correct trials and error trials increase with age in the mPFC. (D) Whole-brain parametric analysis with value code found significant involvement of hippocampus, mPFC, and primary motor cortex. (E) Whole-brain covariate analysis with age for value code. (F) ROI analysis of value code. The top row shows the ROIs. The second row shows the mean activity of value code within the corresponding ROIs for all subjects (AS, grey) and high-performance subjects (HS, purple). The third row shows the correlations between age and the value code within the ROIs. The fourth row shows the correlations with inference performance. (G) ROI analysis shows that the value code in mPFC represents the absolute difference between AP and DP, rather than the difference between AP and DP. (H) A shift in neural representation within the mPFC, from representing the distance code at M2 stage to encoding the value code during the decision time (see also Data S8). All brain maps survived whole-brain FWE corrected at the cluster level (*p* < 0.05) with a cluster-forming voxel threshold of *p* < 0.001. *\* p* < 0.05. ** *p* < 0.01. *** *p* < 0.001. n.s. non-significant.

**Figure S5.**
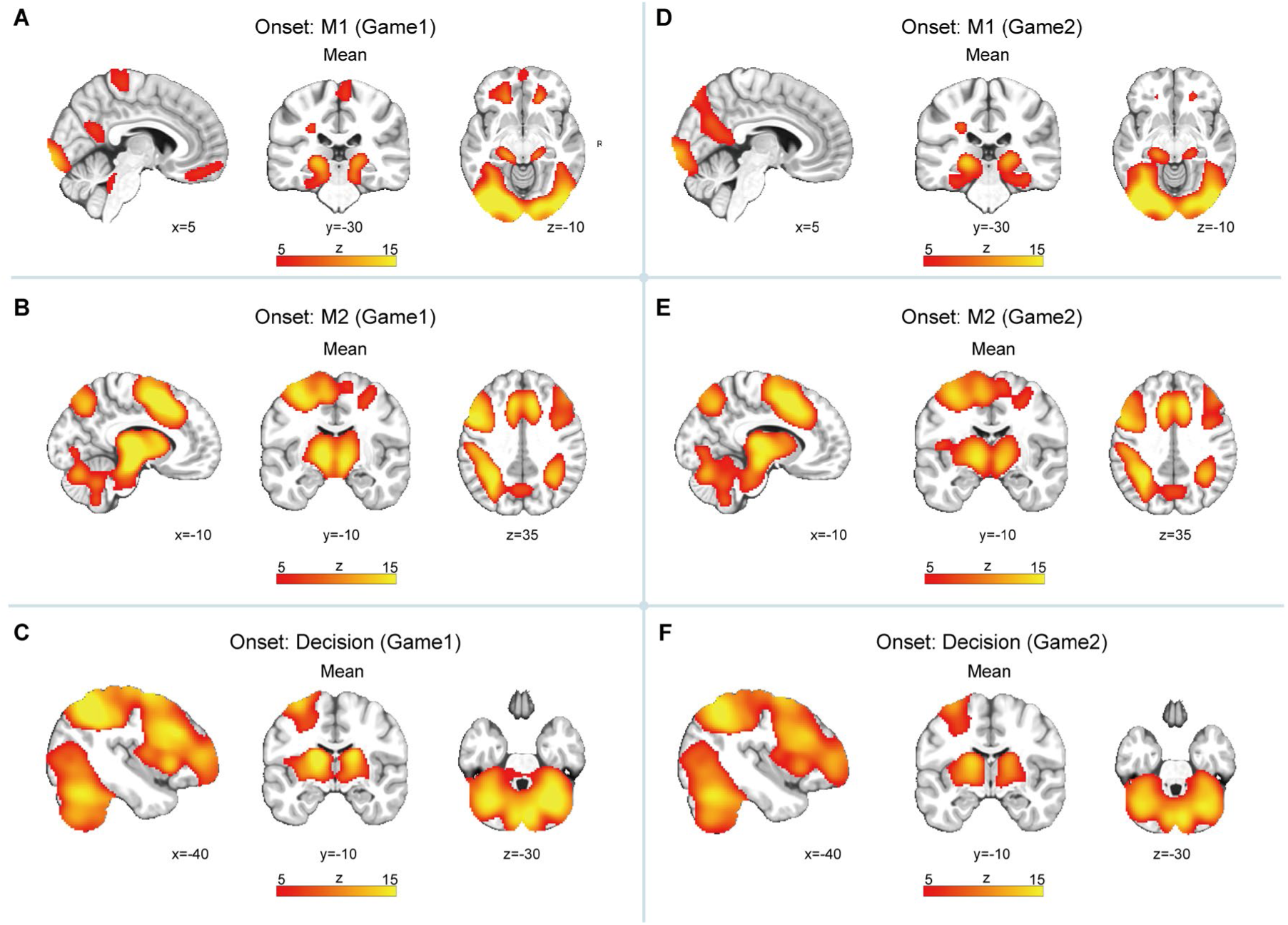
Mean activity during different phases of an inference trial in both Game1 and Game2, related to STAR Methods “Functional MRI data analysis” section. (A) At the onset of M1 (Game1), whole-brain analysis revealed significant activations in visual cortex, posterior cingulate cortex, and hippocampus. (B) At the onset of M2 (Game1), significant activations were found in anterior cingulate cortex, visual cortex, and superior parietal lobule. (C) Upon decision time (Game1), we observed significant activation in the cerebellum, visual cortex, striatum, primary motor cortex and dorsolateral prefrontal cortex. (D-F) In Game2, whole-brain analysis revealed similar activation patterns across each phase. All brain maps were whole-brain FWE corrected at the cluster level (*p* < 0.05) with a cluster-forming voxel threshold of *p* < 0.001. For visualization purpose, whole-brain maps presented here were thresholded at z-value > 5.

**Figure S6.**
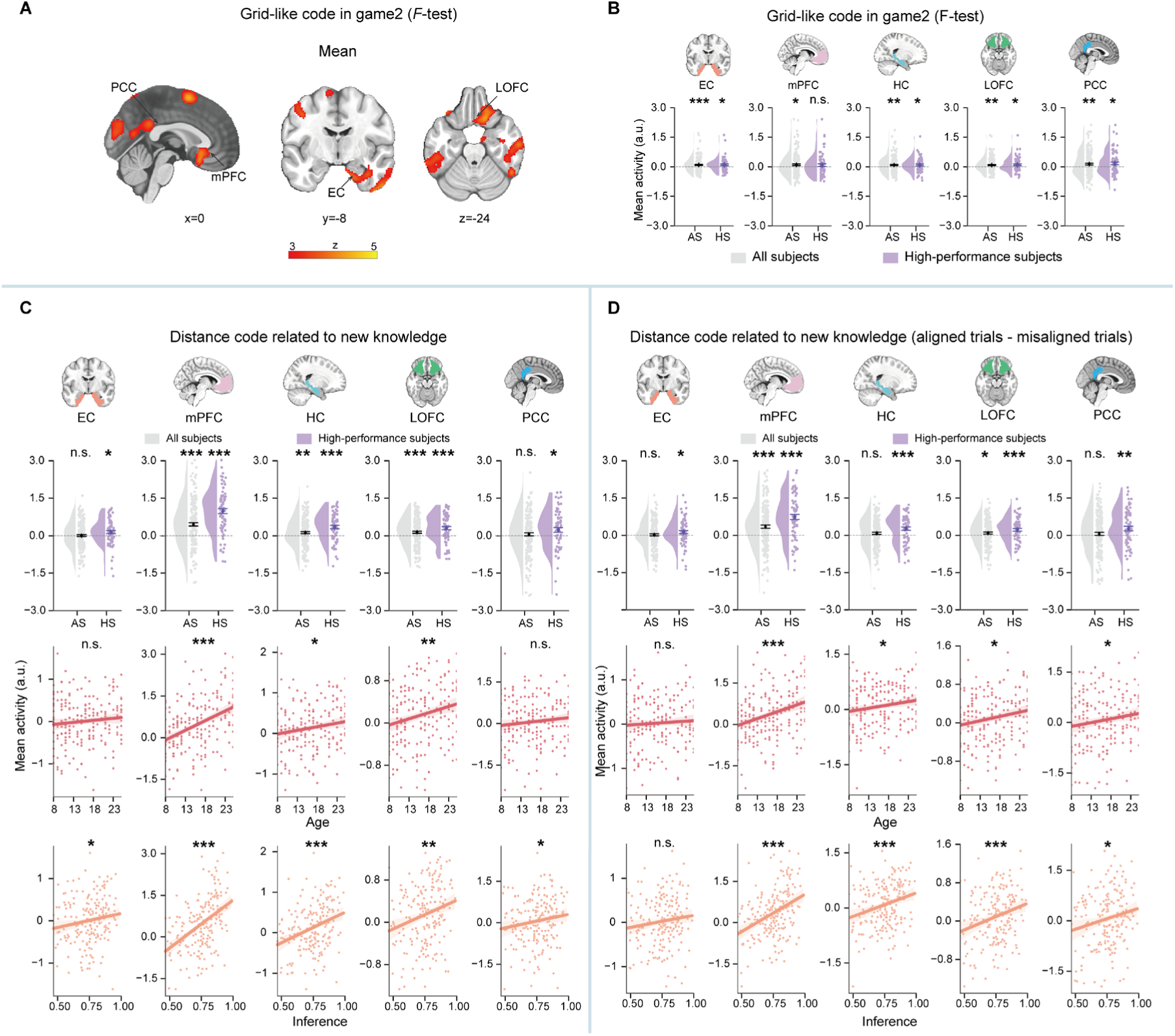
Grid-like code and distance code for inferred trajectories in Game2, related to. **Figure 4 and Figure 5. (A)** Whole-brain results are displayed, showing the clusters exhibiting grid-like code (*F*-test) of Game2. The clusters include the EC, OFC, PCC (see also **Data S11**). This map survived whole-brain FWE correction at the cluster level (*p* < 0.05) with a cluster-forming voxel threshold of *p* < 0.01. **(B)** ROI results of grid-like code (*F*-test) in Game2. The top row shows the ROIs, which were independently defined from previous studies. The second row shows the mean activity of grid-like code within the corresponding ROIs for all subjects (AS, grey) and high-performance subjects (HS, purple). Significance is denoted as: *\* p* < 0.05. ** *p* < 0.01. *** *p* < 0.001. n.s. non-significant. **(C)**, ROI analysis of distance code related to new knowledge (see also **Data S12**). The top row shows the ROIs, which were independently defined from previous studies. The second row shows the mean activity of distance code within the corresponding ROIs for all subjects (AS, grey) and high-performance subjects (HS, purple). The third row shows the correlations between the distance code within ROI and age (red). The fourth row shows the correlations with inference performance of Game2 (orange). **(D)** ROI analysis for grid-like modulation on distance code related to new knowledge. The layout mirrors panel A. Significance is denoted as: *\* p* < 0.05. ** *p* < 0.01. *** *p* < 0.001. n.s. non-significant.

**Figure S7.**
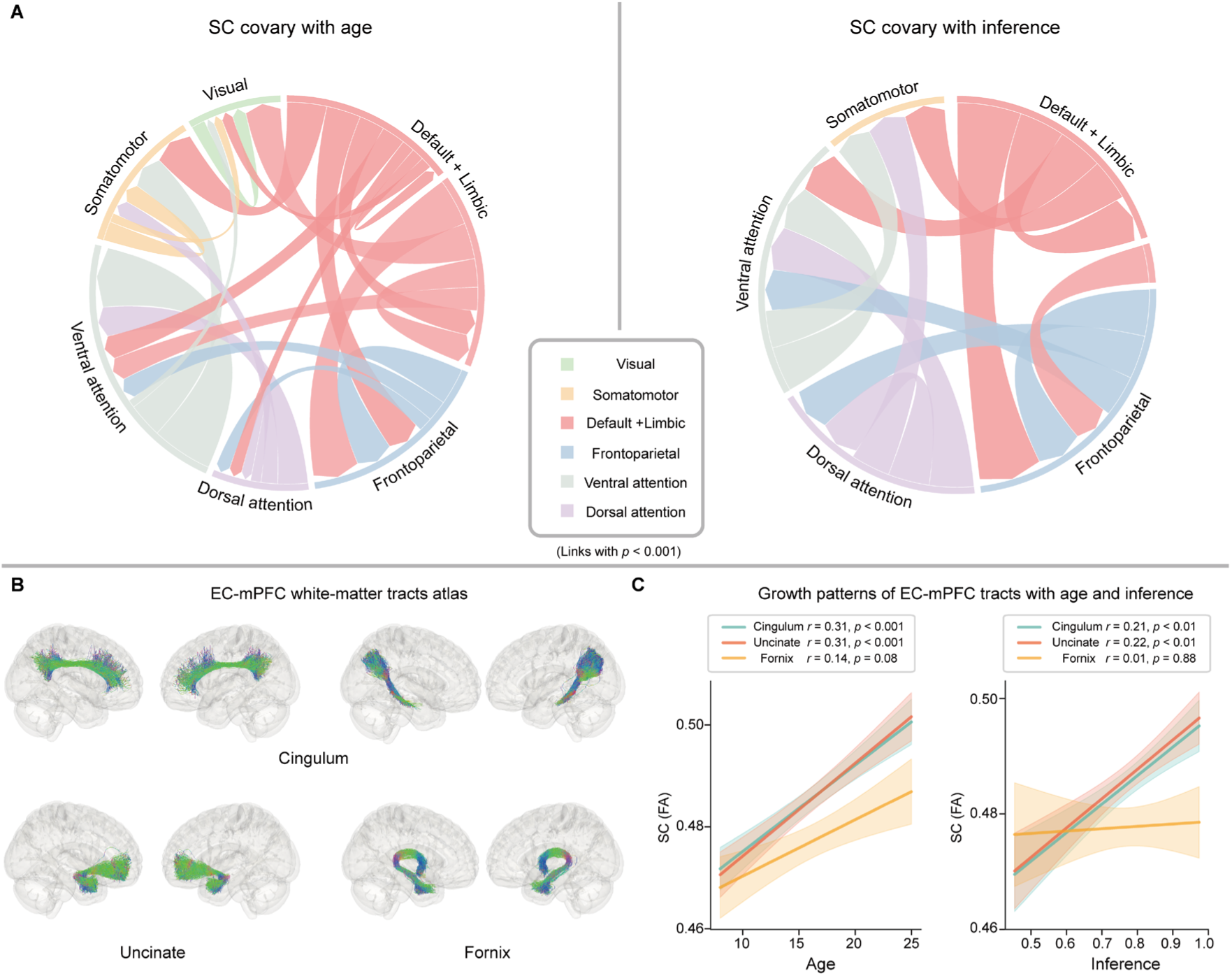
Changes in structural connectivity across brain networks related to age and inferential performance, related to. **Figure 6. (A)** The chord diagram illustrates changes in structural connectivity (measured by FA) associated with age and inference performance, based on 7 brain networks identified by Yeo et al. (2011). Each link represents the correlation between structural connectivity and either age or inferential performance, with the line thickness indicating the relative strength of structural connectivity among the networks. Notably, the default mode and limbic networks, which include the brain regions of interest (ROI), i.e., MTL-mPFC, show a stronger positive correlation with inference performance compared to the visual and somatomotor networks. Only links demonstrating significant correlations (*p* < 0.001) are displayed. **(B)** Three canonical white-matter tracts connecting the MTL and PFC are labelled using an independent dataset by Maffei *et al*. ^99^. These tracts, namely the cingulum, uncinate, and fornix, are used as masks for subsequent analyses. **(C)** All white-matter tracts connecting the EC and mPFC exhibited a positive trend with age. However, significant correlations with inference performance were observed only in the changes of the cingulum and uncinate.

**Table S1.** Demographic information of subjects, related to Figure 1 and Figure 6.

| Characteristics | Subjects |  |
| --- | --- | --- |
|  | Age 8-16 | Age 17-25 |
| **N** | 116 | 87 |
| **Age (SD) in years** | 11.66 (2.55) | 20.94 (2.83) |
| **Female (%)** | 48.28 | 67.82 |
| **Mean SPM score (SD)** | 51.91 (5.88) | -- |
| **WISC-IV** |  |  |
| **Coding** | 67.85 (22.44) | -- |
| **Digit Span** | 28.27 (5.12) | -- |
| **Verbal Similarities** | 33.6 (3.93) | -- |
| **Verbal Comprehension** | 29.32 (6.02) | -- |
| **Perceptual Matrix** | 28.52 (3.92) | -- |
| **Perceptual Block** | 53.24 (10.46) | -- |

SPM, Raven's Standard Progressive Matrices; WISC-IV, Wechsler Intelligence Scale for Children - Fourth Edition. --, not applicable.

## Supplementary information

### Data S1. Exclusion of potential age-related confounding factors

To ensure the validity of our results, we conducted extensive control analyses to rule out age-related confounding factors.

### 1.1 Age distribution

We examined the age distribution of our sample to ensure consistency between behavioural and fMRI analyses, comprising 203 participants for Game1 and 193 for Game2. Chi-square goodness-of-fit tests were conducted to determine whether the age distribution differed from uniformity for either Game1 or Game2. These tests revealed no significant deviation (Game1: *χ²*(18) = 18.59, *p* = 0.35; Game2: *χ²*(18) = 18.05, *p* = 0.39; **Data S1.1**), indicating that our findings were not biased by the sample’s age distribution.

**Data S1.1.**
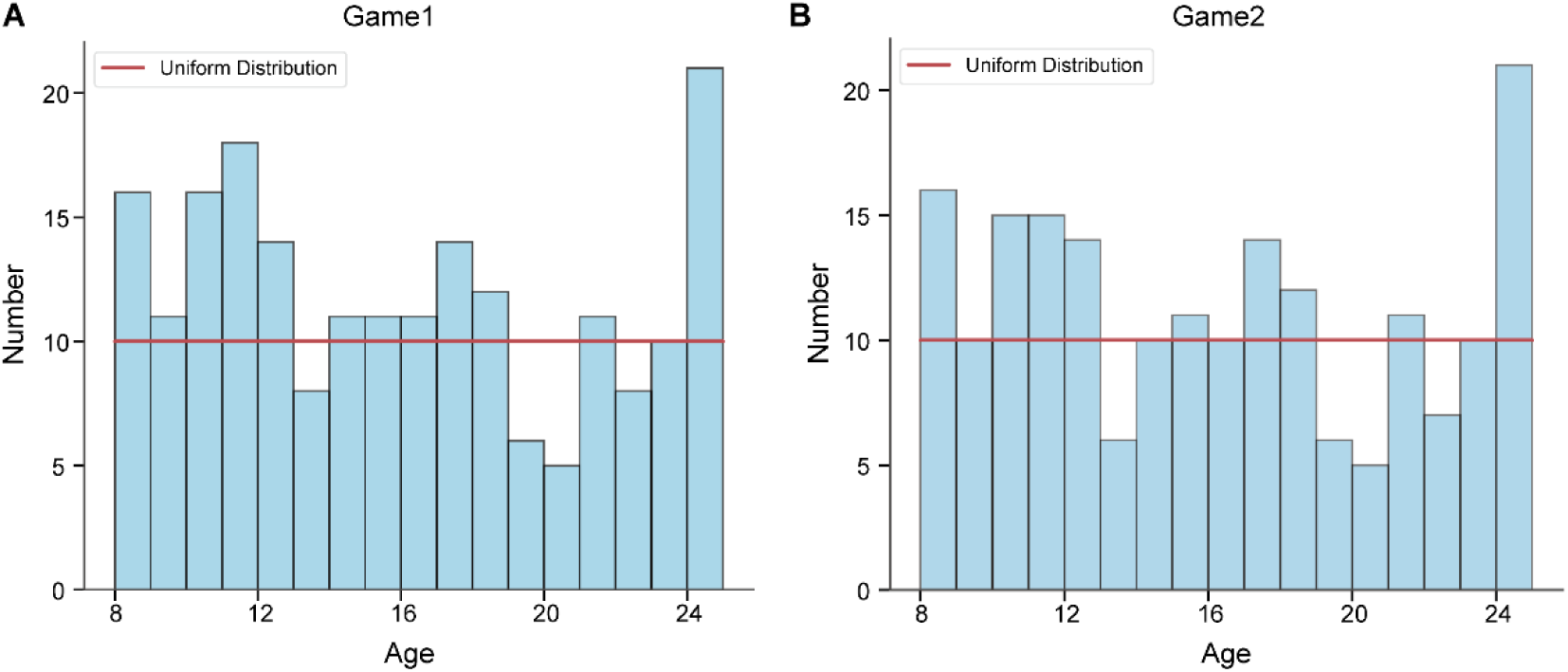
Age distribution of participants. (A) The age distribution of Game1. (B) The age distribution of Game2. Each bar represents a one-year age range, with bar height indicating the number of participants. The red line depicts the expected uniform distribution.

### 1.2 Daily effect

We addressed potential daily biases, such as a systematic relationship between participants’ ages and their scanning times, by randomly assigning them to three time blocks throughout the day: 10:00-12:00, 13:00-15:00, and 15:00-17:00. To examine the relationship between age and experiment timing, we categorised participants into three age groups: children (8–12 years), adolescents (13–17 years), and adults (18–25 years). A chi-square test of independence revealed no significant association (*χ²*(4) = 4.55, *p* = 0.34; **Data S1.2**), indicating that participants were evenly distributed across different times of day.

**Data S1.2.**
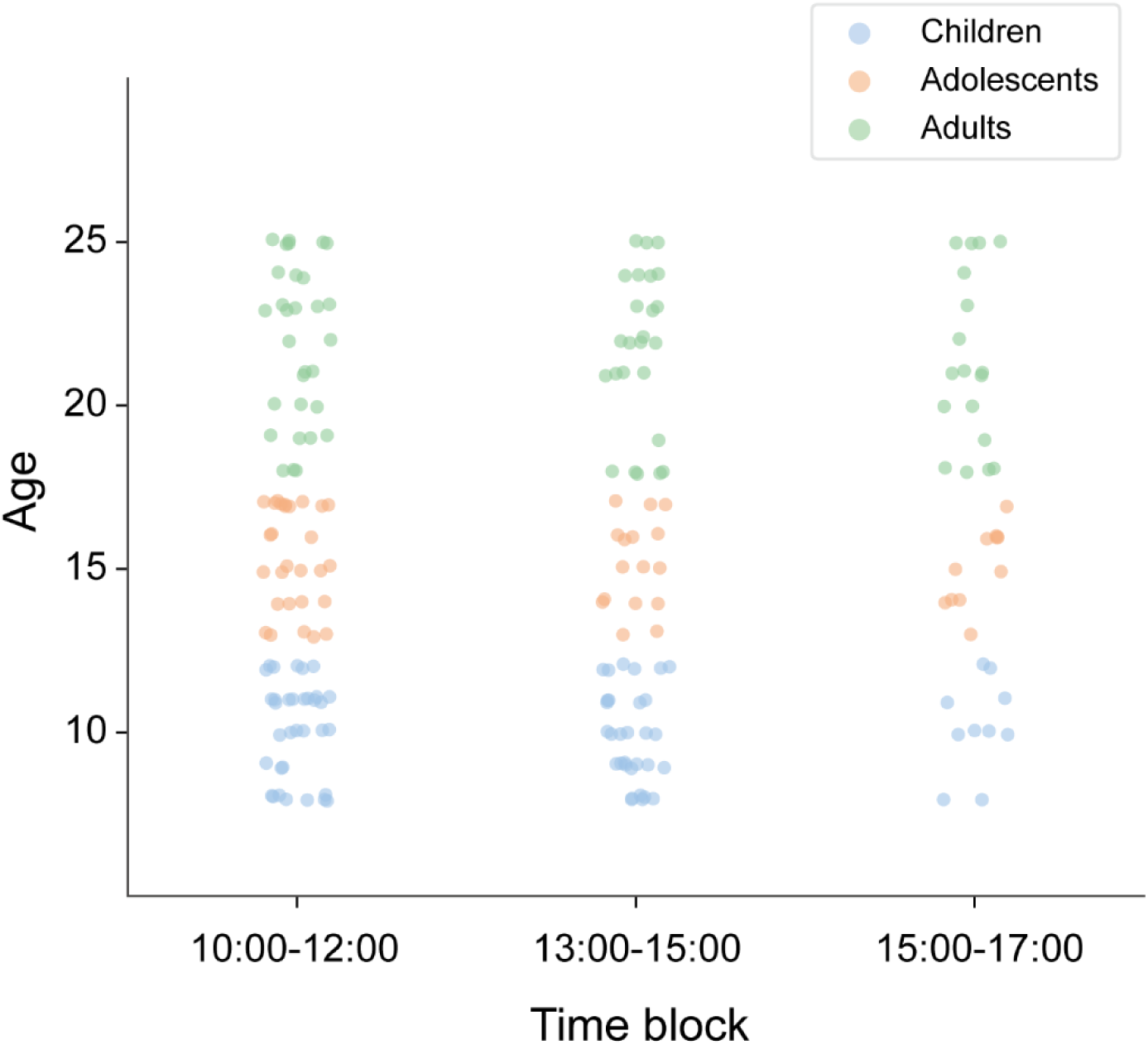
Distribution of participants by age and time block. The scatter plot shows the age distribution of participants across three daily time blocks: block1 (10:00-12:00), block2 (13:00-15:00), and block3 (15:00-17:00). Age groups are colour-coded: children (blue, 8–12 years), adolescents (orange, 13–17 years), and adults (green, 18–25 years).

### 1.3 Sleep effect

Prior to the experiment, we instructed all participants to have adequate rest the night before their MRI scan. We assessed their sleep quality using the Pittsburgh Sleep Quality Index^1^. No participant exhibited severe sleep problems, as indicated by a cutoff score of 10 on the sleep index.

### 1.4 Socioeconomic status

Due to privacy concerns, only 106 participants provided their socioeconomic status (SES). Analysis of this subset revealed a negative correlation between SES and age (Spearman correlation: *r* = −0.23, *p* = 0.03, N = 106; **Data S1.4**), indicating that older participants in our study tended to have lower SES.

To ensure SES did not confound our results, we included it as a control variable in our mediation analysis with grid-like code development. The findings showed that the observed increase in inferential ability remained significantly mediated by the development of the grid-like code (*β*_ab_ = 0.06, *p* = 0.02, CI [0.01, 015], N=106). This means that even after accounting for SES, our main results held true.

These findings indicate that the age-related improvements in inference performance are robust and are not significantly affected by the decrease in SES among older participants. Therefore, SES does not appear to be a significant confounding factor in our study of the cellular mechanisms underlying cognitive development with age.

**Data S1.4.**
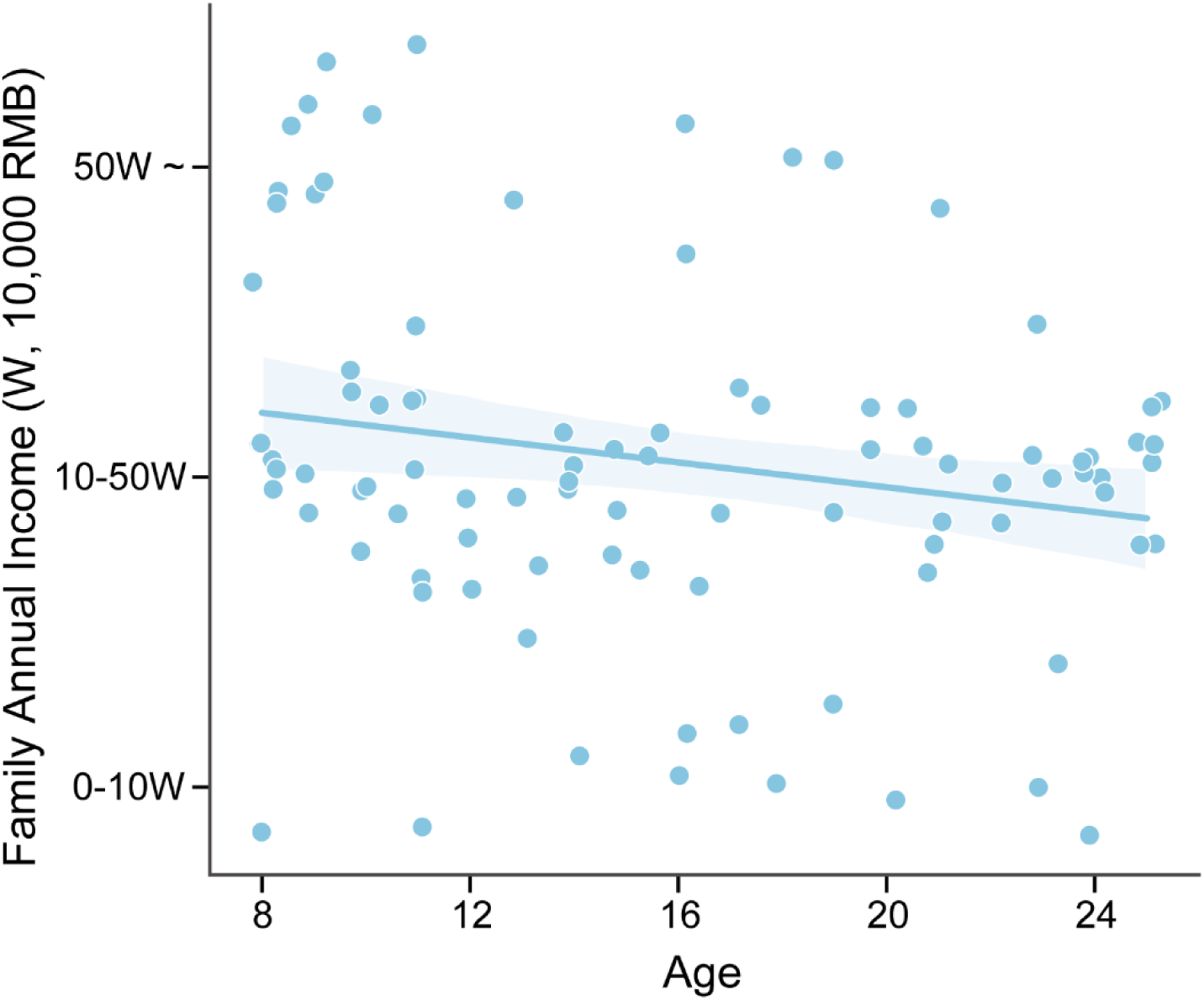
Age-dependent distribution of SES. The scatter plot illustrates the relationship between age (x-axis, years) and annual family income (y-axis, in W, where 1W = 10,000 RMB). Income is categorised into three groups: 0-10W, 10-50W, and >50W. Each blue dot represents an individual participant. The solid blue line indicates the best-fit linear regression, with the shaded area representing the 95% confidence interval.

### 1.5 Memory difference and forgetting effect

Our experimental protocol was designed to minimise memory differences and focus on age-related improvements in inferential reasoning abilities:

#### Pre-Scanning Memory Standard

Participants were invited for MRI scanning only after consistently meeting a predefined standard in recalling local pairwise relationships (above 80%) during the training procedure (**Figure 1C**). This controlled for individual memory differences and forgetting effects.

#### Timing of Knowledge Tests

Knowledge tests were conducted shortly before the formal fMRI scanning session to confirm that participants retained the memories necessary for the inference task during the fMRI experiments.

#### Adjusting for Memory Performance

We found a positive correlation between age and memory performance across subjects (*r* = 0.19, *p* = 0.01, N = 203, **Figure 1C**). To account for this, we employed partial correlation, as well as generalized linear model (GLM) covariate analysis, adjusting for premise pair memory. These methods are detailed in the ‘Behavioural Analysis’ in the Methods section. After adjusting for memory differences, the age-related increase in inference performance remained significant (*r* = 0.54, *p* < 0.001, N = 203; **Figure 1D**). A similar pattern was observed in nonlocal inference ability (*r* = 0.56, *p* < 0.001, adjusted for memory: *r* = 0.54, *p* < 0.001, N = 193, **Figure 1E**).

### 1.6 False memory influence of premise pairs

To confirm that the age-related increase in inference ability was beyond memory capacity, we excluded inference trials containing objects that were incorrectly identified in the pre-scan memory test following ‘premise learning’. The results of this refined analysis, corroborate our main findings (see **Figure 1**). Notably, we still observed a strong age effect in map-based inference (*r* = 0.51, *p* < 0.001; **Data S1.6**). This persistent age effect, despite the more stringent data inclusion criteria, highlights the robustness of our findings.

**Data S1.6.**
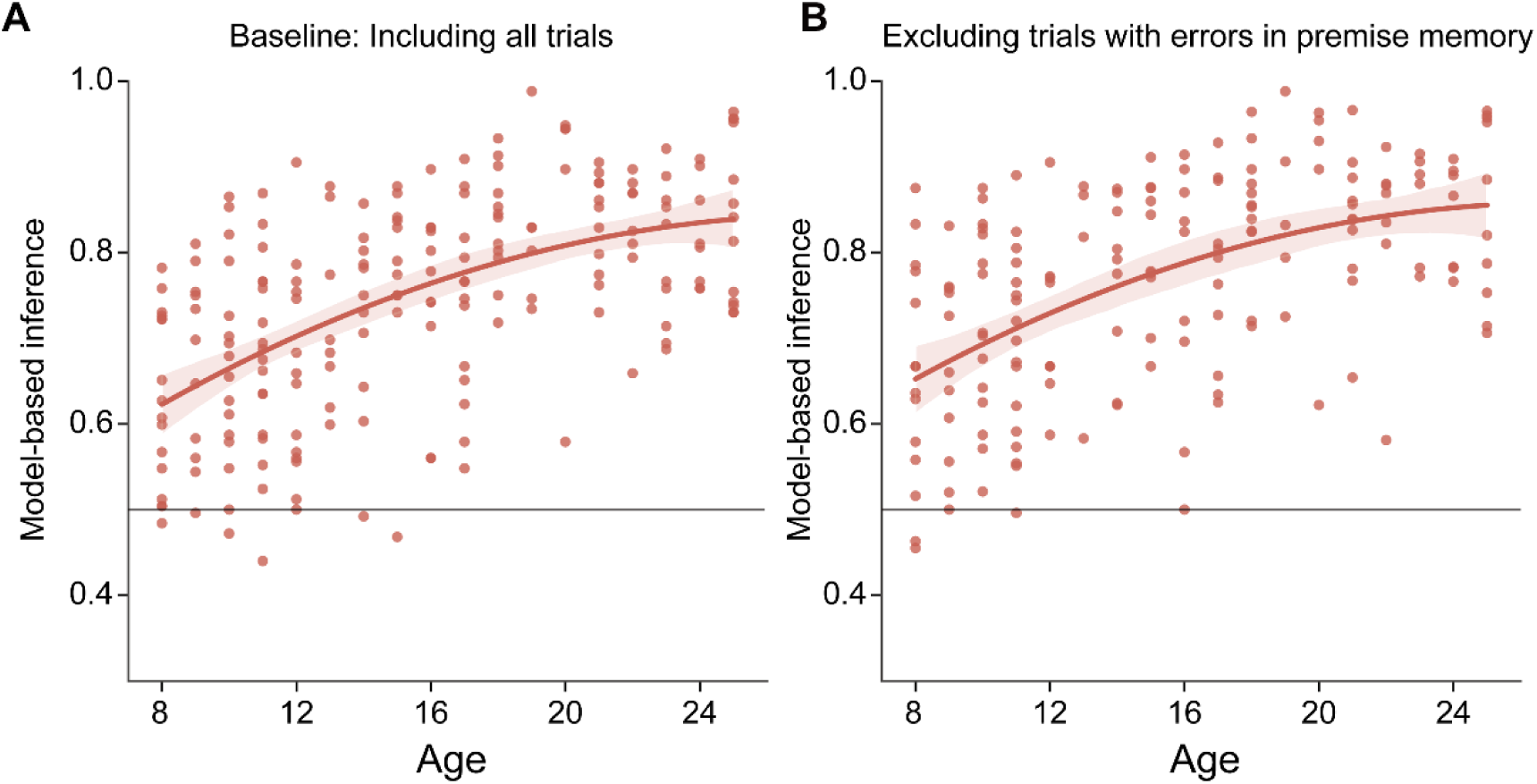
Developmental trajectories of map-based inference performance. **(A)** Baseline analysis including all trials, as shown in Figure 1D. **(B)** Analysis excluding trials where objects were incorrectly identified in the memory test. Each circle represents data from an individual participant. Error bars denote the standard error of the mean (s.e.m.). The black line indicates the change level of the task. Both analyses showed significant results for age-related inference performance.

### 1.7 Processing speed

There is a potential concern that children may not have sufficient time to make decisions compared to adults. However, both our task design and behavioural results demonstrate that this is not the case.

Our task design encourages participants to consider the two objects before making a decision. This is achieved by presenting the two stimuli sequentially, with the decision rule appearing only after both stimuli have been shown. The decision time was set to 3 seconds based on pilot results involving both young children and adults, ensuring that all participants have enough time to respond.

Behavioural results from the fMRI experiment further support that the response time was sufficient. Analysis of the mean reaction times (RT) for each participant showed that all subjects responded well below the 3-second limit (mean RT: 1.16 ± 0.02 seconds, *t* = −85.63, *p* < 0.001), including young children below 12 years old. The correlation between age and RT was also not significant (*p* > 0.05, **Data S1.7**). Therefore, processing speed is unlikely to be a significant confounding factor in our study.

**Data S1.7.**
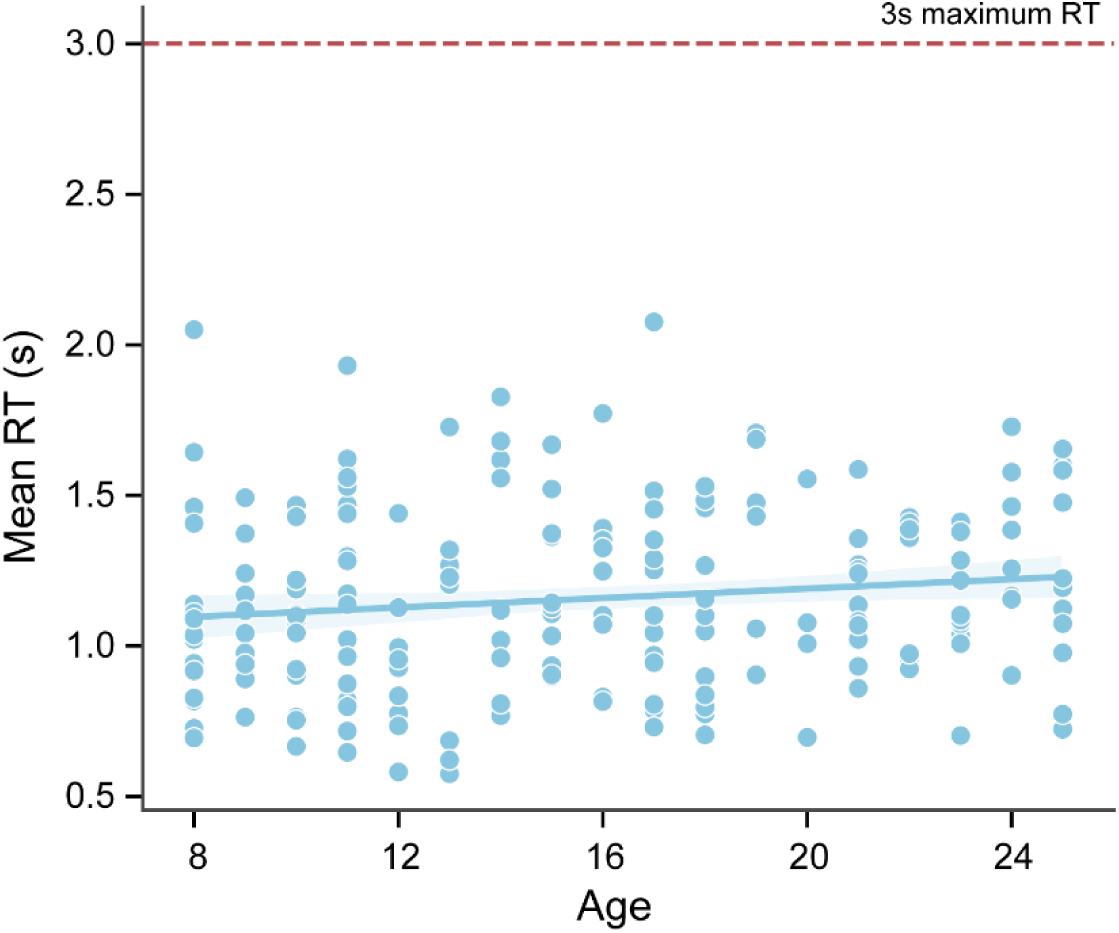
Mean reaction time (RT) of participants across age. The scatter plot shows the mean RT of participants in the inference task. The red line indicates the maximum RT allowed in the decision-making phase. Each circle represents an individual participant. All participants’ mean RTs are well below the maximum line.

### 1.8 Fatigue effect of 8-12 years old children

To assess potential fatigue effects in children during task performance, we provided additional evidence of performance accuracy as the task progressed. For children aged 8 to 12, we divided the inference trials into six runs for each game. Using repeated measures ANOVA, we assessed whether there were significant differences in accuracy between runs. Our analysis revealed no significant fatigue effects across the experiment for children (Game1: *F*(5, 370) = 1.17, *p* = 0.32; Game2: *F*(5, 345) = 0.70, *p* = 0.63; **Data S1.8**).

We should also note that our task design was structured to mitigate potential fatigue effects. The fMRI experiments lasted approximately one hour, with premise learning conducted outside the scanner. Participants were given sufficient time to rest during the experiment. They were allowed a 10-minute rest period after Run 3 of Game1 and a 15–30 minute rest period after Game1, or until they felt ready to continue.

**Data S1.8.**
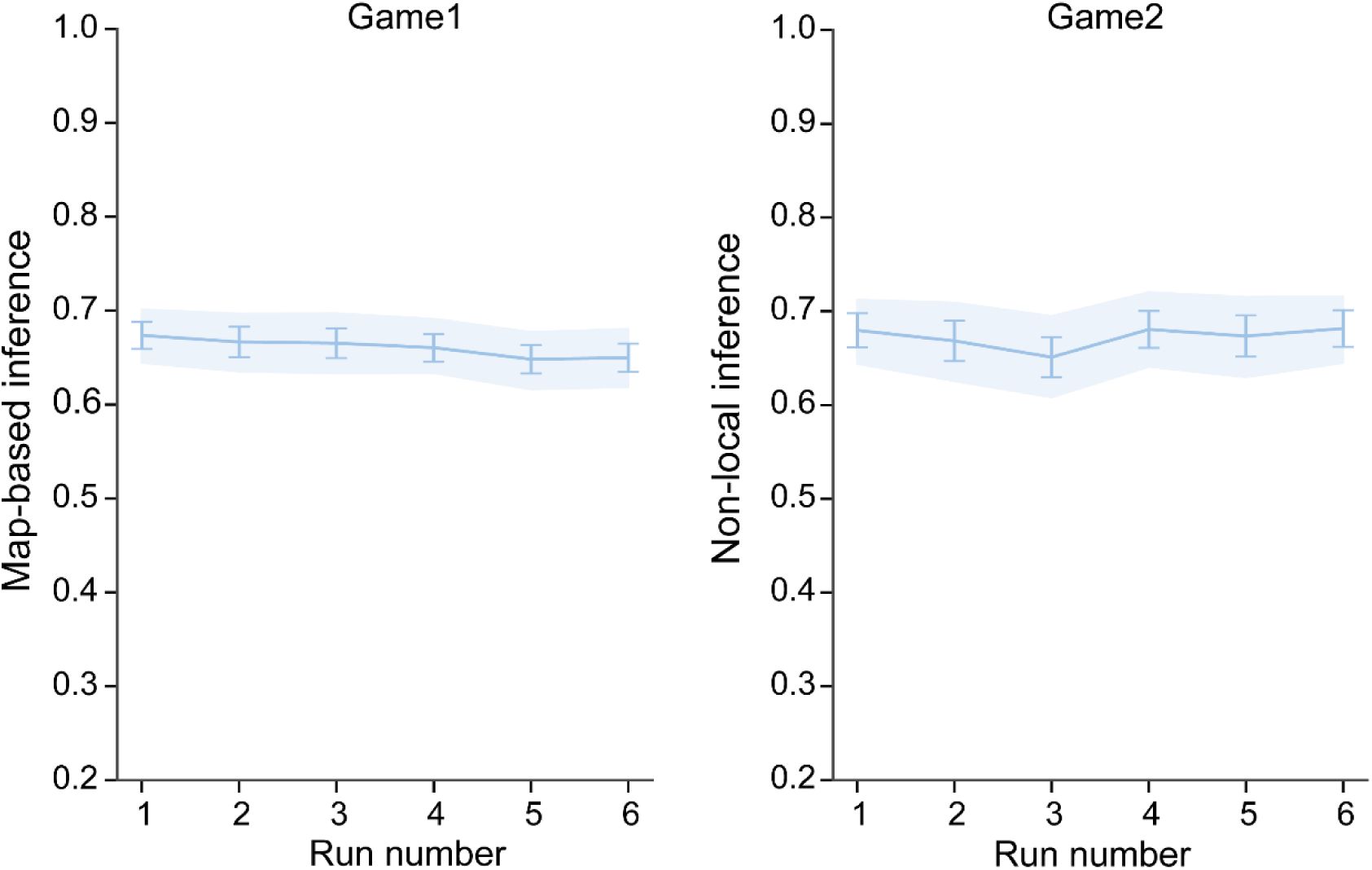
Inference performance as a function of task runs. Behavioural accuracy was shown for each run in Game1 and Game2. For each game, inference trials were divided into six consecutive runs. Each point represents the mean accuracy in a specific run, with error bars indicating the standard error of the mean.

### 1.9 Task engagement of 8-12 years old children

We also examined the task engagement of younger children (age 8-12 years old) to ensure that the observed age-related differences were not simply due to younger children being unable to perform the task.

Our analysis demonstrated that children aged 8–12 years actively participated in our task, achieving above-chance performance in all tasks. Specifically, 69 out of 75 children had inference accuracy above the 50% chance level in Game 1, and 66 out of 70 in Game 2. Statistical tests confirmed that children aged 8–12 years performed significantly above chance on all tasks (map-based inference: *t* = 12.46, *p* < 0.001; non-local inference: *t* = 11.01, *p* < 0.001).

Furthermore, ROI analysis in the entorhinal cortex (EC) revealed significant activation for grid-like codes among children aged 8–12 years (*t* = 3.52, *p* = 0.001). Also, as shown in **Data S1.8**, there was no evidence of fatigue effects acting as a confounding factor for children in this age group. Together, these findings reject the notion that children merely contributed noise to the study. This is not true at either the behavioural or neural level.

### 1.10 Nonlinear age-related hypothesis

We investigated whether a nonlinear relationship better describes our data compared to a linear one. We compared linear and quadratic models for map-based inference in Game1. The results indicated that the quadratic term has minimal influence and does not significantly improve the model (*β*_quadratic_ = −0.0006, *t* = −1.89, *p* = 0.06). Additionally, the Bayesian Information Criterion (BIC) for the linear model (−914.91) was lower than that of the quadratic model (−913.21), suggesting a better fit when accounting for model complexity. These findings were also replicated in non-local inference task in Game2 (*β*_quadratic_ = −0.0005, *t* = −1.36, *p* = 0.18; BIC for linear model: −797.83; BIC for quadratic model: −794.43; **Data S1.10**). These findings suggest that a linear relationship adequately describes our data, with no substantial evidence supporting a nonlinear relationship.

**Data S1.10.**
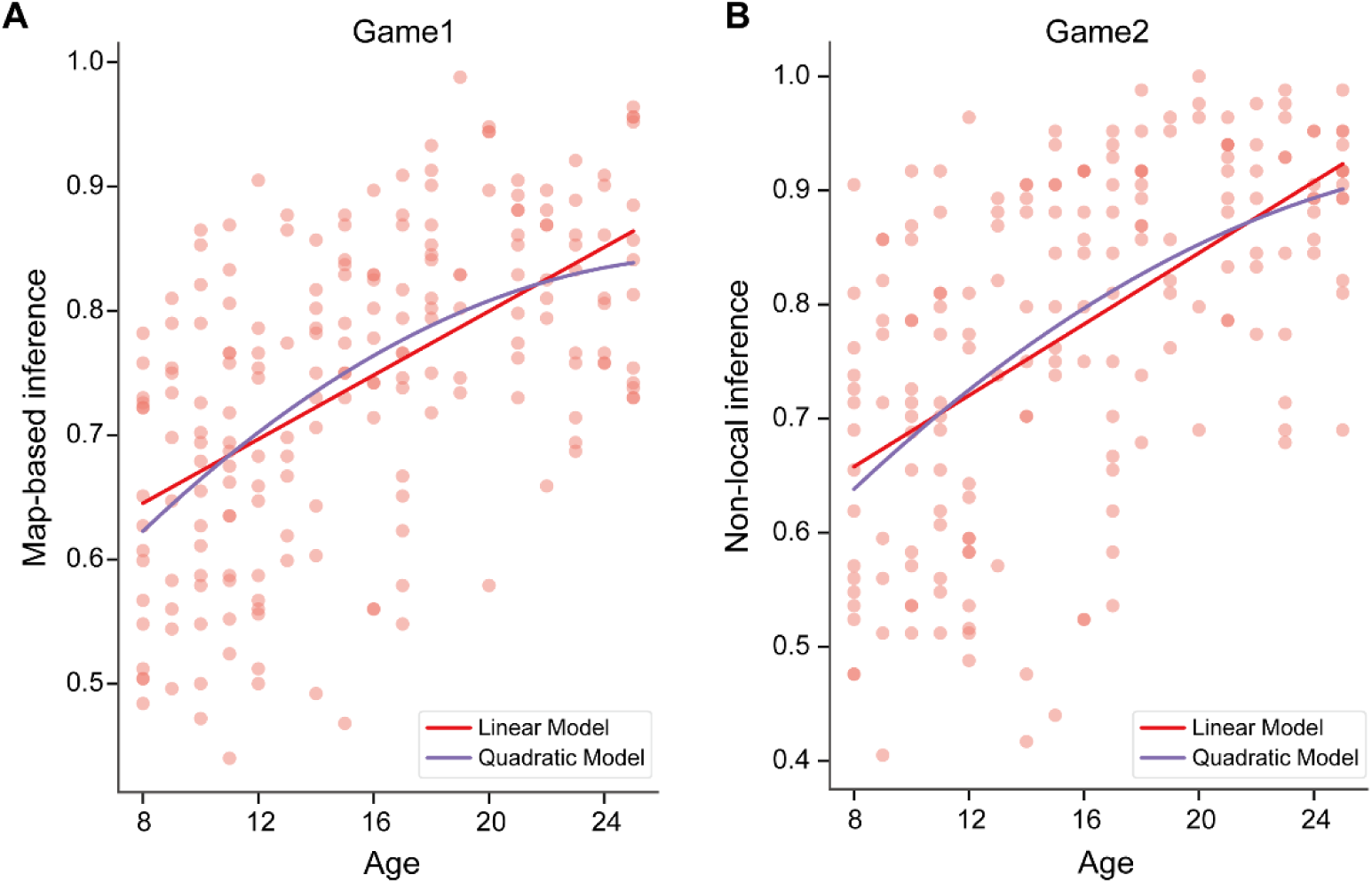
Comparison of linear and quadratic models for inference performance across age. **(A)** The fitting trajectories of age-related increase in map-based inference. This figure illustrates the relationship between age and map-based inference performance using both linear and quadratic models. The scatter plot represents individual inference accuracy scores. The red line depicts the linear fit, while the blue curve shows the quadratic relationship. **(B)** The layout mirrors panel **A**. Notably, the quadratic model fails to reach statistical significance as a predictor. Our analysis reveals that the linear model (red line) is a better fit to the data compared to the quadratic model (blue curve).

**Data S1.11.**
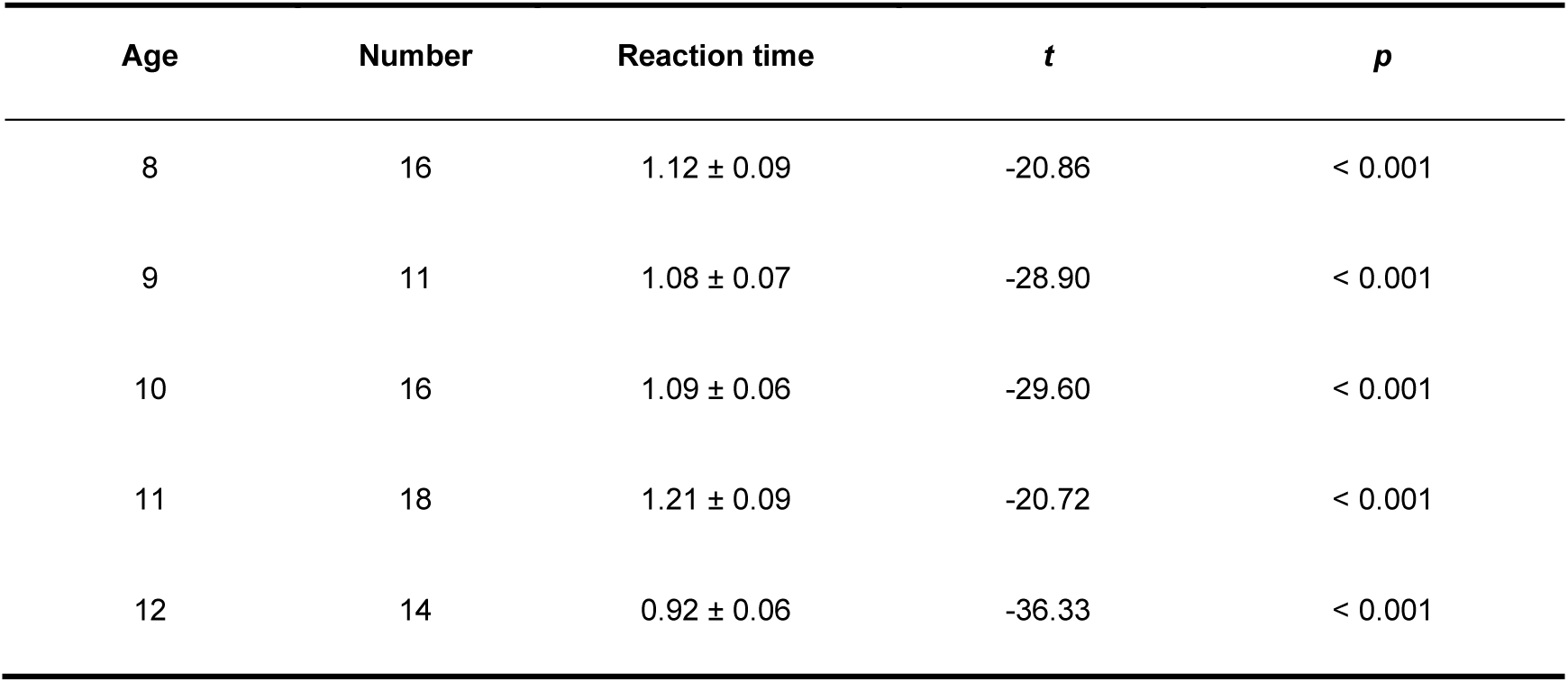
Reaction time of children aged 8-12 years. Mean reaction times (in seconds) with standard errors are shown for each age group. The number of participants per age group is indicated. Two tailed one-sample t-tests were conducted to compare each group’s mean reaction time against the maximum allowed time of 3 seconds. All groups demonstrated significantly faster reaction times than the 3-second threshold (*p* < 0.001 for all comparisons).

| Age | Number | Reaction time | *t* | *p* |
| --- | --- | --- | --- | --- |
| 8 | 16 | 1.12 ± 0.09 | -20.86 | < 0.001 |
| 9 | 11 | 1.08 ± 0.07 | -28.90 | < 0.001 |
| 10 | 16 | 1.09 ± 0.06 | -29.60 | < 0.001 |
| 11 | 18 | 1.21 ± 0.09 | -20.72 | < 0.001 |
| 12 | 14 | 0.92 ± 0.06 | -36.33 | < 0.001 |

**Data S2.**
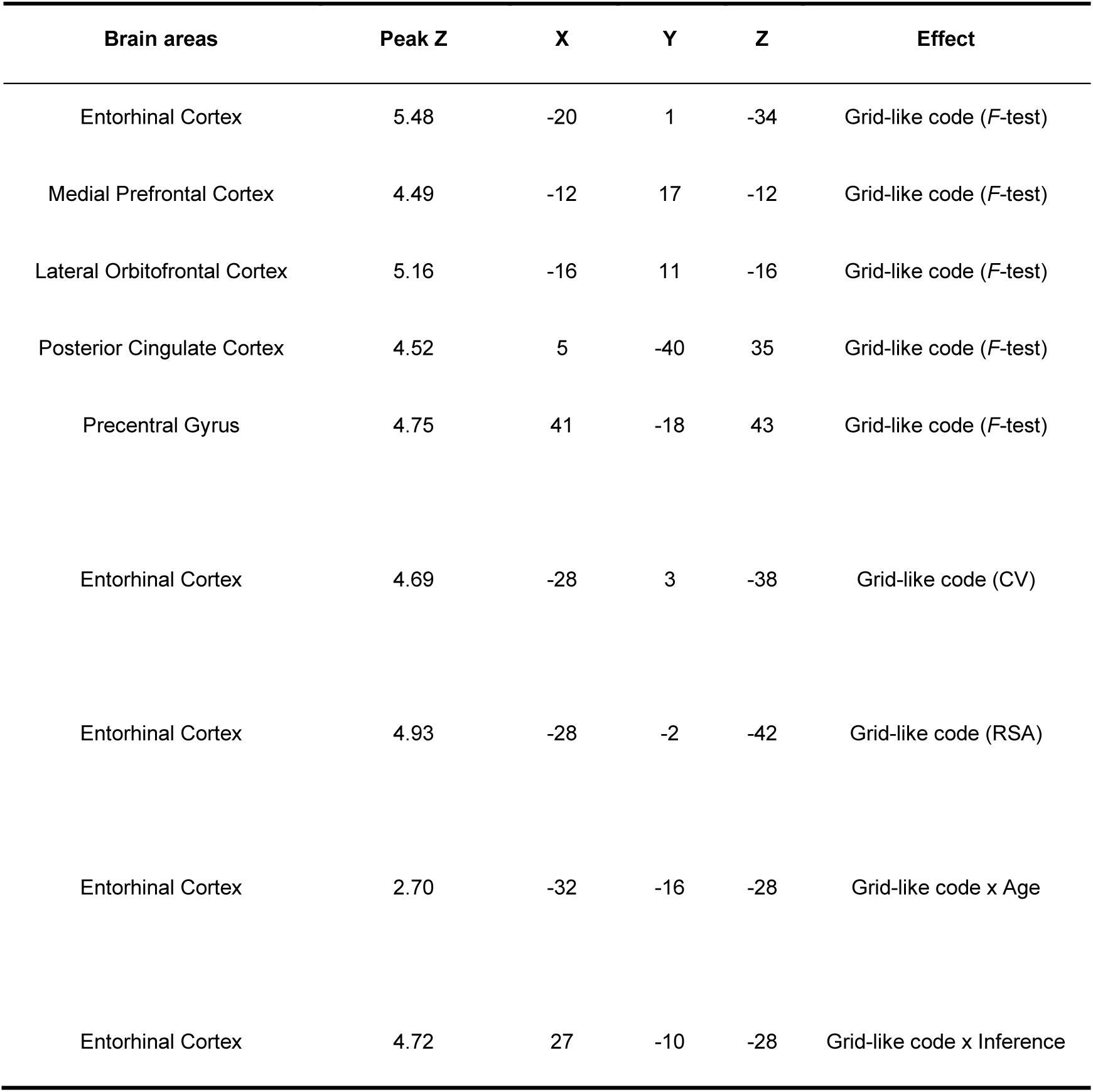
Grid-like code in Game1. Whole brain analysis shows the effect related to grid-like code in Game1. The table provided information such as the Peak Z, which was the peak *z*-value of the corresponding cluster survived from the multiple comparison correction in group-level results. Additionally, the table showed the MNI coordinates of peak for the corresponding cluster, denoted as X, Y, and Z. The effect of “Grid-like code (*F*-test)” was associated with **Figure S2**, while the effects of “Grid-like code (CV)” and “Grid-like code (RSA)” were associated with **Figure 2B**. “CV” (cross-validation) measures the hexagonal consistency effect, and “RSA” (representational similarity analysis) is another independent, multivariate approach to measuring the grid-like code. Lastly, the “Grid-like code x Inference” and “Grid-like code x Age” denoted the whole-brain covariate analysis with age and inference performance.

| Brain areas | Peak Z | X | Y | Z | Effect |
| --- | --- | --- | --- | --- | --- |
| Entorhinal Cortex | 5.48 | -20 | 1 | -34 | Grid-like code (*F*-test) |
| Medial Prefrontal Cortex | 4.49 | -12 | 17 | -12 | Grid-like code (*F*-test) |
| Lateral Orbitofrontal Cortex | 5.16 | -16 | 11 | -16 | Grid-like code (*F*-test) |
| Posterior Cingulate Cortex | 4.52 | 5 | -40 | 35 | Grid-like code (*F*-test) |
| Precentral Gyrus | 4.75 | 41 | -18 | 43 | Grid-like code (*F*-test) |
| Entorhinal Cortex | 4.69 | -28 | 3 | -38 | Grid-like code (CV) |
| Entorhinal Cortex | 4.93 | -28 | -2 | -42 | Grid-like code (RSA) |
| Entorhinal Cortex | 2.70 | -32 | -16 | -28 | Grid-like code x Age |
| Entorhinal Cortex | 4.72 | 27 | -10 | -28 | Grid-like code x Inference |

### Data S3 Data quality in the entorhinal cortex

We optimised the echo-planar imaging (EPI) parameters to reduce susceptibility-induced BOLD sensitivity losses. This was achieved by specifically adjusting the slice tilt and the direction of phase-encoding. According to Weiskopf, et al. ^2^, this method can increase BOLD sensitivity by more than 60% and by at least 30% on a 3T scanner.

To ensure the reliability of our data, we conducted a signal-to-noise ratio (SNR) analysis focusing on the entorhinal cortex (EC), a brain region crucial for our study. Our results demonstrated that the temporal SNR (tSNR) in the EC did not correlate with participants’ age (Game1: *r* = −0.09, *p* = 0.19; Game2: *r* = −0.06, *p* = 0.38).

**Data S3.**
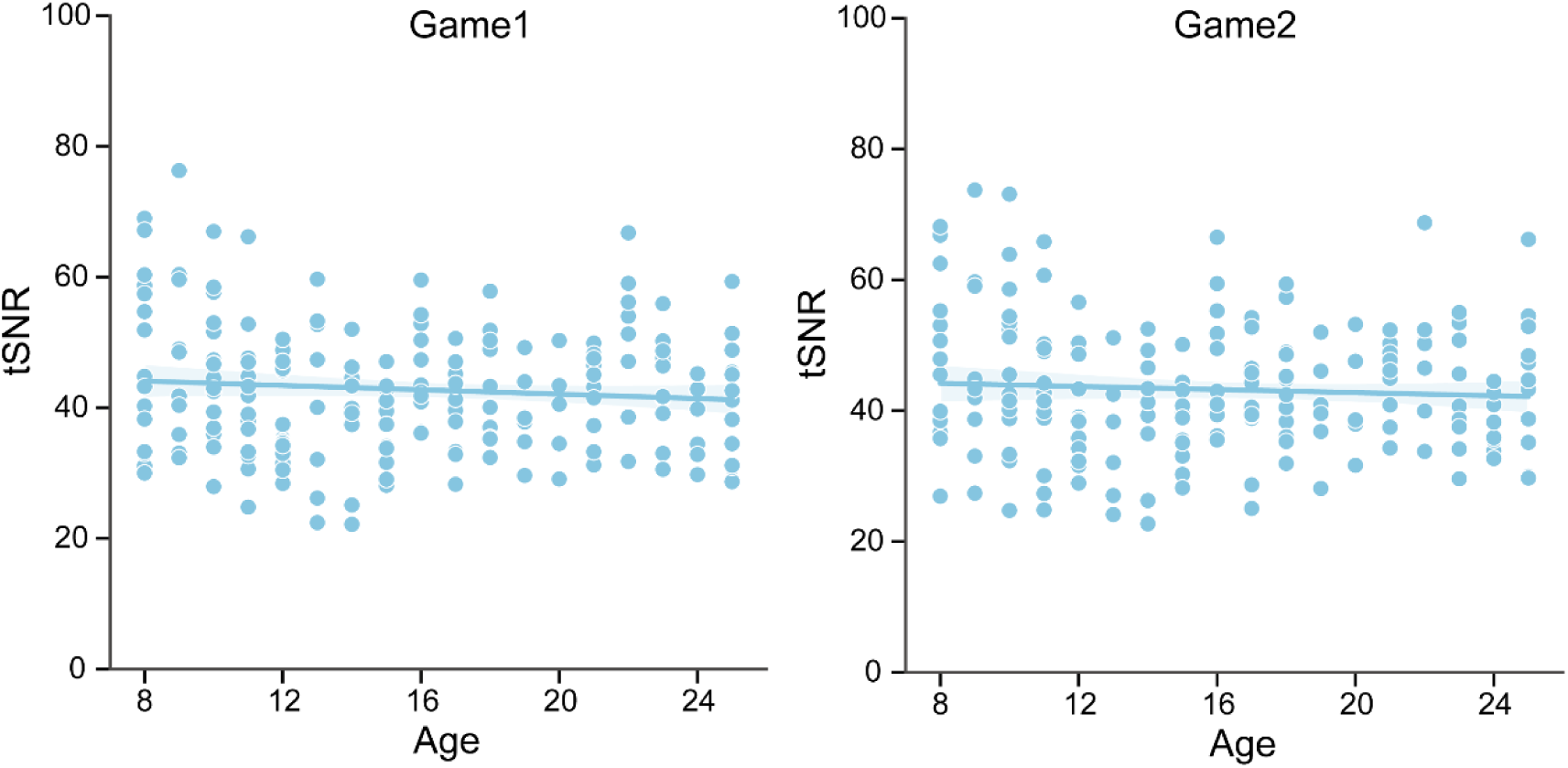
Temporal signal-to-noise ratio (tSNR) in the EC across age. Scatter plots show the relationship between tSNR and age in the EC for Game1 and Game2. Each point represents the tSNR value for an individual participant. No significant age-related effect was observed.

**Data S4.** Distance code in Game1. Whole brain analysis shows the effect related to distance code in Game1, associated with **Figure 3** and **Figure S3**.

| Brain areas | Peak Z | X | Y | Z | Effect |
| --- | --- | --- | --- | --- | --- |
| Medial Prefrontal Cortex | 6.23 | 7 | 47 | 23 | Distance code |
| Hippocampus | 4.15 | -32 | -14 | -12 | Distance code |
| Lateral Orbitofrontal Cortex | 4.99 | 41 | 33 | -12 | Distance code |
| Posterior Cingulate Cortex | 5.51 | 5 | -26 | 37 | Distance code |
| Middle Temporal Gyrus | 6.43 | 61 | -18 | -12 | Distance code |
| Angular Gyrus | 6.7 | 60 | -49 | 40 | Distance code |
| Medial Prefrontal Cortex | 4.3 | 5 | 57 | -2 | Distance code x Age |
| Lateral Orbitofrontal Cortex | 4.25 | 27 | 25 | -20 | Distance code x Age |
| Supramarginal Gyrus | 4.79 | -60 | -28 | 30 | Distance code x Age |
| Medial Prefrontal Cortex | 4.58 | -10 | 63 | -6 | Distance code x Inference |
| Lateral Orbitofrontal Cortex | 4.12 | 27 | 19 | -18 | Distance code x Inference |
| Angular Gyrus | 4.38 | 61 | -41 | 41 | Distance code x Inference |

### Data S5 Schematic of structured knowledge - schema vs. map

We illustrated the concept of structured knowledge—specifically, schema and map (**Data S5**). With development, subjects can form two types of representations: a general schema and a concrete map. The general schema is consistent across different environments and is characterised by the grid-like code. In contrast, the concrete map is environment-specific and is characterised by the distance code.

Our findings indicate that the grid-like schema representation in the EC supports the construction of map-like representations in the mPFC. We also observed significant age-related improvements in structural connectivity between the EC and mPFC, primarily through the cingulum bundle. The cingulum bundle serves as a crucial structural link in the default mode network, which includes mPFC, and PCC. These areas are key regions associated with the grid code in adults.

**Data S5.**
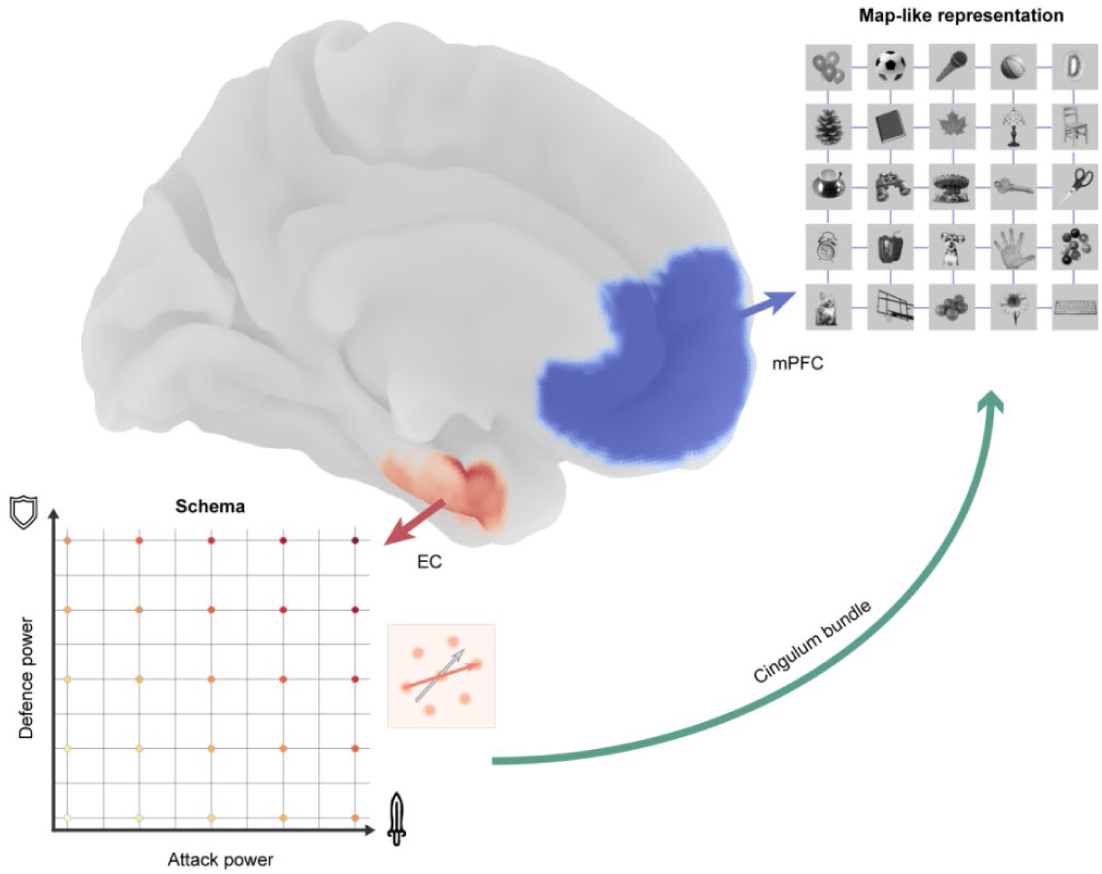
Schematic of structured knowledge in the brain. The brain organises knowledge using two distinct representations: schema and map. The EC, shown in orange, utilises a general schema characterised by a grid-like code that is consistent across environments. The mPFC, depicted in blue at the front of the brain, constructs concrete maps specific to the current environment, characterised by a distance code. The cingulum bundle, represented by a green curved arrow, connects the EC and mPFC. This white matter tract facilitates communication between the two regions, supporting the integration of schema and map representations. The cingulum bundle is part of the default mode network, which also includes the PCC, and plays a key role in advanced cognitive functions associated with the grid code in adults.

### Data S6 Inference growth with age beyond value computations

During decision time, stronger activations for correct trials were observed in the ventral striatum (peak *z* = 5.93, MNI coordinate [−17, 9, −12]), amygdala (peak *z* = 5.08, MNI coordinate [26, 0, −16]), and primary motor cortex (peak *z* = 5.52, MNI coordinate [7, −20, 57], whole-brain cluster-based FWE correction at *p* < 0.05, see also **Figure S4**), consistent with prior research on value-based decision-making^3,4^.

In our task, the key decision variable was the absolute value of AP-DP, with a larger value indicating an easier decision. Behaviourally, “|AP–DP|” predicted choices better than “AP-DP” (linear mixed model, *z* = 32.48, *p* < 0.001). Neurally, the mPFC encoded this absolute value code during decision time (peak *z* = 6.25, MNI coordinate [−8, 53, −4], see also **Data S7**). ROI analysis showed the mPFC represented the absolute value of *A*P− *D*P more than their direct difference (paired *t*-test *t*(202) = 2.36, *p* = 0.02, Cohen’s *d* = 0.23). Further ROI analysis showed that the value code in the mPFC, and the hippocampus correlated with both age and performance (**Figure S4F**), but no mediation effect was found (*β_ab_* = 0.03, *p* = 0.09, CI [−0.002, 0.066]), (*β_ab_* = 0.01, *p* = 0.56, CI [−0.02, 0.04]).

**Data S7.** Value code at decision time. Whole brain analysis shows the value code at decision. The effect of “Value code” was associated with **Figure S4.**

| Brain areas | Peak Z | X | Y | Z | Effect |
| --- | --- | --- | --- | --- | --- |
| Entorhinal Cortex | 6.06 | -16 | -6 | -18 | Value code |
| Medial Prefrontal Cortex | 6.25 | -8 | 53 | -4 | Value code |
| Hippocampus | 7.18 | -22 | -10 | -16 | Value code |
| Lateral Orbitofrontal Cortex | 6.36 | 25 | 7 | -18 | Value code |
| Posterior Cingulate Cortex | 6.32 | 5 | -32 | 47 | Value code |
| Precentral Gyrus | 7.26 | -12 | -30 | 45 | Value code |
| Supramarginal Gyrus | 8.41 | 61 | -28 | 31 | Value code |
| Frontal Pole | 6.01 | -31 | 41 | 26 | Value code |
| Medial Prefrontal Cortex | 5.2 | -8 | 51 | -8 | Value code x Age |
| Hippocampus | 4.74 | -26 | -14 | -14 | Value code x Age |
| Anterior Cingulate Cortex | 5.13 | -2 | -14 | 41 | Value code x Age |
| Superior Temporal Gyrus | 5.77 | -50 | -17 | 0 | Value code x Age |
| Medial Prefrontal Cortex | 5.66 | -6 | 25 | -8 | Value x Inference |
| Hippocampus | 3.85 | -26 | -10 | -16 | Value x Inference |
| Lateral Orbitofrontal Cortex | 4.16 | -20 | 35 | -12 | Value x Inference |
| Posterior Cingulate Cortex | 5.49 | -2 | -15 | 42 | Value x Inference |
| Temporal Pole | 5.99 | 58 | 5 | -1 | Value x Inference |

### Data S8 Independent effects between distance and value computation

We also tested whether the map-like neural representations observed during mental navigation could be confounded by value-based decision processes (e.g., value code and decision difficulty).

First, our experimental design temporally separated mental navigation from decision-making by introducing a jittered delay between the presentation of Monster-2 (M2) and the decision rule (choosing the attacker and defender), enabling us to dissociate map-like representations from value-related signals. Behaviourally, we examined the correlation between distance code, 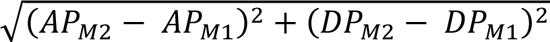, and value code, |*A*P − *D*P| for each object pair during inference. The analysis revealed a negligible correlation coefficient (*r* = −0.001 ± 0.004) across subject, which, after Fisher’s z-transformation, did not differ from zero (*t*(202) = −0.32, *p* = 0.75; see **Data S8**), indicating distance and value effects are independently modulated.

To confirm this separation neurally, we implemented a GLM with both distance and value regressors at two key time points: M2 presentation (mental navigation) and the decision-making phase. Distance was always modelled as the Euclidean distance, and value at decision time was modelled as |*A*P − *D*P| at decision time. At the M2 presentation stage—when the rule was unknown—value was defined by the mean of two potential decision values, |*A*P_*M*1_ − *D*P_*M*2_| and |*A*P_*M*2_ − *D*P_*M*1_|. Whole-brain analysis identified significant activation in the mPFC related to the distance code (**Data S8B;** whole-brain cluster-based FWE correction at *p* < 0.05, peak z = 6.24, MNI coordinate [9, 48, 25]). Follow-up ROI analysis revealed that this mPFC distance code was positively correlated with both age (*r* = 0.25, *p* < 0.001) and inference performance (*r* = 0.24, *p* = 0.001). Notably, we observed a significant interaction between task phase and neural representation in the mPFC (*F*(1,808) = 4.29, *p* = 0.04; see also **Figure S4H**). During mental navigation, the mPFC predominantly encoded Euclidean distance (0.26 ± 0.09) rather than value (0.14 ± 0.02), but shifted to value coding at decision-making (distance: 0.17 ± 0.08; value: 0.37 ± 0.08). These findings confirm that value coding does not confound our distance-related results.

We next considered whether the distance effect instead reflects decision difficulty. A linear mixed-effects logistic regression, modelling trial-by-trial accuracy with distance as the predictor (and including random intercepts and slopes for participants), yielded no significant relationship between distance and accuracy (*β* = 0.004, *p* = 0.11, CI [−0.001, 0.009]). This finding aligns with our design, whereby decision difficulty often varied even for the same distance. For example, comparing object A (AP: 4, DP: 5) with B (AP: 2, DP: 3) is straightforward because one option is superior on both dimensions, whereas comparing object C (AP: 4, DP: 3) with D (AP: 2, DP: 5) is more challenging, despite both pairs sharing the same Euclidean distance. Thus, distance was effectively decoupled from decision difficulty. Consistent with this, no association was found between the decision variable (|AP–DP|) and the distance code.

Finally, we checked whether brain signals might reflect decision difficulty by contrasting smaller (≤3) and larger (>3) Euclidean distances. Under a repetition-suppression perspective^5^, more overlapping representations should produce stronger suppression for smaller distances. Indeed, neural responses for objects separated by smaller distances showed significantly stronger suppression (paired t-test on mPFC activity: *t*(202) = −3.81, *p* < 0.001, Cohen’s *d* = −0.14), indicating a map-like similarity effect rather than an index of decision ease. Collectively, these observations rule out value coding and decision difficulty as confounding factors and reinforce that the mPFC distance code reflects map-like representations.

**Data S8.**
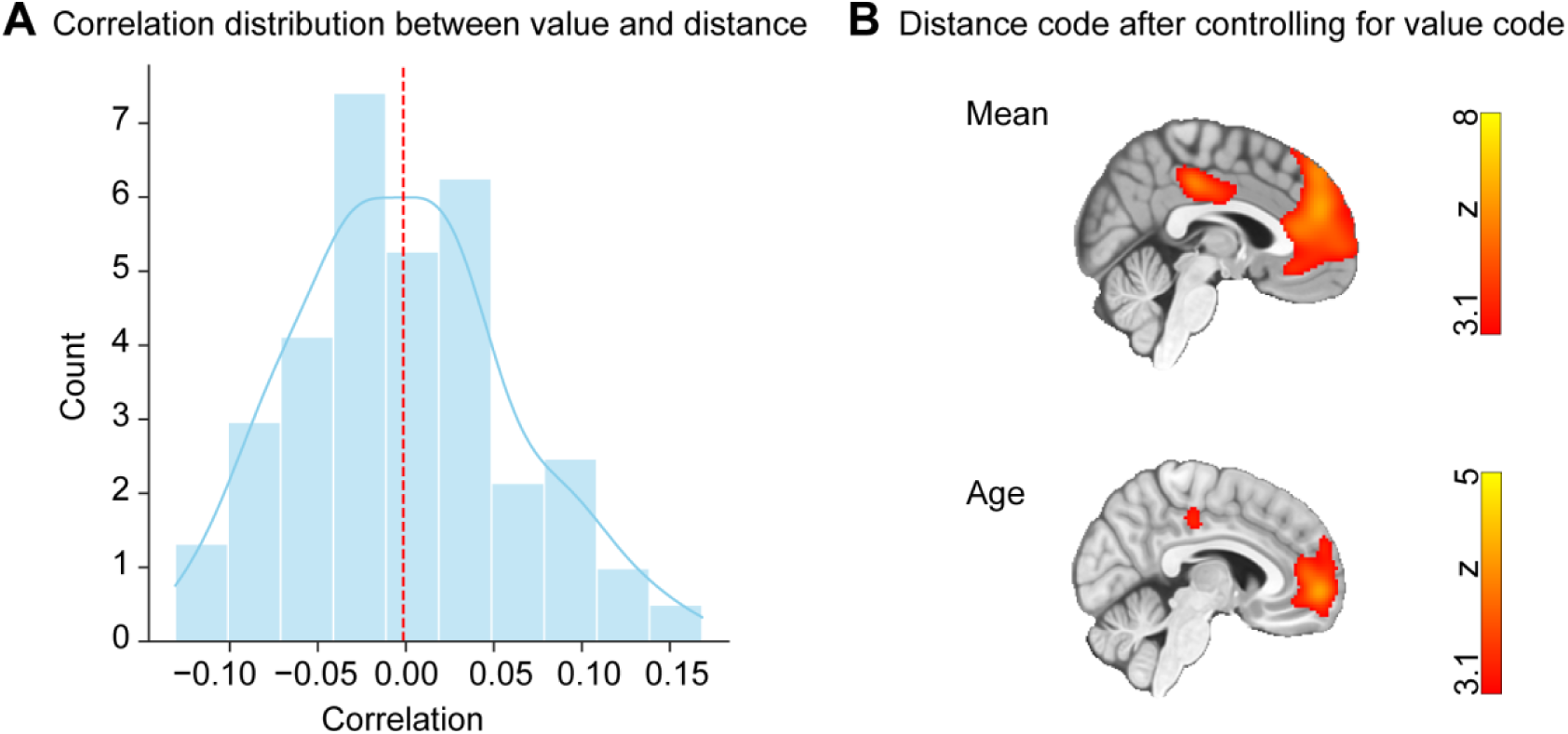
Independent effects between distance and value computation. **(A)** The histogram shows how each participant’s Euclidean distance–value correlation is distributed. Each bar represents the number of participants whose correlation fell within a given range. The mean correlation coefficient is –0.001, indicating that Euclidean distance and value are independently modulated across subjects. **(B)** After controlling for value effects, a whole-brain analysis still revealed significant mPFC activation encoding Euclidean distance (top panel; cluster-based FWE correction at *p* < 0.05). An age-related increase in the mPFC distance code was also observed (bottom panel). These results confirm that distance coding in the mPFC is not by value-based decision signals.

### Data S9 Schema and map in Game1 and Game2

We illustrated the task structures of both games in **Data S9**.

In the piecemeal learning session, participants learned pairwise relationships by comparing objects with their neighbours along one dimension. The distance between neighbouring objects was defined as a one-rank difference, establishing the task structure for Game 1.

In the local learning session, new objects were introduced. Participants acquired new relationships using the same comparison method as before. These new objects were positioned around the centre of the map, and their relationships with existing objects were updated. Participants were informed that the relationships between existing objects would remain unchanged, allowing them to assimilate the new knowledge into a consistent space. This formed the task structure for Game 2.

**Data S9.**
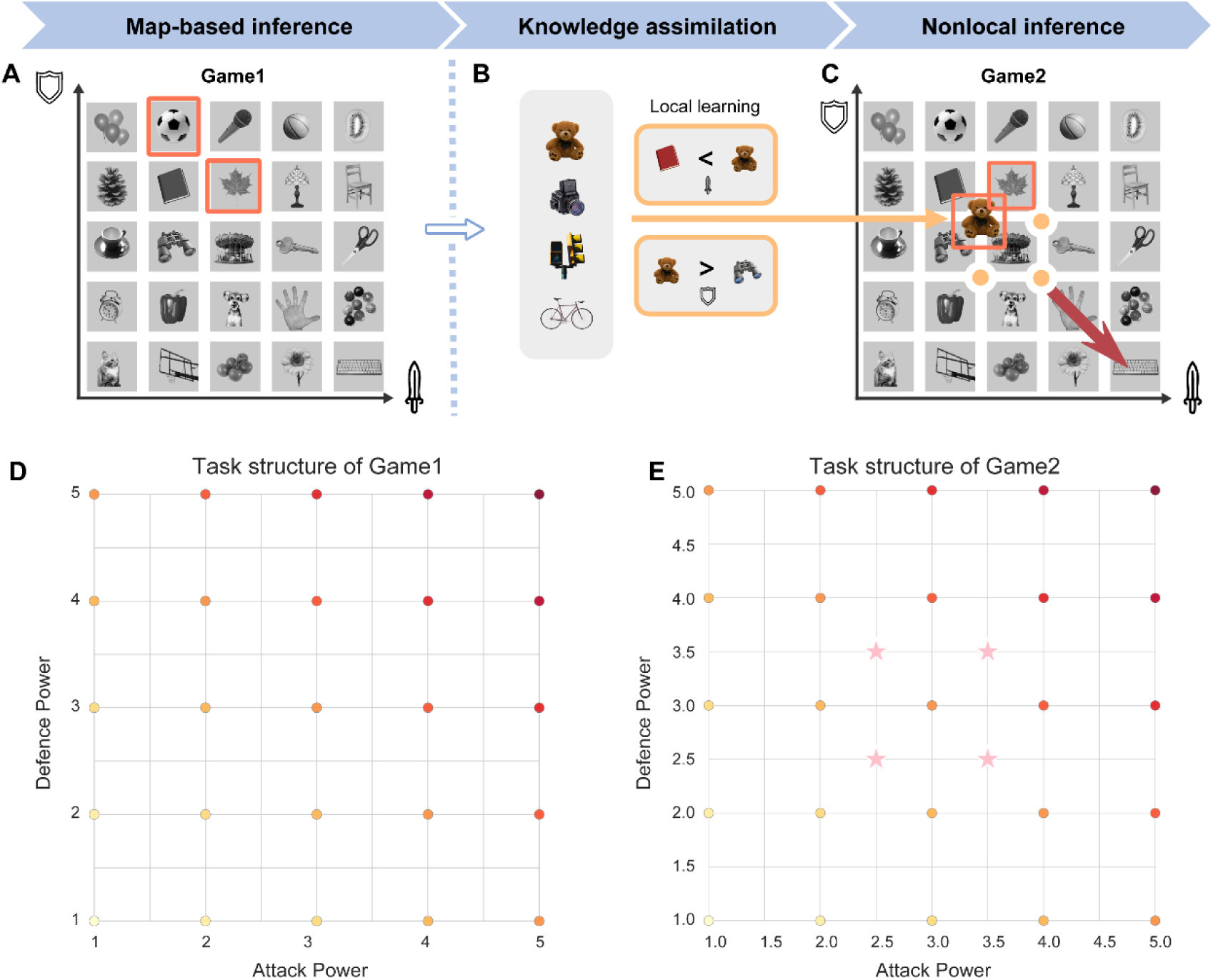
Schema and map in Game1 and Game2. **(A)** The 2D knowledge map of Game 1. Red rectangles illustrate two adjacent characters (football and maple leaf) with a one-rank difference in both dimensions. This is an example of a pairwise relationship in learning. This relationship persists in Game 2. **(B)** Local learning process for assimilating new information. Participants learnt about four new objects by comparing them to adjacent existing objects in each dimension. This comparison process helps participants recognise the distinct attributes of new objects. **(C)** Updated map incorporating new characters (e.g., bear). New objects are positioned around the centre of the map, maintaining a half-rank distance from adjacent existing objects. **(D)** Task structure of Game 1. Each point represents an object’s position in this space, defined by its rank in attack power and defence power. Existing objects are colour-coded from yellow to red. **(E)** Task structure of Game 2. Each point represents an object’s position in this space. Existing objects are colour-coded from yellow to red. New objects are represented by larger, pink stars.

### Data S10 Consistent neural activities in Game1 and Game2

We observed consistent neural activities in both Game1 and Game2. Specifically, in Game1, at the onset of M1, whole-brain analysis showed significant activity in the visual cortex, posterior cingulate cortex, and posterior hippocampus (**Figure S5A**). Moving to the onset of M2, there were notable activations in the anterior cingulate cortex, visual cortex, and superior parietal lobe (**Figure S5B**). At the decision-making stage, significant activation was observed in areas including the cerebellum, visual cortex, striatum, primary motor cortex, and dorsolateral prefrontal cortex (**Figure S5C**).

Similarly, in Game2, the activation patterns across each phase closely resembled those in Game1, indicating similar neural involvement in both tasks. This is depicted in the **Figure S5D-F**. All these brain maps were subjected to whole-brain family-wise error (FWE) correction at the cluster level (*p* < 0.05), with a cluster-forming voxel threshold of *p* < 0.001.

**Data S11.** The effect related to grid-like code in Game2. Whole brain analysis shows the effect related to grid-like code in Game2. The effect of “Grid-like code (*F*-test)” was associated with **Figure S6**, while the covariate results of “Map-alignment x Inference” were associated with **Figure 4C**, no significant covariate results with age were found.

| Brain areas | Peak Z | X | Y | Z | Effect |
| --- | --- | --- | --- | --- | --- |
| Entorhinal Cortex | 3.75 | 19 | -10 | -32 | Grid-like code (*F*-test) |
| Medial Prefrontal Cortex | 4.1 | -4 | 21 | -6 | Grid-like code (*F*-test) |
| Lateral Orbitofrontal Cortex | 4.31 | 15 | 21 | -26 | Grid-like code (*F*-test) |
| Posterior Cingulate Cortex | 3.9 | 1 | -44 | 27 | Grid-like code (*F*-test) |
| Supplementary Motor Cortex | 4.5 | -4 | 5 | 60 | Grid-like code (*F*-test) |
| Cuneal Cortex | 3.87 | 6 | -86 | 25 | Grid-like code (*F*-test) |
| Entorhinal Cortex | 4.32 | 29 | 0 | -42 | Map-alignment x Inference |

**Data S12.** Distance code in Game2. Whole brain analysis shows the effect related to distance code in Game2, associated with **Figure 5** and **Figure S6**.

| Brain areas | Peak Z | X | Y | Z | Effect |
| --- | --- | --- | --- | --- | --- |
| Medial Prefrontal Cortex | 7.13 | -8 | 55 | 3 | Distance code |
| Hippocampus | 5.76 | 29 | -38 | -10 | Distance code |
| Lateral Orbitofrontal Cortex | 6.77 | 23 | 13 | -16 | Distance code |
| Posterior Cingulate Cortex | 6.96 | -5 | -19 | 43 | Distance code |
| Supramarginal Gyrus | 7.3 | 58 | -37 | 36 | Distance code |
| Middle Temporal Gyrus | 6.37 | 60 | -13 | -13 | Distance code |
| Medial Prefrontal Cortex | 6.41 | -4 | 39 | 9 | Distance code x Age |
| Hippocampus | 4.50 | -24 | -10 | -16 | Distance code x Age |
| Lateral Orbitofrontal Cortex | 5.18 | -34 | 13 | -18 | Distance code x Age |
| Posterior Cingulate Cortex | 5.18 | 1 | -22 | 41 | Distance code x Age |
| Middle Temporal Gyrus | 5.83 | 55 | -19 | -6 | Distance code x Age |
| Supramarginal Gyrus | 5.10 | 55 | -36 | 31 | Distance code x Age |
| Medial Prefrontal Cortex | 7.69 | 3 | 33 | -12 | Distance code x Inference |
| Hippocampus | 5.82 | -26 | -12 | -14 | Distance code x Inference |
| Lateral Orbitofrontal Cortex | 6.37 | -32 | 17 | -22 | Distance code x Inference |
| Posterior Cingulate Cortex | 6.57 | 7 | -20 | 37 | Distance code x Inference |
| Supramarginal Gyrus | 6.42 | 57 | -33 | 32 | Distance code x Inference |
| Superior Temporal Gyrus | 5.89 | 55 | -3 | -12 | Distance code x Inference |

### Data S13 Reproducible results across tasks, modalities and different parcellations

Our study consistently demonstrated reproducible effects across different tasks and analyses. For the cognitive map representation, we observed a grid code effect in the EC using three independent methods (**Figure 2** and **Figure S2**), and this effect was also replicated in Game 2 (**Figure 4**). The distance code effect in the mPFC was consistently observed across both tasks (**Figure 3** and **Figure 5**). We further demonstrated consistent developmental coupling between the structural maturation of the EC and mPFC, which supports their functional interaction and facilitates inference (**Figure 6**). These neural representations of the cognitive map also underpin the well-established relationships between age and IQ (**Figure 6**). These converging results across tasks and modalities strengthen the robustness of our findings.

We also observed reproducible structural results across different parcellation methods, confirming that our developmental trends are genuine and not due to artefacts related to atlas selection. Both the Desikan-Killiany (DK) atlas and the masks used in our functional analyses exhibited similar developmental patterns in the EC and mPFC. We conducted a ROI analysis of total grey matter volume in the bilateral EC and mPFC using the DK parcellation (**Data S13A**), finding an age-related increase in EC volume that correlated with improved inference performance, measured as mean accuracy across Game 1 and Game 2. We also showed a similar ROI analysis using the EC and mPFC masks as used in our functional analyses (**Data S13B**), which revealed comparable developmental patterns and reinforced the reliability of our findings. Overall, these findings indicate our results are robust and consistently replicated across multiple methods and datasets.

**Data S13.**
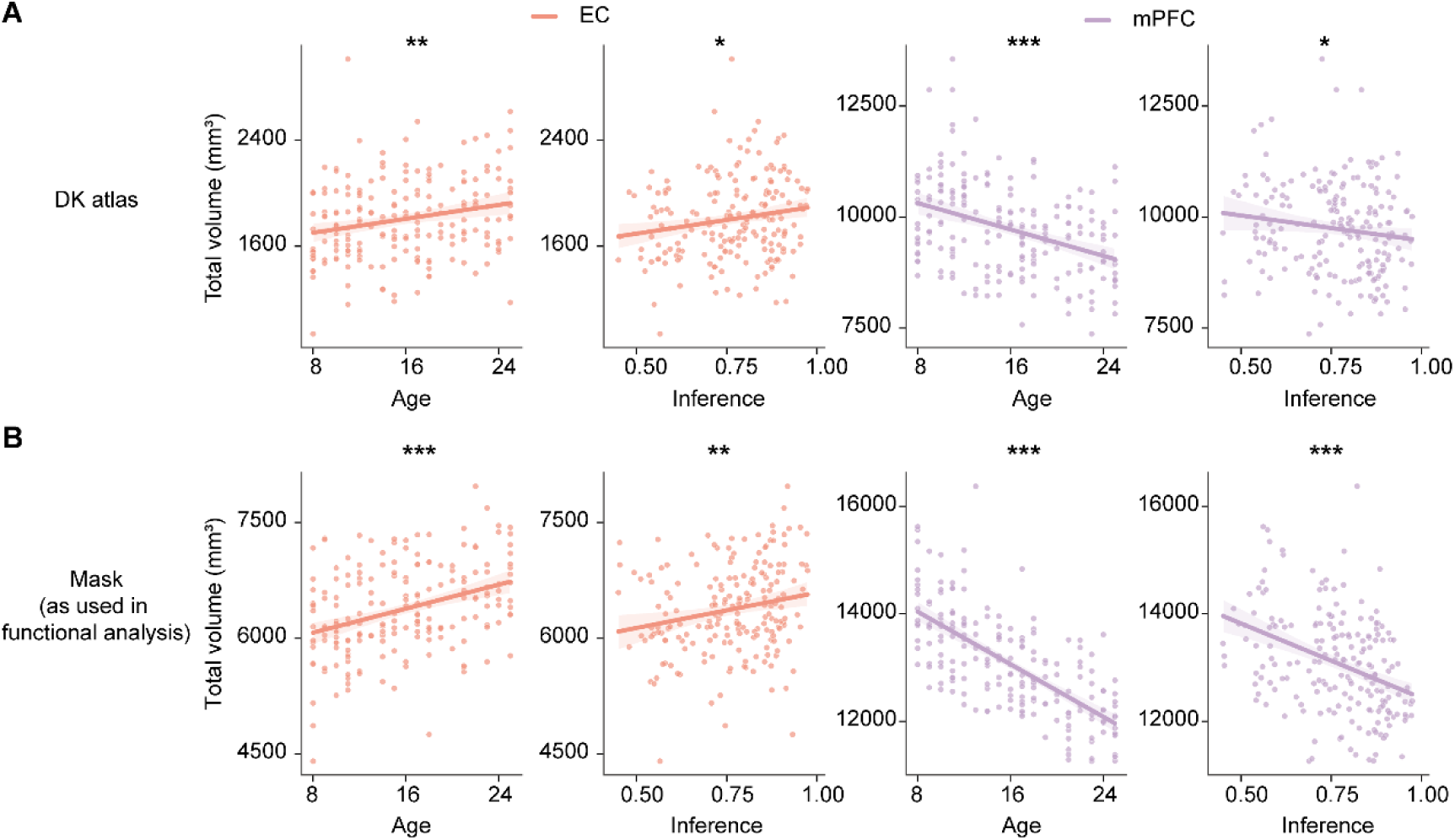
Consistency of developmental patterns across different parcellation methods. **(A)** ROI analysis of the grey matter volume in the bilateral EC and mPFC using the Desikan-Killiany parcellation. We observed an age-related increase in EC volume, which also correlated with improved inference performance (measured as the mean accuracy across Game 1 and Game 2). **(B)** ROI analysis using the EC and mPFC masks as used in our functional analyses. The results show similar developmental patterns to those in panel **A**.

### Data S14. Training sessions for each age group and learning phase

We provide a detailed breakdown of training sessions. As also illustrated in **Figure S1**, a clear developmental trend emerges in how many sessions participants require. Younger children (aged 8–12) needed significantly more sessions (20.05 ± 1.49, N = 75) compared with adolescents (13–17; 9.11 ± 0.37, N = 55) and adults (18–25; 8.00 ± 0.29, N = 73). This pattern was consistent across training stages:

#### Single-dimensional training

Children typically required three to four sessions (3.24 ± 0.34) to reach the ≥80% criterion for each dimension, whereas older participants usually completed this stage in one to two sessions (adolescents: 1.62 ± 0.15; adults: 1.15 ± 0.04).

#### Intermixed training

The most pronounced age differences appeared during the intermixed phase. Children aged 8–12 needed nine to sixteen sessions (10.93 ± 1.11), while adolescents and adults both completed it in roughly three to four sessions (adolescent: 3.91 ± 0.32; adult: 3.73 ± 0.27). A significant age × phase interaction (*F*(6,800) = 16.73, *p* < 0.001) and simple effects analyses (Dim-1: *F*(2, 200) = 22.45, *p* < 0.001; Dim-2: *F*(2, 200) = 22.41, *p* < 0.001; intermixed: *F*(2, 200) = 31.92, *p* < 0.001; review: *F*(2, 200) = 6.47, *p* = 0.002; see also **Figure S1**) indicate that children particularly struggled to separate and integrate these two dimensions into a coherent 2D knowledge structure.

#### Review

In the final review stage, all age groups spent a comparable number of practice sessions: 2.22 ± 0.09 on average. Children reviewed 2.64 ± 0.22 sessions, while adolescents and adults reviewed 1.96 ± 0.11 and 1.97 ± 0.06 sessions, respectively, before starting the formal experiment.

**Data S14.** Training sessions for each age group and learning phase. Mean ± SEM sessions spent in each learning phase (Single-Dimensional, Intermixed, and Review), as well as total sessions (All phases), are listed for children (N = 75), adolescents (N = 55), and adults (N = 73). In the Single-Dim phase, we report the mean number of sessions across AP and DP dimensions. Children required significantly more training time across all phases compared to adolescents and adults.

| Age group | Single-Dim | Intermixed | Review | All phase | N |
| --- | --- | --- | --- | --- | --- |
| Children | 3.24 ± 0.34 | 10.93 ± 1.11 | 2.64 ± 0.22 | 20.05 ± 1.49 | 75 |
| Adolescents | 1.62 ± 0.15 | 3.91 ± 0.32 | 1.96 ± 0.11 | 9.11 ± 0.37 | 55 |
| Adults | 1.15 ± 0.04 | 3.73 ± 0.27 | 1.97 ± 0.06 | 8.00 ± 0.29 | 73 |
| All participants | 2.05 ± 0.16 | 6.44 ± 0.49 | 2.22 ± 0.09 | 12.75 ± 0.69 | 203 |

